# Phenotype diversity and extinction dynamics of the European Narrow-Headed Vole, *Stenocranius anglicus* (Hinton, 1910) (Arvicolinae, Cricetidae, Rodentia), in Central Europe

**DOI:** 10.64898/2026.02.09.704770

**Authors:** Nikoleta Dubjelová, Tereza Hadravová, Martin Ivanov, Ivan Horáček

## Abstract

The European Pleistocene populations of the narrow-headed vole (*Stenocranius gregalis)*, an index species of the Palearctic glacial communities, were recently found to differ from the extant Asian species by a deep genetic divergence and are to be considered a separate species, *Stenocranius anglicus,* which had to persist through the interglacial stages in local European refugia. Here, we analyze over 2000 first lower molars from 14 stratified localities in the Czech Republic and Slovakia, spanning the Middle Pleistocene to Holocene, employing geometric morphometrics, biometric measurements, and morphotype classifications to assess molar shape variation.

Our results demonstrate persistent morphological variability, with particularly high morphotype diversity during MIS 5–3, followed by simplification and reduced variance in post–LGM populations. Morphological divergence was greater among geographic localities than stratigraphic stages, suggesting strong regional and ecological influences. Stratified sequences reveal diverse evolutionary trajectories from long-term morphological stability in refugia to gradual simplification preceding extinction in the early Holocene. These patterns align with broader Eurasian trends but also highlight regionally specific responses to climatic and ecological change accompanying the species’ extinction dynamics during the early to middle Holocene.

The paper underscores the importance of integrating detailed morphometrics with stratigraphic and ecological evidence to shed light on these topics.

## Introduction

The narrow-headed vole is considered, ever since the works of Nehring (1875, 1890), Newton (1894), Woldřich (1897) and Hinton (1910), a robust indicator of glacial environments (Nadachowski 1982; Horáček and Sanchéz-Marco 1984; Horáček and Ložek 1988; Baca et al. 2019). Detailed information on the species abundance and dominance throughout the Northern Eurasian Middle and Late Pleistocene localities can be found in numerous publications (Sutcliffe and Kowalski 1976; Horáček and Sanchéz-Marco 1984; Pazonyi 2004; Bogićević et al. 2012; López-García et al. 2015, 2017, 2019; Laplana et al. 2016; Németh et al. 2017; Luzi 2018; Popov 2018; Magyari et al. 2022) detailing descriptions of the species’ biometric changes and morphological variability. The extant range of *Stenocranius gregalis* (Pallas, 1779) sensu stricto covers the tundra zone of northern Asia, the steppe zones and mountains of Central Asia and southern Siberia and extends up to north China (GBIF.org); Wilson and Reeder 2005; Shi et al. 2021). In central and western Europe, it represented a core element of glacial communities throughout the Middle and Late

Pleistocene, but it has never been found in any clearly interglacial contexts. This has been long kept as the most pronounced example of the range dynamics, characterized by alternation of large-scale expansions with mass local extinction events during each glacial cycle, providing a robust support to the central paradigm of European historical biogeography which predicts the large scale migrations as the major agent of the radical rearrangements of community structure along the Quaternary climatic cycles: glacial retreat of interglacial elements into the Mediterranean refugia synchronous with by range expansions of the glacial communities elements expanding from their interglacial refugia in tundra and steppe zones of Eastern Europe. This concept, which arose at the turn of the 19^th^ and 20^th^ centuries (Nehring 1890), was supported by many studies throughout the 20^th^ century and became a default hypothesis for molecular phylogeography of European taxa (Hewitt 2004). Stratigraphical distribution of lemmings or *Microtus gregalis* (or *Stenocranius gregalis* in a current generic assignment - see Kryštufek and Shenbrot 2022) was taken as a robust example of its validity. Yet, the analyses of ancient DNA by Baca et al. (2019) demonstrated that all the Late Pleistocene European samples of narrow-headed vole form a single clade which distinctly differs from extant populations of *M. gregalis* being separated for more than 200–300 ka B.P. Consequently, the European form is to be looked upon as a separate species (for which the prior name *Microtus anglicus* Hinton, 1910 is available) distributed continuously in Central and Western Europe at least during the last two or three glacial cycles. Therefore, contrary to traditional paleobiogeographical hypotheses (comp. e.g. Shi et al. 2021), the Late Pleistocene appearance of the clade in Europe did not arise of the westward expansion of *Stenocranius gregalis*, and, at the same time, the European narrow-headed vole, *Stenocranius anglicus* (Hinton, 1910), did not retreat from Europe during interglacial periods but survived the unfavorable conditions in European interglacial refugia, from which it then recolonized Europe during glacial stages.

Anyhow, that species is missing from the extant mammal fauna of Europe, knowledge of which is based on an extraordinarily robust record of field data, with numerous detailed molecular genetic analyses in all covered species (Mitchell-Jones et al. 2026). Thus, it seems for granted that the European narrow-headed vole went extinct in Europe sometime between today and the end of the last glaciation (Pazonyi 2004; Markova et al. 2009; Stuart 2015) and that it disappeared from Central Europe by the mid–Holocene (∼5000 years ago) (Sommer and Nadachowski 2006). The specifics of how and where this extinction unfolded remain, sadly, unclear. Unlike the extinction dynamics observed in large mammals, such as mammoths, the European narrow-headed vole’s story allows us to examine the course and causes of species extinction apart from human influence.

In any case, the novel view on the taxonomic status of the European *Microtus gregalis* (sensu lato) clade calls for more detailed information on patterns of phenotypic variation and abundance dynamics of that form. The still incomplete database of Czech and Slovak small ground mammals of the Late Pleistocene and Holocene age covers 775 community samples of 102 sites with MNI 24100 individuals. *Stenocranius anglicus*, with MNI 6352 recorded in 235 community samples from 68 sites (26,4 % of all clades MNI), is the most common element there. The vast majority of the database records come from continuous sedimentary sequences, providing a possibility to trace variation dynamics and extinction process in detail (comp. e.g., Horáček and Sanchéz-Marco 1984; Ložek and Horáček 1984, 2006, 2007; Ložek et al. 1987, 1989; Horáček and Ložek 1988, 1990, 1993; Horáček et al. 2002; Horáček and Jahelková 2005; Horáček 2007).

This study investigates morphological variability in the first lower molar (m1) of *Stenocranius anglicus* across 14 stratigraphically and geographically diverse localities in the Czech Republic and Slovakia. The objectives are to: (i) characterize the extent of m1 phenotypic variation and patterns of between-site differences; (ii) compare the effects of geographic versus stratigraphic drivers of phenotypic variation; (iii) identify trends in phenotypic rearrangements across glacial cycles and the Last Glacial Maximum (LGM); and (iv) shed light on phenotype and community changes associated with the extinction dynamics of the species. Using 2D geometric morphometrics and traditional morphometric methods, we examined patterns of phenotypic variation throughout the last glaciation and early Holocene in model sites representing diverse regions of the Czech Republic and Slovakia and undertook detailed between-populations comparisons.

## Material and methods

### The material and sites

A total of 2081 first lower molars (m1, either left or right, supposedly of adult individuals) of *Stenocranius anglicus* were analyzed from 48 community samples of 14 fossil localities in the Czech Republic and Slovakia, spanning from MIS 12/14 to the early Holocene (Fig. 1)- for further details see Supplementary file S1. All specimens are deposited in curated collections of the Department of Zoology, Charles University, Prague.

**Figure 1.**
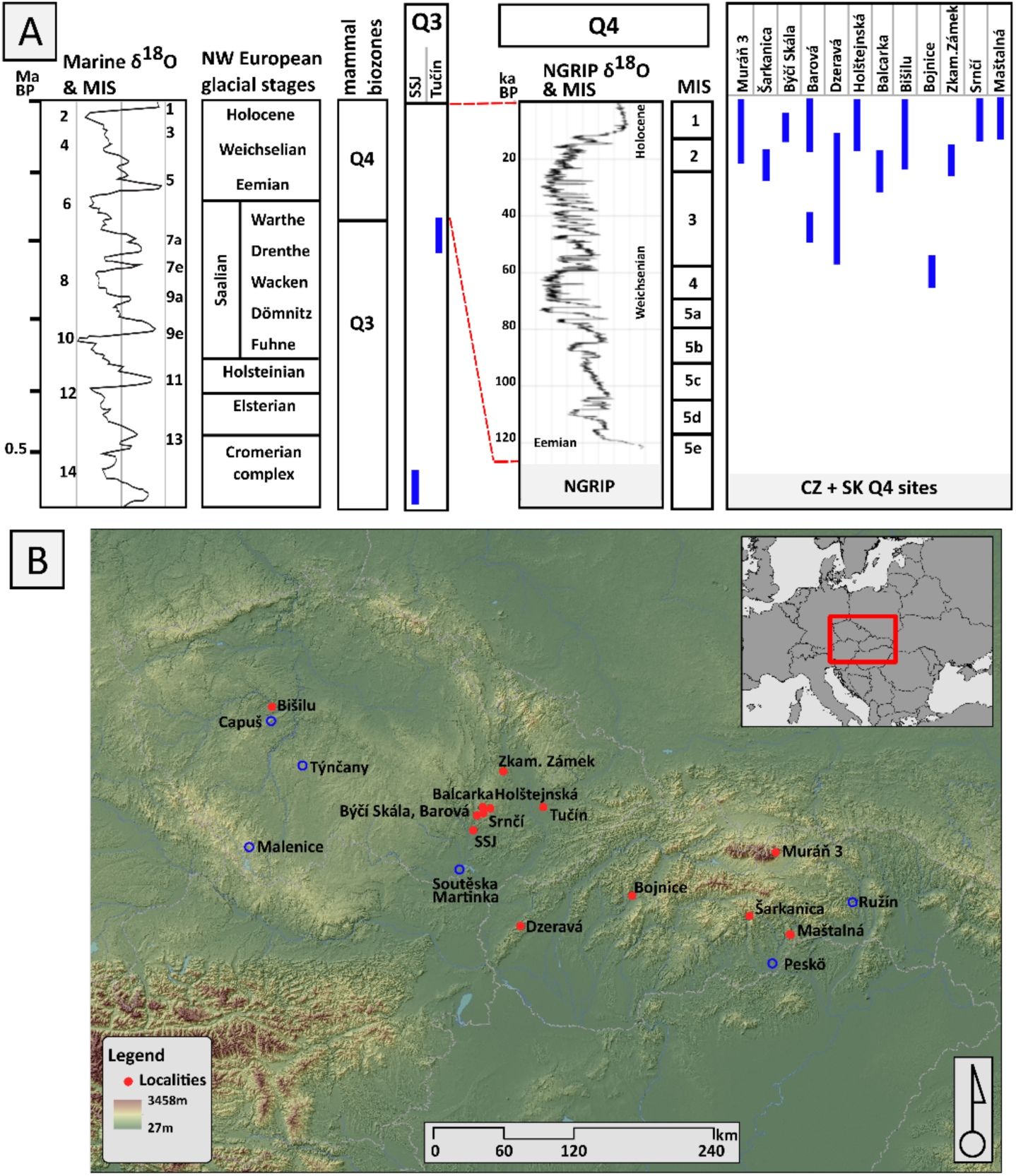
Stratigraphic and geographic position of localities. (**A**) Stratigraphic position of the studied localities shown in relation to the global marine oxygen isotope record (MIS), NW European glacial stages, and the NGRIP δ¹⁸O record. Correlation of regional stratigraphic units with MIS follows the global marine isotope framework (Bassinot et al. 1994; Lisiecki and Raymo 2005) and its calibration against Greenland ice-core records and NW European stratigraphy (NGRIP Members 2004; Andersen et al. 2006). (**B**) Geographic position: studied localities are indicated by a red circle, other localities discussed in the paper are indicated by a blue hollow circle.

The analyzed sites cover a broad temporal and geographic range across Central Europe: Middle Pleistocene: Stránska skála (MIS 12), Tučín (MIS 6). Late Pleistocene: Bojnice (MIS 4), Balcarka and Zkamenělý Zámek (MIS 3), Šarkanica (MIS 2), and continuous sedimentary sequences covering deeper stages of MIS 3–2 period (i.e., from 50 to 20ky B.P.: Dzeravá skala, Barová partim) and those covering the period from LGM (or late MIS 3) to late Holocene (MIS 2–1: Muráň 3, Bišilu, Holštejnská, Maštalná, Srnčí, Barová partim, Býčí Skála). Stratigraphic interpretation of community samples reflects their position in the sedimentary sequence, hints of biostratigraphic correlation (see Horáček and Ložek 1988; Horáček et al. 2002; Kaminská et al. 2005), and radiocarbon dating (surveyed in detail by Horáčková et al. 2014 (appendix); Horáček et al. 2015; Baca et al. 2019 (appendix)). The geographic position of individual sites and their stratigraphic settings are in Fig. 1; all further details, as well as characteristics of their *S. anglicus* populations, are in Supplementary file S1, and the sample sizes from community samples are in Supplementary file S2. Layers in multilayered sequences are denoted with a slash (/x).

The stratigraphic position of samples is expressed in terms of standard subdivisions of the Late Pleistocene–Holocene past (Walker et al. 2019; Sommer 2020), biostratigraphic units proposed by Horáček and Ložek (1988), and/or Marine Isotopic Stages (MIS) (Railsback et al. 2015). The taxonomic framework follows proposals by Kryštufek and Shenbrot (2022), particularly regarding the independent generic status of *Stenocranius*, which contrasts with its former arrangement in *Lasiopodomys* (cf., e.g., Baca et al. 2019).

## Methods

Specimens were photographed at Masaryk University, Brno, using a 3D microscope (Hirox MX-G5040SZ). Tooth orientation was standardized using a custom sample leveling press. Two-dimensional landmark digitization was conducted using TPS series software (Rohlf 2015).

*Landmark schemes and measurements:* Three landmark schemes were employed: A 24–landmark scheme covering the complete occlusal outline and major morphological structures. A 12–landmark scheme focused on the anteroconid complex. A 6–landmark scheme targeting the most variable points identified in previous analyses. Morphometric measurements (taken from the photographs with aid of TPSDig software) were based on Van der Meulen and Zagwijn (1974) and Nadachowski 1982 and adapted to integrate the landmarking approach of Wallace (2006). The positions of the landmarks, the descriptive scheme, and the linear variables are shown in Fig. 2.

**Figure 2.**
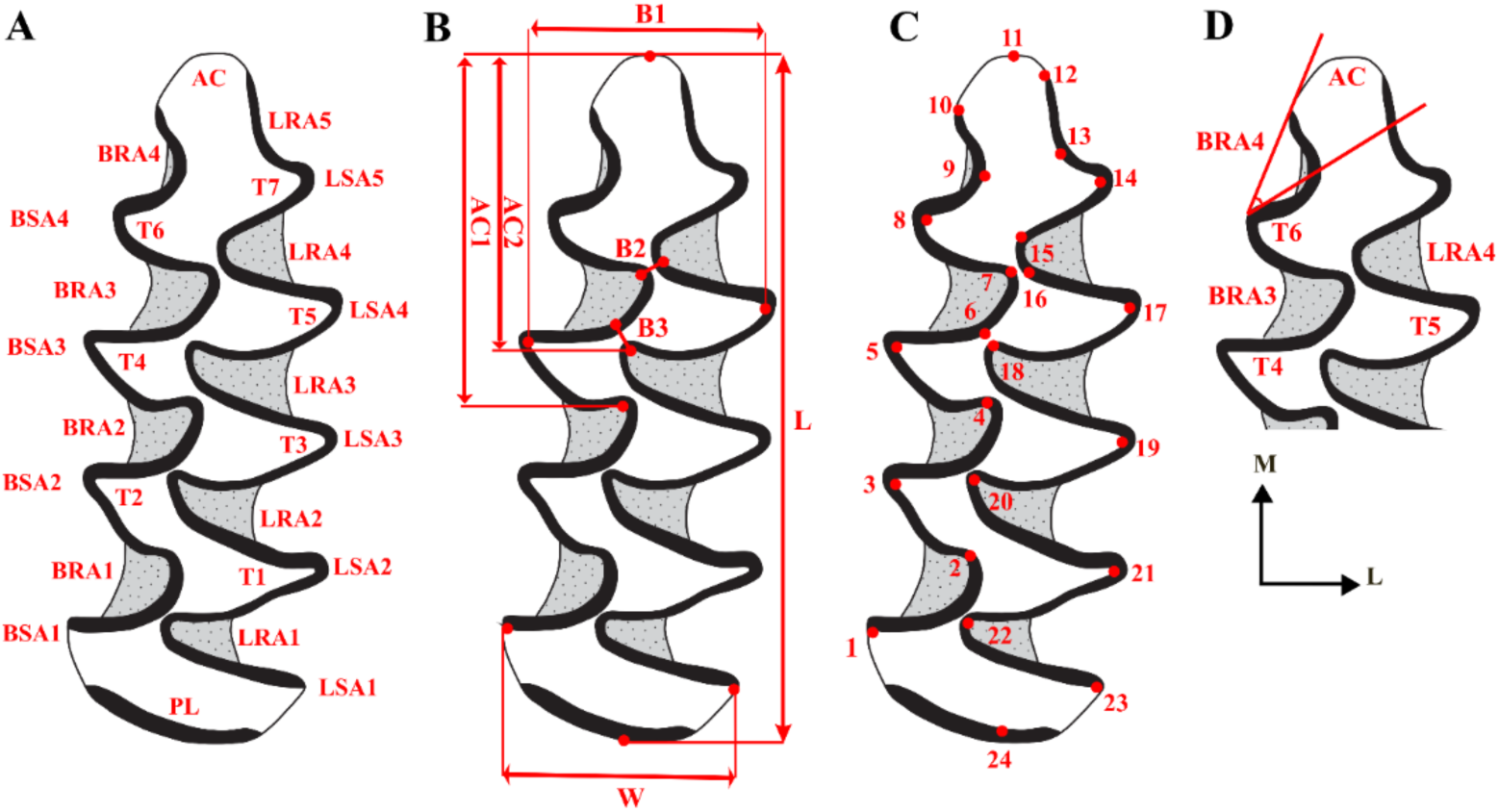
Nomenclature, measurements, landmark scheme, and morphotype analysis of the first lower molar in narrow-headed voles. (**A**) Nomenclature for the description of the first lower molar of the narrow-headed vole. AC: anterior cap; LRA: lingual reentrant angle; LSA: lingual salient angle; BRA: buccal reentrant angle; BSA: buccal salient angle; T: triangle; PL: posterior lobe. (**B**) Measurements of the first lower molar (m1) applied in this study: L: length of the tooth; AC1: length of the anteroconid complex; AC2: length of the anteroconid mesial cap; B1: width of the tooth; B2: width LRA4–BRA3; B3: width BRA3–LRA3. Note: some literary sources (e.g. Van der Meulen and Zagwijn 1974; Nadachowski 1982) denote the same measurements by different labels, i.e., AC1 = a; B1 = W; B2 = b; B3 = c. (**C**) Landmark scheme after Wallace 2006. (**D**) Morphotype measurement after Smirnov et al. 1986; Ponomarev and Puzachenko 2017.

Morphotypes were classified using both (i) BRA4 angle–based system (after Smirnov et al. 1986; Ponomarev and Puzachenko 2017) measured as the degree of tilt of the fourth buccal re-entrant angle on the first lower molar, and (ii) Nadachowski’s (1982) structural classification, which distinguishes 13 morphotypes (A–M) based on enamel field arrangement and developmental complexity. See Fig. 3 for details.

### Morphometric and Statistical Analyses

Geometric morphometric analyses were conducted using MorphoJ (Klingenberg 2011) and PAST5.3 (Hammer et al. 2001). Goodall’s F-values (Goodall 1991) and Pillai’s trace (Pillai 1955) were applied for basic between-sample comparisons. After Procrustes superimposition (Rohlf and Slice 1990), we conducted Principal Component Analysis (PCA) (Jolliffe 2002) to explore shape variance. Canonical Variate Analysis (CVA) (Fisher 1936; Klingenberg 2011) to assess stratigraphic and geographic clustering. Procrustes ANOVA (Goodall 1991; Klingenberg and Monteiro 2005) to test the effects of geography (locality) and stratigraphy (MIS) on molar shape. Discriminant Function Analysis (DFA) (Klecka 1980) to evaluate classification success between groups. Procrustes and Mahalanobis distances were used to quantify divergence between populations. Regression analyses explored trends in size and shape across stratigraphical and spatial units. Morphotype frequency dynamics were evaluated using Shannon diversity indices (H′) and coefficient of variation (CV), with additional consideration of skewness and kurtosis in key traits (L, W, AC1, AC2, B1–B3). Standard statistics and between-samples comparisons (using Pearson correlation, Bray-Curtis distances, significance test of differences, etc.) were computed in Microsoft Excel and PAST5.3 software and are presented in full in Supplementary file S2.

**Figure 3.**
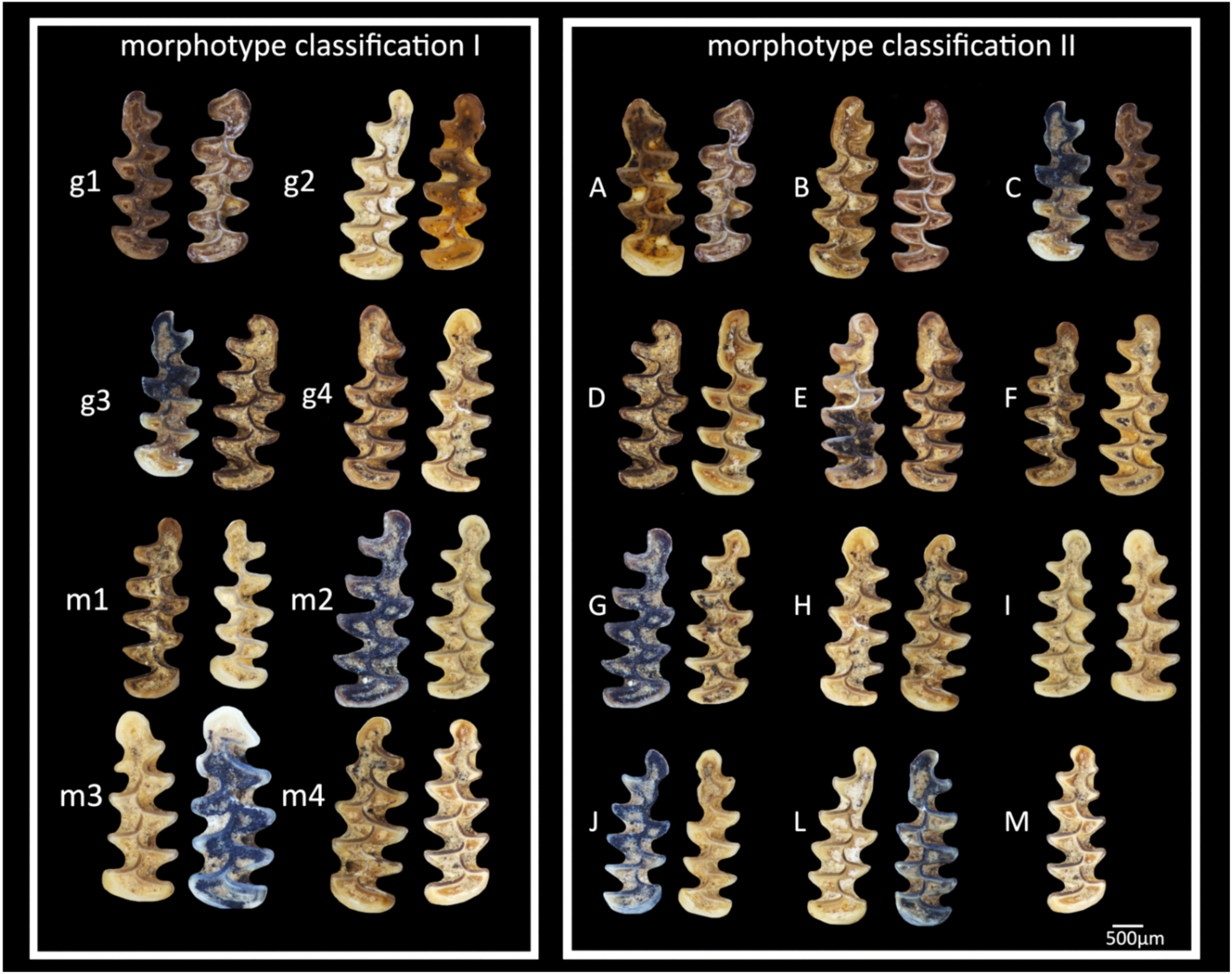
Morphotype classification schemes used in the study. Morphotype classification I is based on the BRA4 angle–based system after Smirnov et al. 1986; Ponomarev and Puzachenko 2017 (g1–m4). Morphotype classification II follows Nadachowski 1982 (A–M). Scale bar: 500 μm.

## Results

### (i) Stratigraphical distribution of *S. anglicus* in the Vistulian and Holocene community samples

The database of small-mammal assemblages from the Late Pleistocene and Holocene in the Czech Republic and Slovakia (covering the samples we physically reexamined, i.e., not all sites available in the countries) comprises 775 community samples from 102 sites, totaling 24100 individuals (MNI). Most of them originated from continuous sedimentary series mainly covering the period from the late glacial to the Recent. *Stenocranius anglicus,* with a MNI of 6352 in 235 community samples, is the most common element in the total sample. It was recorded in almost all communities dated to MIS 3 and MIS 2, including the LGM, where it appeared as the dominant element of the community (composing 10–70% of the MNI). Of the total sample, 533 records (with MNI > 15) are considered real community samples; those with *S. anglicus* detected in regions under study are listed in Table 1. Dominance of the species in these samples varies from 4.6–43.9% (20% on average) for MIS 3 and 10.2–78.2% (36% on average) for MIS 2. Yet, regarding the poor representation of MIS 2 stage (recorded mostly as erosion events preceding sedimentation of the studied series), the vast majority of *S. anglicus* records (62 = 31.8%) come from the Late Pleistocene stages. Besides, the species is regularly represented also in assemblages of the early Holocene ranging from 1.1–66.6% (22.6% on average) in 37 communities of the Preboreal stage, 0.4–41% (15.18% on average) in 21 communities of the Boreal stage, and 0.8–14.7% (7.4% on average) in 8 communities of the middle Holocene. Notwithstanding the middle Holocene records, often represented by only a single specimen, which could be considered doubtful in some cases, quite robustly support the survival of *S. anglicus* in most regions throughout the early Holocene. Selected examples are surveyed below.

**Table 1.** Number of relevant community samples (MNI > 15) of the Vistulian and Holocene age from individual regions of the Czech Republic and Slovakia, and the number of those with *S. anglicus*.

| Region | Total community samples (MNI >15) | Communities (MNI > 15) with *S. anglicus* |  |  |  |  |  |  | Total | Number of studied samples |
| --- | --- | --- | --- | --- | --- | --- | --- | --- | --- | --- |
|  |  | MIS 4 | MIS 3 | MIS 2 | Late glacial | Preboreal | Boreal | Middle/ late Holocene |  |  |
| Bohemian karst | 148 |  | 7 | 6 | 13 | 5 | 2 |  | 33 | 3 |
| Central Bohemia | 7 |  |  |  | 1 | 1 |  |  | 2 |  |
| Southern Bohemia | 3 |  |  | 2 | 1 |  |  |  | 3 |  |
| Northern Bohemia | 59 |  |  |  |  |  |  |  | 0 |  |
| Northern Moravia | 13 |  |  | 3 | 3 | 3 | 1 | 1 | 11 | 2 |
| Moravian karst | 115 | 1 | 1 | 10 | 28 | 16 | 9 | 2 | 67 | 20 |
| Southern Moravia | 41 |  | 2 | 2 | 2 | 5 | 5 | 2 | 18 |  |
| Western Slovakia | 25 | 3 | 9 | 2 | 1 |  |  |  | 15 | 10 |
| Southeastern Slovakia | 25 | 1 | 4 | 2 | 4 | 5 | 2 | 2 | 20 | 1 |
| Slovakian karst | 57 |  |  | 2 | 5 | 2 | 1 | 1 | 11 | 7 |
| Northern Slovakia | 40 |  | 3 | 7 | 4 |  | 1 |  | 15 | 4 |
| Total | 533 | 5 | 26 | 36 | 62 | 37 | 21 | 8 | 195 | 48 |
| % |  | 2.6 | 13.3 | 18.5 | 31.8 | 19.0 | 10.8 | 4.1 | 100.0 |  |

### (ii) Phenotype characteristics of *Stenocranius anglicus*

#### (i) Linear morphometric variables

Comparing overall variation in mean values of standard linear measurements (21 variables) across all populations, we found no significant differences between samples (ANOVA, F (df = 10) = 0.029; p = 1; Kruskal–Wallis test: H = 0.774; p = 0.999). All samples exhibited significant normal distributions (Shapiro–Wilk W = 0.808–0.835; p = 0.0035–0,0011) and high similarity in metric profile as indicated by high between-sample correlations (r: 0.991–0.999), high Bray–Curtis similarity values (0.93–0.97), and low Mahalanobis distances (0.581–1.271). These results suggest that all samples represent a single phenotypic unit consistent with the taxonomic identity of the mid-European *Stenocranius anglicus*.

However, the detailed analysis revealed notable variations in both metric and proportional characteristics across different stratigraphic and geographic contexts. In general, specimens from MIS 3 and older showed larger molar dimensions than those from Late Vistulian/Early Holocene localities. The largest molars were found in Bojnice (MIS 4) and Bišilu (MIS 2), with maximum recorded lengths of 3.900 mm and 3.088 mm, respectively. In contrast, the smallest were from Maštalná (MIS 1) and Stránska skála SSJ (MIS 12/14), with minimum lengths of 2.069 mm and 2.162 mm, respectively. The width of the molars followed a similar pattern, with the widest tooth recorded in Bojnice (1.333 mm) and the narrowest in Maštalná (0.660 mm).

Among proportional characteristics, the anteroconid complex showed substantial variation, with the highest relative values observed in Zkamenělý Zámek and Bojnice, whereas the lowest values appeared in Holštejnská and Srnčí. The most extreme AC1/L ratio (relative anteroconid length) was observed in Stránska skála SSJ (0.562) and Bišilu (0.574), while the lowest values were recorded in Tučín (0.503) and Barová (0.502). Similarly, the AC2/AC1 ratio varied, with the highest values in Maštalná (0.832) and the lowest in Zkamenělý Zámek (0.775). The relative width B1/W was the largest in Bišilu (1.148) and the smallest in Zkamenělý Zámek (0.944). The variability in shape proportions, as measured by coefficients of variation, was higher in some Late Vistulian/Early Holocene samples, particularly in features of the anteroconid complex (AC1 and AC2). Detailed survey of biometric data is available in Supplementary files S1-S3.

Geographically, specimens from Moravian localities (e.g., Stránska skála SSJ, Balcarka, Zkamenělý Zámek) exhibited slight but consistent differences from Slovak samples (e.g., Šarkanica, Bojnice, Muráň 3), particularly in the width and proportions of the anteroconid and basin structures. The highest coefficients of variation (CV) were found in Šarkanica, especially for AC2 and AC1/L, suggesting higher morphological diversity, while the lowest variability was observed in Holštejnská and Barová. Extreme skewness and kurtosis values for specific characteristics, such as the B2 and B3 proportions in Šarkanica and Tučín, the sites with the highest dominance of *S. anglicus*, suggest strong tendencies toward rearrangements of specific phenotypic traits.

#### (ii) Geometric morphometrics

First, we compared the strength of three different landmark schemes against the geographical and stratigraphical positions of particular samples using ANOVA and Canonical Variates Analysis (CVA). Both analyses revealed that the geographical position of the localities, as well as their stratigraphy (recorded by the marine isotope stage (MIS) of the respective locality/stratigraphic layers), significantly affect shape variation across all landmark schemes.

Procrustes ANOVA for the 24–landmark scheme showed a strong effect of locality on shape (F = 28.28, p < 0.0001; Pillai’s trace = 2.51) and MIS (F = 51.12, p < 0.0001; Pillai’s trace = 1.67). When the number of landmarks was reduced to 12, Goodall’s F–values slightly increased for both locality (F = 31.63) and MIS (F = 57.86), but Pillai’s trace values decreased (1.86 for locality; 1.33 for MIS), indicating a reduction in the proportion of variance explained. The 6–landmark scheme showed the weakest effects, with locality (F = 29.19; Pillai’s trace = 0.85) and MIS (F = 56.47; Pillai’s trace = 0.63), confirming that shape differentiation remains significant but is notably weaker with fewer landmarks (Table 2). Overall, the effect of geographical position consistently shows more substantial shape variation effects than stratigraphy (MIS) across all landmark schemes. Locality has consistently higher Pillai’s trace values than MIS in the 24–landmark scheme (2.51 vs. 1.67), meaning that geographic differences explain more of the shape variation than stratigraphic differences. Goodall’s F–values are higher for MIS than for locality in the 12– and 6–landmark schemes, but this likely reflects statistical sensitivity rather than a true biological pattern (Table 2).

**Table 2.**
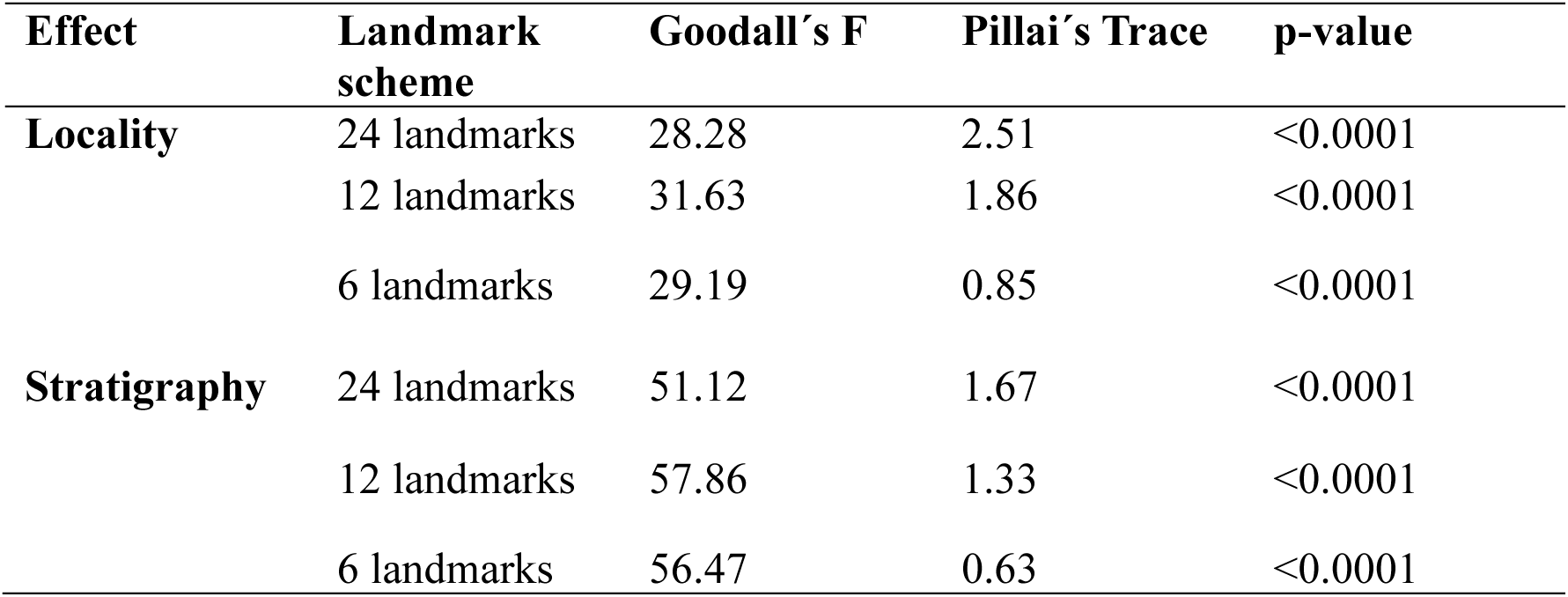
Comparison of 24–, 12–, and 6– landmark schemes.

Mahalanobis and Procrustes distances are generally larger for geographical comparisons, suggesting that shape differences among localities are greater than those among stratigraphic stages. Geographical position could therefore be a more dominant factor influencing shape variation. However, MIS still has a significant effect, particularly in the full landmark dataset, although it explains less overall shape variation. This pattern holds across all landmark schemes, but when reducing the number of landmarks, the stratigraphic effect appears less affected than the locality effect. Pairwise Mahalanobis distances indicated that shape differentiation among localities and MIS groups was strongest in the 24–landmark scheme. For locality, the highest Mahalanobis distance was observed between Stránska skála SSJ and Barová (5.82), while the lowest (1.74) was between Maštalná and Bojnice. In the 12–landmark scheme, the highest distance decreased to 5.14, and in the 6–landmark scheme, it dropped to 2.34, indicating a progressive reduction in shape distinctiveness. For stratigraphy, the 24–landmark scheme showed the highest Mahalanobis distance spanned between MIS 12 and MIS 4 (5.10), i.e., Stránska skála SSJ and Bojnice, while the 6–landmark scheme reduced the span to 2.93. The smallest distances were also reduced, suggesting that fewer landmarks lead to less separation among stratigraphic groups. Procrustes distances followed a similar trend. While group separation remained significant in all schemes (all p < 0.0001), Procrustes distances were consistently lower in the 6–landmark scheme, meaning the amount of shape variation captured was reduced (Table 3).

**Table 3.**
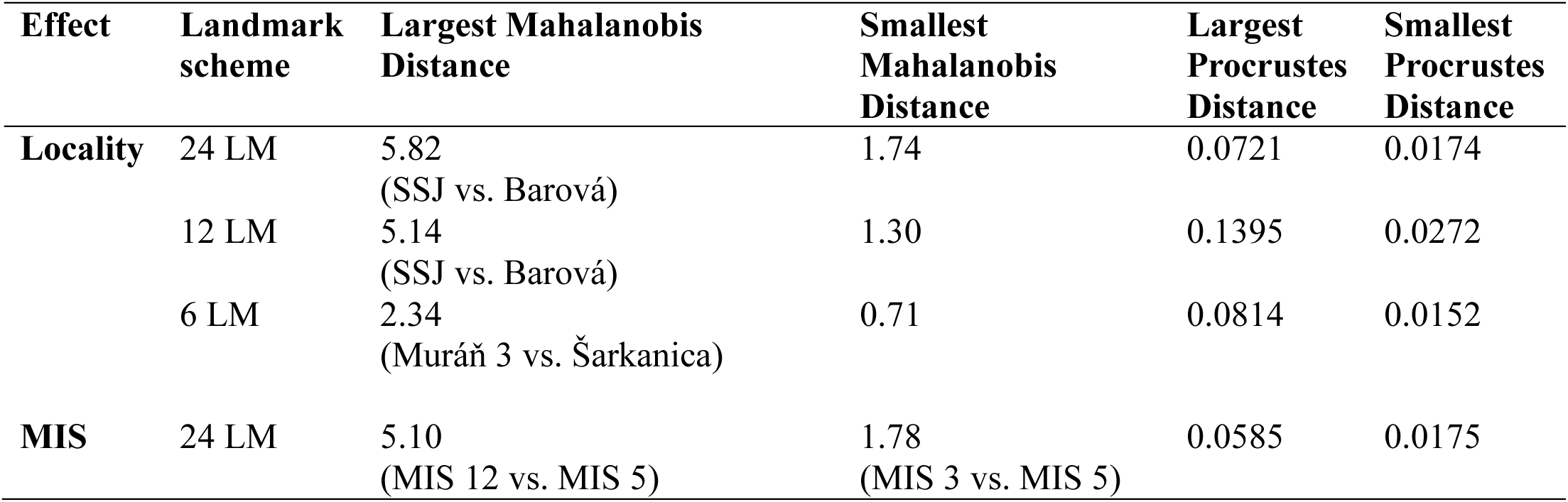
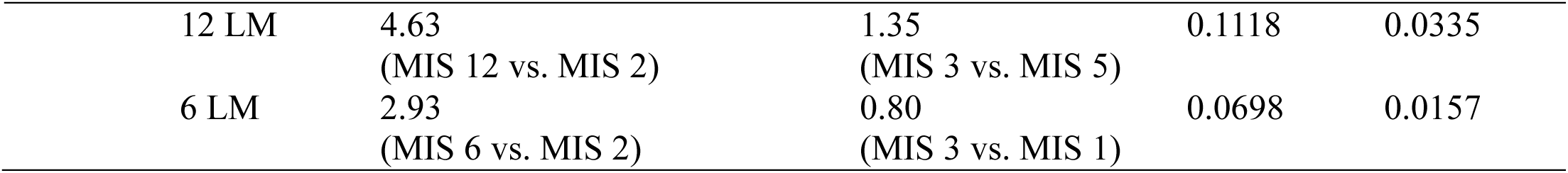
Group separation (Mahalanobis and Procrustes distances). LM = number of landmarks.

Overall, the 24–landmark scheme provides the most comprehensive representation of shape variation, while the 12–landmark scheme retains significant, though slightly reduced, explanatory power. The 6–landmark scheme still detects locality and MIS effects but at a much weaker level, suggesting that landmark reduction may lead to a loss of biologically meaningful shape variation. These findings indicate that landmark reduction should be carefully considered, particularly when studying complex morphological differences across space and time.

The correspondence between stratigraphic classification based on CVA (Canonical Variates Analysis) and the actual stratigraphic position of particular samples revealed varying levels of classification accuracy. The highest classification success was observed in MIS 6 (Tučín), with 88% of specimens correctly classified, indicating distinct molar morphology during that glacial period. Similarly, the Last Glacial Maximum (MIS 2, Šarkanica) achieved 80% accuracy, supporting the notion that glacial periods produced morphologically cohesive vole populations. MIS 12 (Stránska skála SSJ) showed moderate accuracy (58%), consistent with its early Middle Pleistocene position and potential ancestral morphology. In contrast, Late Vistulian/ Early Holocene MIS 1 (multiple localities) had 69.5% accuracy, with notable misclassifications into pre–LGM periods, suggesting morphological convergence or stabilization. MIS 3 (Balcarka, Zkamenělý Zámek) had 71% accuracy, while MIS 4 (Bojnice) was lower at 38.6%, with substantial misclassification into MIS 3 and MIS 1, implying morphological overlap during this period. Regarding the centroid positions of particular MIS units, those of the Middle Pleistocene age (i.e., MIS 12 and MIS 6) show considerable distance from the Late Pleistocene and Holocene samples. At the same time, MIS 4 and MIS 2 (both extreme pleniglacials) appeared close to each other, similarly to MIS 3 and MIS 1 (Fig. 4).

**Figure 4.**
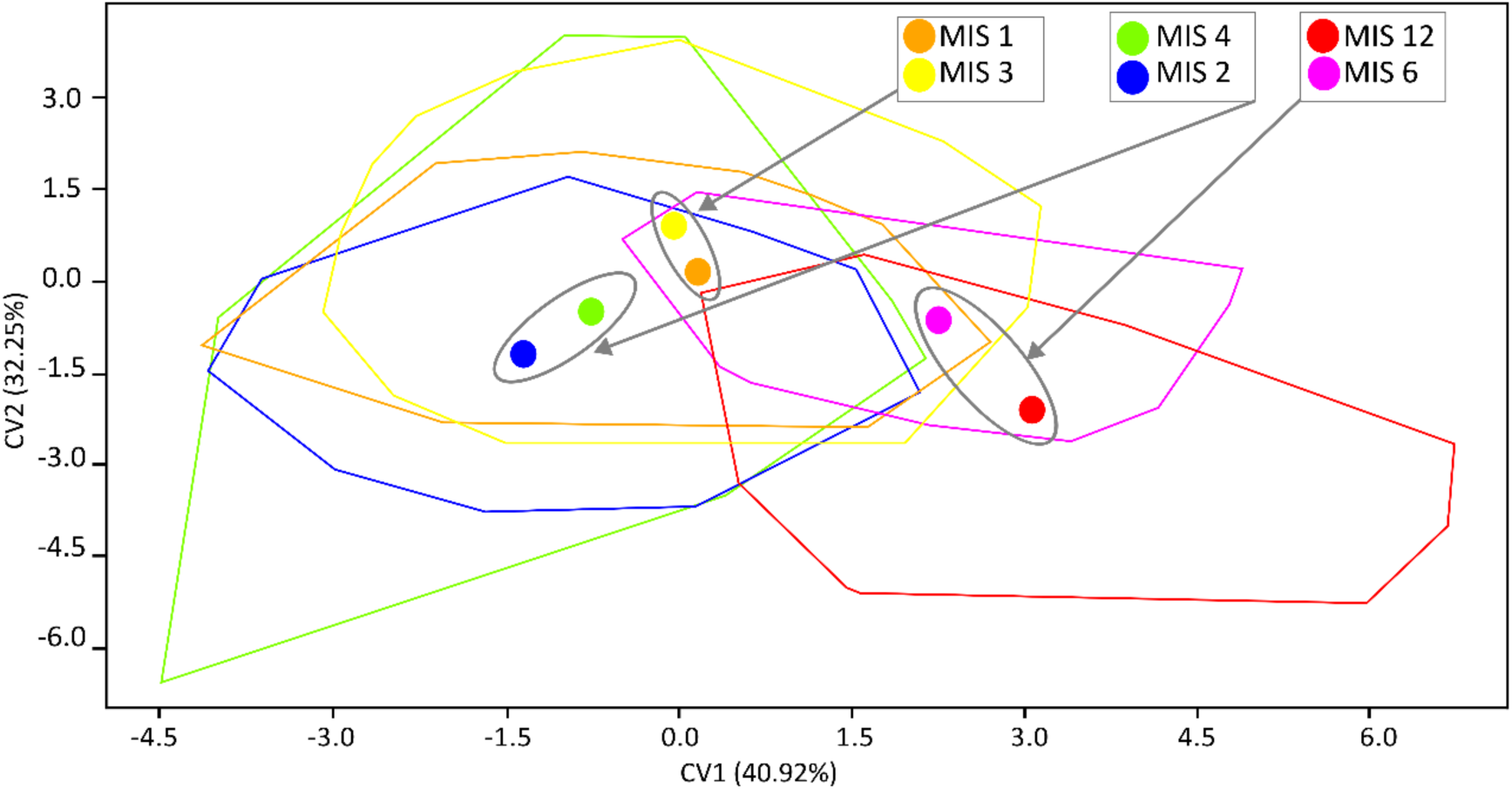
Canonical variates analysis of individual stratigraphic units. The overall pattern of morphometric variation among individual stratigraphic units is shown in a scatter plot of Canonical Variate Analysis. The corresponding colors of MIS units indicate their variation clouds and centroids.

Across localities, the classification accuracy from the CVA varied notably. Tučín displayed the highest correct classification rate at 82%, suggesting that its narrow-headed vole molars possessed distinct morphological traits, likely shaped by the MIS 6 glacial environment. Other localities with relatively high accuracy included Muráň 3 (68.1%), Býčí Skála (66.5%), Šarkanica (64.2%), and Balcarka (63.2%), all of which indicate strong morphological differentiation. In contrast, Bojnice had the lowest accuracy (24.1%) and was frequently misclassified as Šarkanica, suggesting morphological convergence or shared traits.

Similarly, Maštalná, Zkamenělý Zámek, and Srnčí showed moderate to low accuracy (around 41%), with misclassifications spread across several groups, indicating either internal variability or morphological overlap with other localities. Worth mentioning is a resemblance of the sites Barová (57.3%), Dzeravá (50.0%), and Býčí Skála, which displayed moderate classification success, corresponding to their similarities in other phenotype comparisons, and the distant position of Stránska skála SSJ (with 58% of correct classification) and Tučín, both representing a Middle Pleistocene context (comp. centroid positions in Fig. 5).

**Figure 5.**
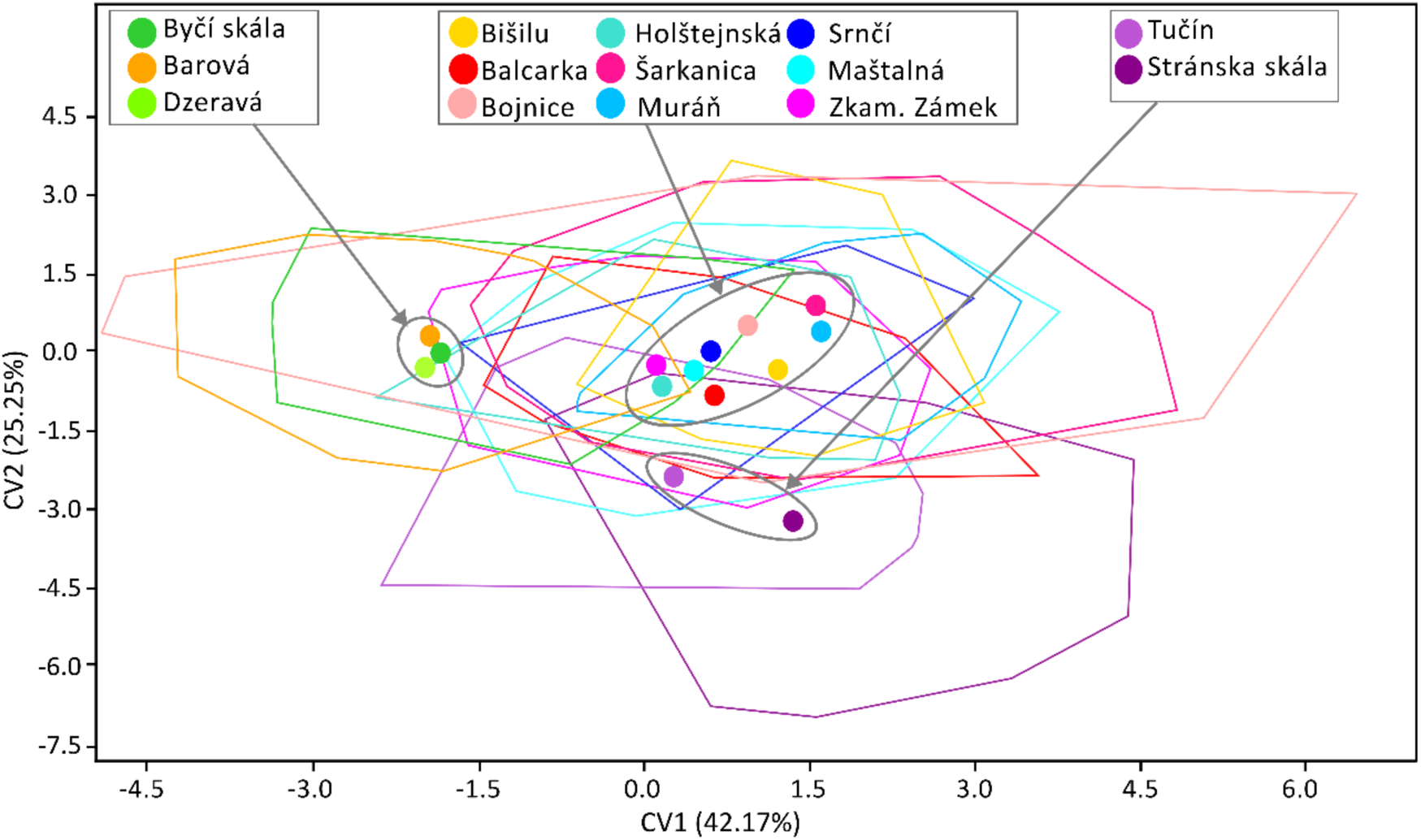
Canonical variates of individual localities. The overall pattern of morphometric variation among individual localities is shown in the scatter plot of the Canonical Variates Analysis. Centroids and variation clouds are indicated by corresponding colors of the studied localities.

Both classification schemes demonstrated the highest accuracy for cold-adapted populations during glacial stages-Tučín (MIS 6) and Šarkanica (MIS 2) stood out in both analyses with high classification rates (∼82% and ∼80%, respectively). Localities tied to transitional periods, such as Bojnice (MIS 4), showed poor classification accuracy in both schemes (24% locality-based, 39% stratigraphic), underscoring morphological variability or convergence during these times. Holocene localities had moderate success in both systems (∼50–57%), suggesting some morphological stabilization before extinction. Overall, both classification matrices show consistent patterns that highlight the narrow-headed vole’s evolutionary responses to climatic fluctuations. A relatively low number of correctly classified specimens (locality ∼55%; stratigraphy ∼70%) indicates that the phenotypic characteristics of the European narrow-headed vole are stable but exhibit plasticity, with a high potential for phenotypic rearrangements during periods of unfavorable environmental conditions (e.g., Holocene localities).

#### (iii) Morphotype diversity

i. based on the morphotype scheme I - BRA4 angle classification (Ponomarev and Puzachenko 2017): During the Middle to Late Pleistocene (pre–LGM), diversity values ranged widely across both Czech and Slovak localities (Fig. 6). In the oldest locality, Stránska skála SSJ, the diversity index was moderate (H′ = 1.637). This pattern continued into Tučín (H′ = 1.585) and Bojnice (H′ = 1.639), suggesting relatively stable diversity during early glacial cycles. A notable increase in diversity occurred during MIS 3, particularly at Zkamenělý Zámek (Czech Republic), which recorded the highest pre–LGM diversity (H′ = 1.980) across all localities, exceeding values in all other samples. Similarly, Balcarka showed high diversity (H′ = 1.787). Contrasting patterns of diversity across Slovakia and the Czech Republic were evident during the Last Glacial Maximum (LGM). In Šarkanica (Slovakia), an apparent decline in diversity was observed (H′ = 1.520). While the Muráň 3 sequence (Slovakia), Muráň 3/6 (24.4ky BP) exhibited one of the highest diversity values recorded during the LGM (H′ = 1.923), Muráň 3/4 (16.3ky BP) and Muráň 3/3 declined to H′ = 1.230–1.332, foreshadowing population contractions toward the beginning of the Holocene. In the Czech Republic, diversity during the LGM suggests high morphological variability. Following the LGM, a general increase in diversity was evident across both countries, with most post–glacial localities exhibiting H′ values above 1.6. The highest diversity post–LGM was recorded in Maštalná /13 (H′ = 1.955) and Barová 14–10ky (H′ = 1.979), while Dzeravá /3 (Slovakia) reached H′ = 1.889, showing high variability across regions. In Holštejnská (Czech Republic), layer 5 (H′ = 1.759) and layer 6 (H′ = 1.864) showed elevated diversity, supporting a trend of increasing variability toward the Holocene. In Slovakia, the Dzeravá sequence showed stable high diversity (H′ = 1.796–1.889), consistent with long-term population stability or refugial continuity. Similarly, Býčí (Czech Republic) showed high diversity in layers dated to ∼11.2–12.1ky BP (H′ = 1.686–1.845), aligning with early Holocene warming. Overall, Czech localities (e.g., Balcarka, Barová) tended to exhibit slightly higher and more variable diversity compared to Slovak sites, which, while also diverse, showed more consistent values (Dzeravá and Muráň 3 sequences).
ii. based on the morphotype scheme II - morphotype classification after Nadachowski 1982 Morphological diversity assessed using Nadachowski’s (1982) classification proposal revealed trends that broadly align with those derived from the BRA4 angle-based classification, still with notable local and chronological distinctions (Fig.7). Stránska skála SSJ (MIS 12) exhibited lower diversity (H′ = 1.501), consistent with a more primitive morphotype spectrum and less morphological variation. Diversity then peaked during the MIS 5–3 interval, with the highest values at Bojnice (H′ = 1.93) and Zkamenělý Zámek (H′ = 1.843). Tučín (MIS 6) also showed relatively high diversity (H′ = 1.616), possibly reflecting morphotype turnover during the pre–LGM cold period. During the LGM (MIS 2), diversity remained moderate to high, as seen at Šarkanica (H′ = 1.527), suggesting the continued presence of multiple morphotypes and potentially adaptive responses to glacial conditions. However, some late LGM localities, such as Muráň 3/6 (H′ = 1.484) and Holštejnská /4 (H′ = 0.9503), began to exhibit decreasing diversity, indicating incipient morphological homogenization preceding the species’ decline. In the post–LGM (Late Vistulian/Early Holocene) period, diversity patterns varied geographically: Býčí Skála and Dzeravá localities generally retained moderate to high diversity (e.g., Býčí /8a: H′ = 1.831, Dzeravá /7: H′ = 1.713), suggesting refugial stability or persistent morphological variability until the species’ final disappearance. Srnčí and Maštalná populations showed lower and fluctuating diversity (e.g., Srnčí /8: H′ = 0.6365, Maštalná /10: H′ = 0.4506), likely reflecting small, isolated populations with restricted morphotype spectra at the edge of extinction. Notably, Barová layers and Maštalná /14–15 exhibited high Shannon diversity (H′ > 1.95), comparable to peak glacial values. In contrast, Holštejnská /4 and Maštalná /10 had the lowest diversity values (H′ = 0.9503 and 0.4506), underscoring their role as terminal occurrences of the species in the region.

**Figure 6.**
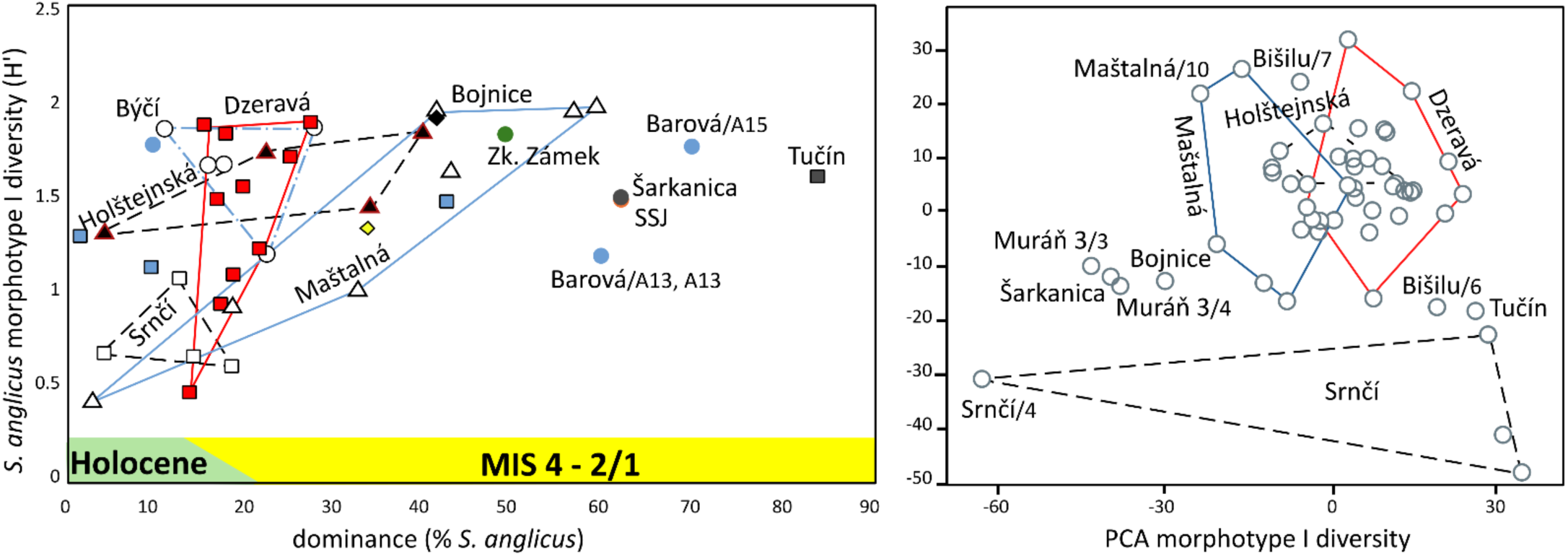
Morphotype diversity (group I) in populations of *S. anglicus* plotted against dominances of respective populations in communities of small ground mammals (left) and results of corresponding PCA (PC1 vs. PC2). Note a broad variation span in sequences of Dzeravá and Maštalná, covering a variation in the vast majority of other samples.

**Figure 7.**
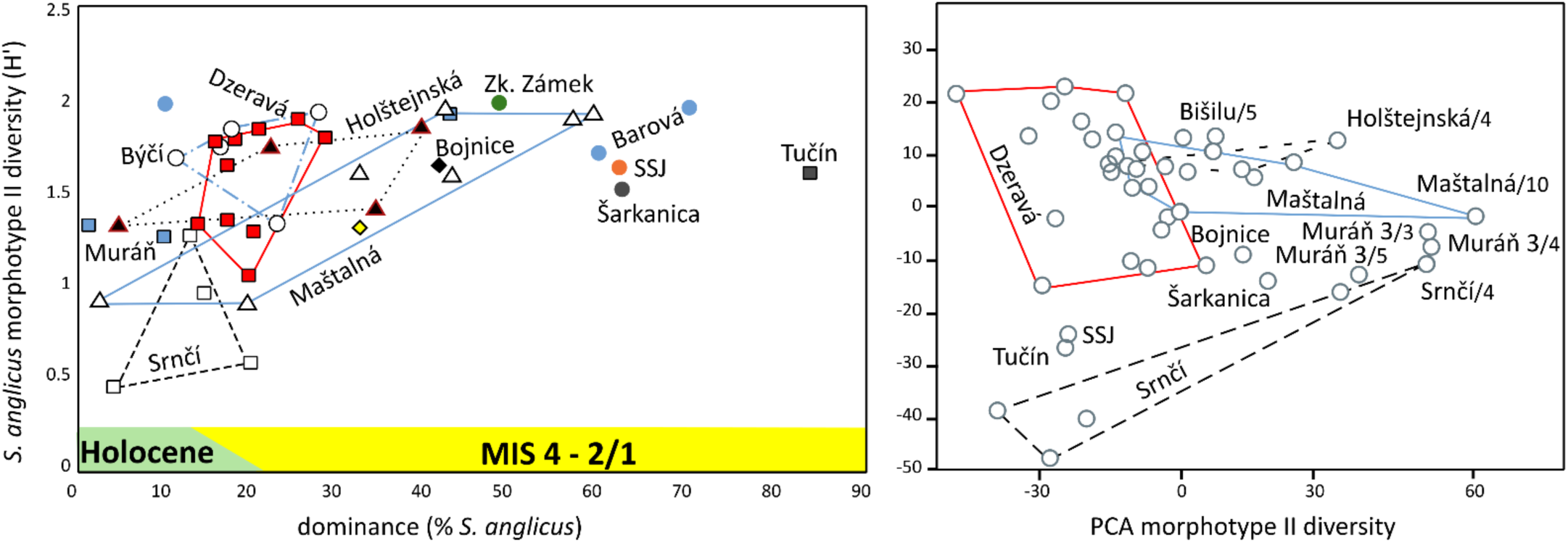
Morphotype diversity (group II) in populations of *S. anglicus* plotted against dominances of respective populations in communities of small ground mammals (left) and results of corresponding PCA (PC1 vs. PC2). Note a broad variation span in sequences of Dzeravá and Maštalná, covering a variation in the vast majority of other samples.

Overall patterns of these two approaches align (Fig. 8). Both classification systems identify high diversity during MIS 3–LGM and a post–LGM decline, but Nadachowski’s scheme (II) highlights fine-scale morphotype turnover and rare morphotype occurrences (J–M) in late periods.

**Figure 8.**
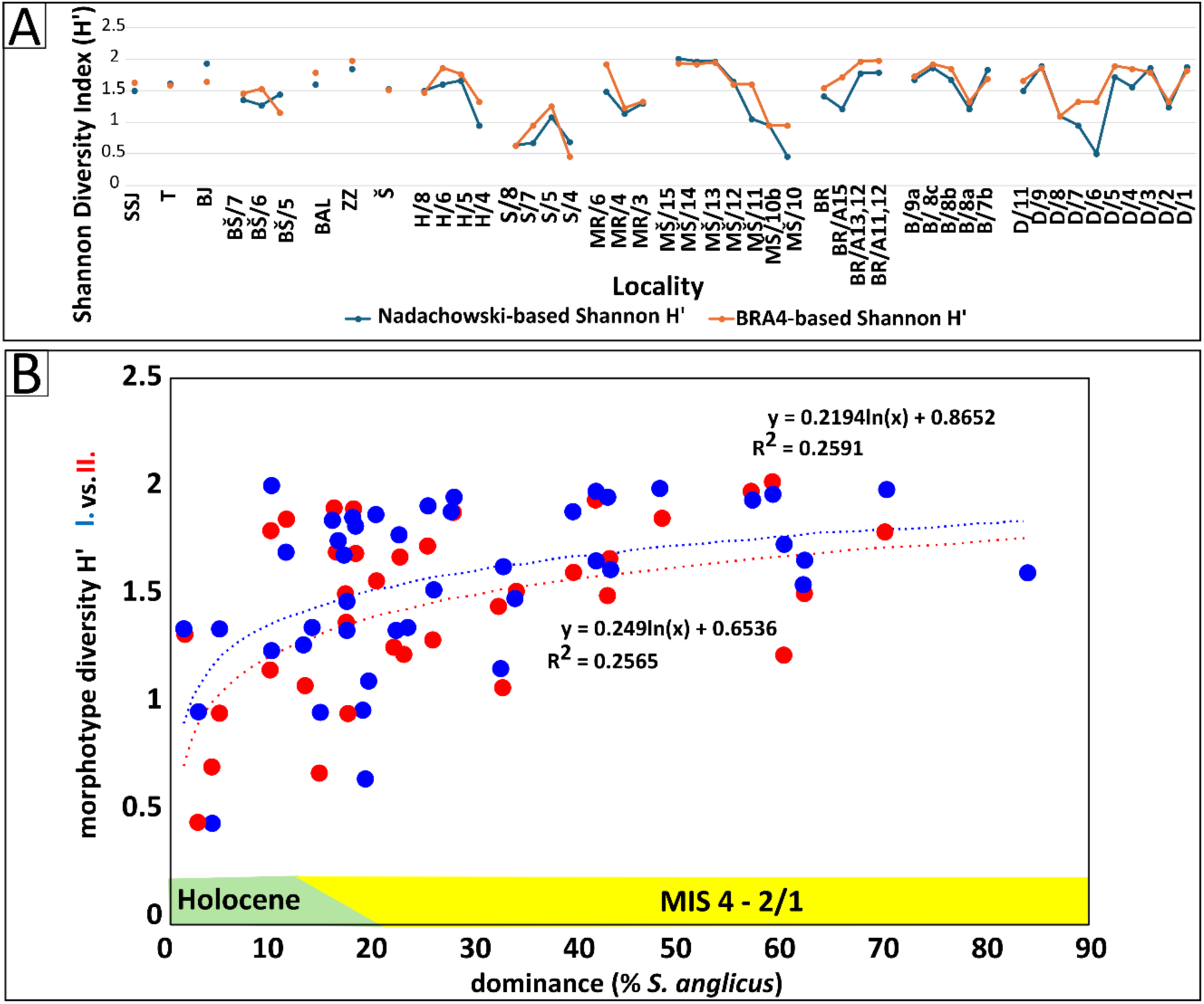
Comparisons of morphotype diversity. A: Shannon diversity in individual samples based on morphotype classification scheme I (A–M after Nadachowski 1982 - in red), and morphotype classification scheme II (g1–m4 BRA4–based - in blue). Values for community samples (**A**), plotted against the dominance of *S. anglicus* in respective communities (**B**). SSJ – Stránska Skála cave, T – Tučín, BJ – Bojnice, BŠ – Bišilu, BAL – Balcarka, ZZ – Zkamenělý Zámek, H – Holštejnská, S – Srnčí, MR – Muráň 3, MŠ – Maštalná, BR – Barová, B – Býčí Skála, D – Dzeravá skala

Geographic differences are more pronounced in II- Slovak localities, consistently showing higher diversity than the Bohemian and Moravian sites. It is markedly illustrated by a comparison of morphotype diversity (scheme II) in Czech Republic, Slovakia and Poland arranged in stratigraphical units proposed by Nadachowski (1982) (Fig. 9). It demonstrates similarity in the phenotype diversity trends among the Czech Republic (mostly Moravia) and Poland during the glacial stages, contrasting to the specific situation in the Slovak Carpathians (characterized by significantly higher diversity during MIS 4 – MIS 2). The characteristics of the Holocene samples, both in Slovakia and the Czech Republic, are situated quite apart from the cluster of glacial samples. Worth mentioning are also the diversity values respective to all studied samples (Czech and Slovak sites together- Fig. 9A), which (particularly in MIS 4–2 samples) considerably exceed values of individual regions (Fig. 9B), indicating thus a greatly pronounced effect of beta diversity in phenotype pattern of the species.

**Figure 9.**
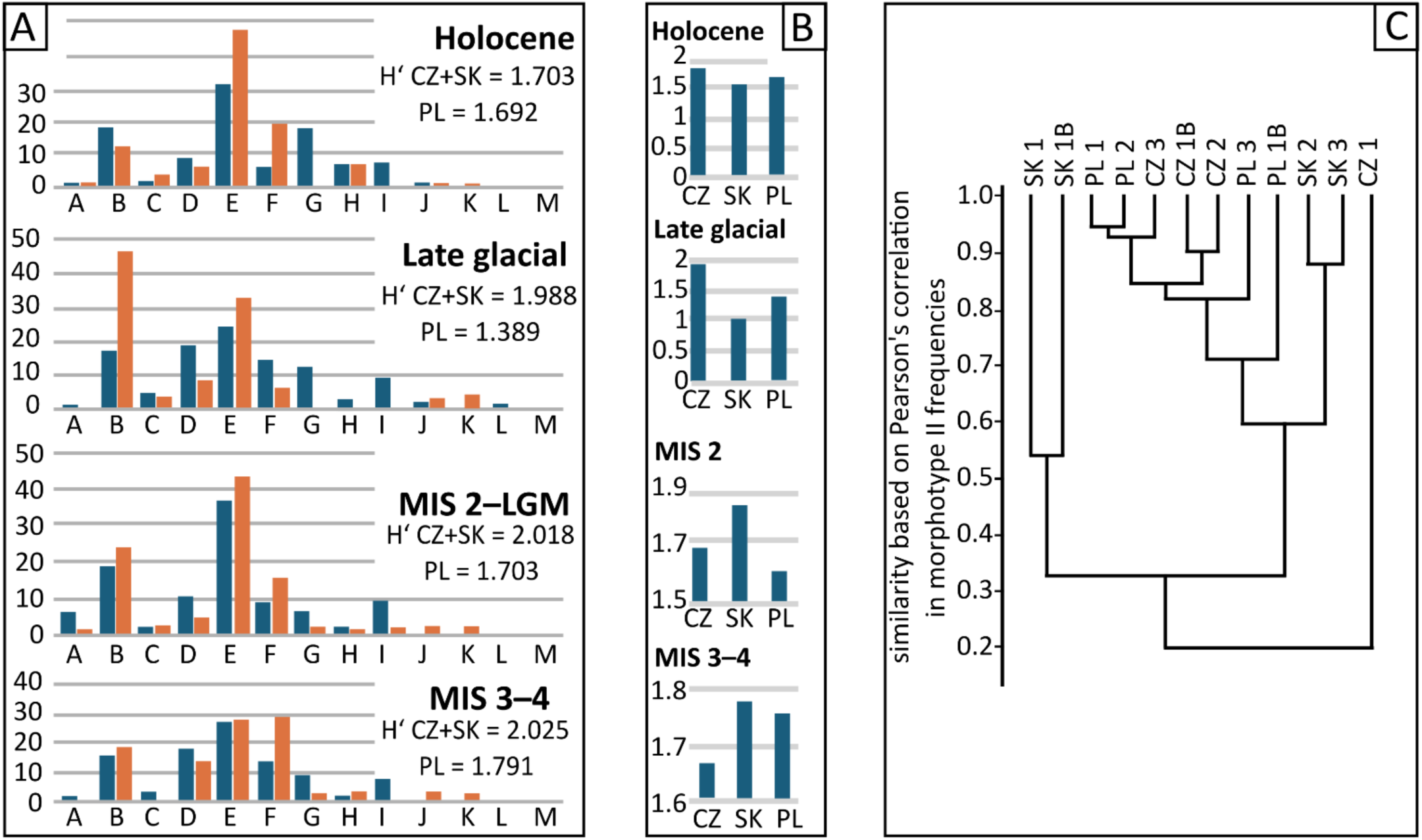
Phenotype variation among stratigraphic units in the Czech Republic, Slovakia, and Poland. (**A**) Mean frequencies of individual morphotypes (series II) and corresponding values of Shannon diversity (H′) in stratigraphic units proposed by Nadachowski (1982): own record (dark blue) vs. data from Polish sites (red). (**B**) Shannon diversity index H’ in samples from the Czech Republic (CZ), Slovakia (SK), and Poland (PL). (**C**) UPGMA clustering of mean morphotype frequencies in particular stratigraphic units (1- Holocene; 1b- late glacial; 2- MIS 2; 3- MIS 3–4) from Moravia and Bohemia (CZ), Slovakia (SK), and Poland (PL).

### (iii) Phenotype variation along the dataset means

To assess the degree of morphological divergence of the first lower molars across localities, each locality’s mean shape was compared to the dataset mean using Procrustes and Mahalanobis distances. Significant shape differences were observed in most localities, as confirmed by permutation tests (1000 runs; p < 0.001 unless noted otherwise) (Fig. 10).

**Figure 10.**
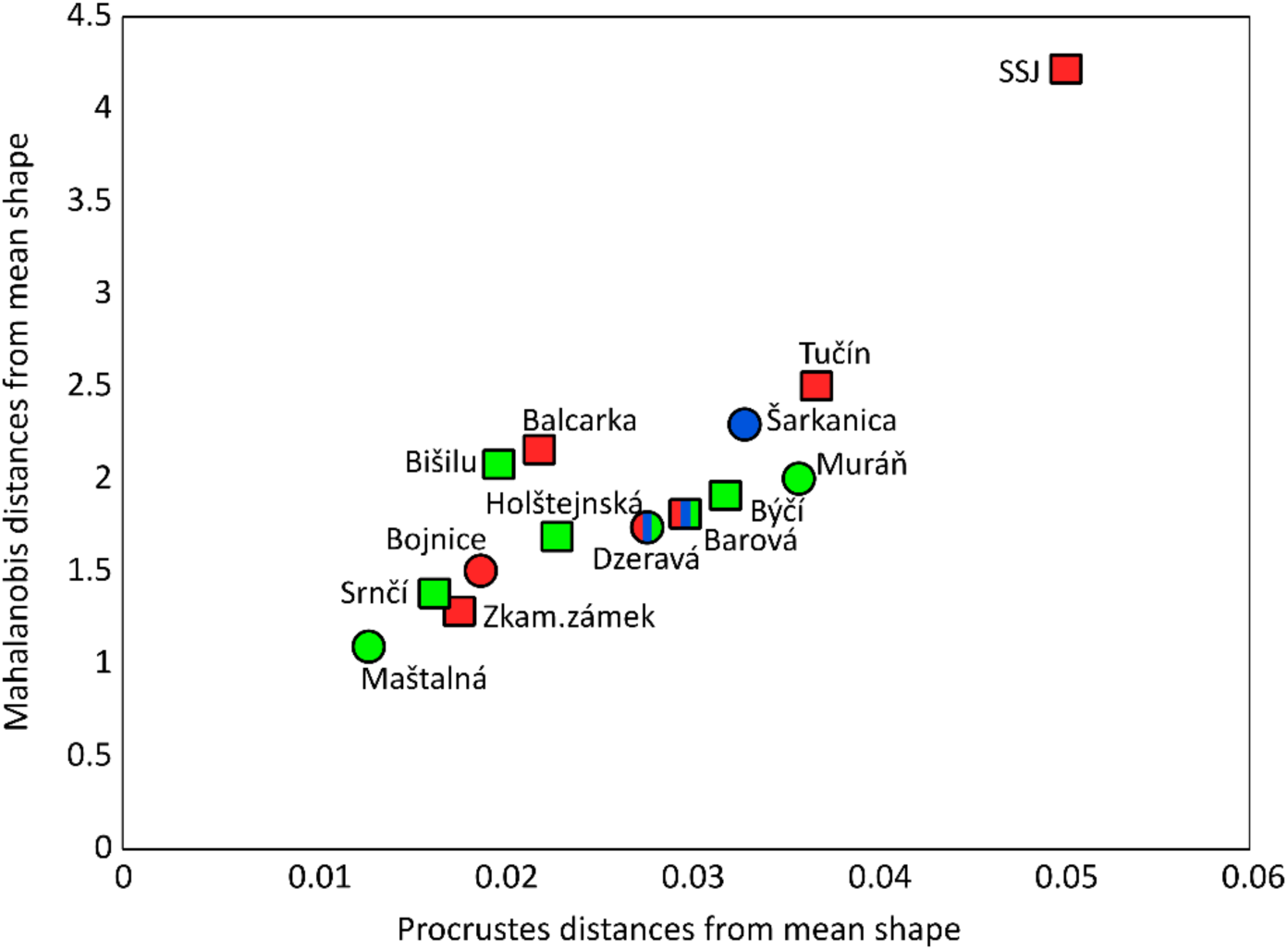
Phenotype variation by geometric morphometrics. Procrustes and Mahalanobis distances of m1 shape in individual local populations from the mean shape pattern based on geometric morphometric data. Squares: Czech localities, circles: Slovak localities. Red color: pre–LGM localities, blue color: LGM localities, green color: post–LGM localities.

Regarding the Procrustes distances, the highest shape divergence was observed at Stránska skála SSJ (0.0510, p < 0.0001), likely representing a primitive Middle Pleistocene (MIS 12) morphotypes. Other notable deviations were found at Tučín (0.0369) and Muráň 3 (0.0354), indicating distinctive morphologies associated with pre–LGM and glacial periods. Šarkanica (0.0327) also exhibited significant divergence, potentially reflecting cold-adapted forms. In contrast, Holocene sites such as Maštalná (0.0133) and Srnčí (0.0156) showed the lowest divergence, suggesting shape convergence toward the dataset mean in post–LGM populations. Srnčí was the only site with non-significant shape differentiation (Procrustes p = 0.236), indicating high similarity to the global mean, however, the number of specimens from this locality was very low (22 individuals), which explains the statistical insignificance. Regionally, Slovak sites (Muráň 3, Šarkanica, Bojnice, Dzeravá skala) tended to show greater shape divergence than Bohemian and Moravian sites, except for Maštalná and Srnčí, which were closest to the mean (Fig. 10). The Mahalanobis distances indicate clear morphological differentiation among the examined localities, with all pairwise comparisons being highly significant (p < 0.0001), except, again, for a case of Srnčí, which shows relatively low differentiation from Bišilu (p = 0.0461), Holštejnská (p = 0.0383), and Maštalná (p = 0.1575). Stránska skála SSJ shows the highest Mahalanobis distances across comparisons, reinforcing its distinctiveness and alignment with early Pleistocene morphotypes. Conversely, the smallest distances are observed among Late Vistulian/ Early Holocene sites such as Maštalná, Srnčí, and Zkamenělý Zámek, indicating greater morphological homogeneity. The distinctiveness of Tučín and Šarkanica further highlights morphological shifts associated with the extreme glacial conditions and eudominant position of *S. anglicus* in mammal communities (84% and 62%).

### (iv) Between-population phenotype variation: common patterns and divergences

As demonstrated above, a particular population might differ significantly in both metric and non-metric traits. While there is extensive overlap in metric variables, the proportions, the frequency of specific morphotypes, morphotype diversity, and ratios of metric variables exhibit clearly pronounced between-population differences, particularly when confronted with the actual dominance of the species in small ground-mammal communities (Figs 6–7). Despite specific differences between the two morphotype sets, the overall pattern of morphotype dynamics is the same: as population abundance declines, morphotype diversity decreases dramatically (Figs 8–9). The other phenomenon worth mentioning here is a significant between-population variation in the sites representing continuous records of local history- Dzeravá skala (53–20 ky) and Maštalná (16–8 ky). The extensive range of variation recorded in these two sites encompasses the states of the respective variables in most other populations. This is also demonstrated by the results of PCA (Figs 6B, 7B), where a distinct outlier position shows either (i) the populations from a deeper Pleistocene past (SSJ, Tučín); (ii) peak glacial stages in the Carpathians (MIS 4 Bojnice, MIS 2 Šarkanica); and (iii) the populations of a terminal stage of the species extinction (Srnčí /4, Holštejnská /4, Maštalná /10, Muráň 3/ 3–4).

A comparison of metric variables (Figs 11–12) revealed an unexpected pattern of between-site divergence. Both in mean differences in metric variation from overall mean values (Fig. 11) and PCA first component of all linear variables (Fig. 12), the total set of samples splits into two distinct clusters, separating closely related samples of Býčí Skála, Barová, and Dzeravá skala from all the remaining samples, as revealed also by a comparison of non-linear variables (comp. Figs 5 and 10).

**Figure 11.**
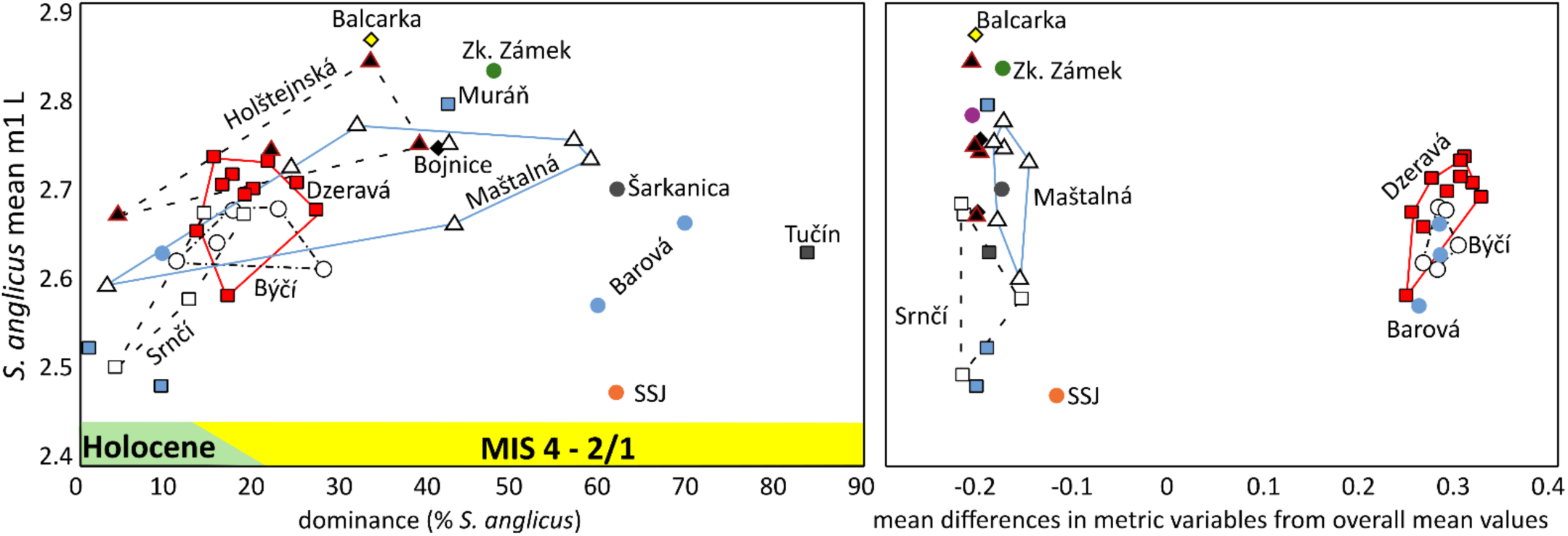
Variation in the main metric variable (mean m1 length). Mean values of m1 length of *S. anglicus* in individual community samples plotted against its dominance (left) and mean differences in all metric variables from overall mean values (diff = (avg((x-avg x)/avg x) - avg all) (right).

**Figure 12.**
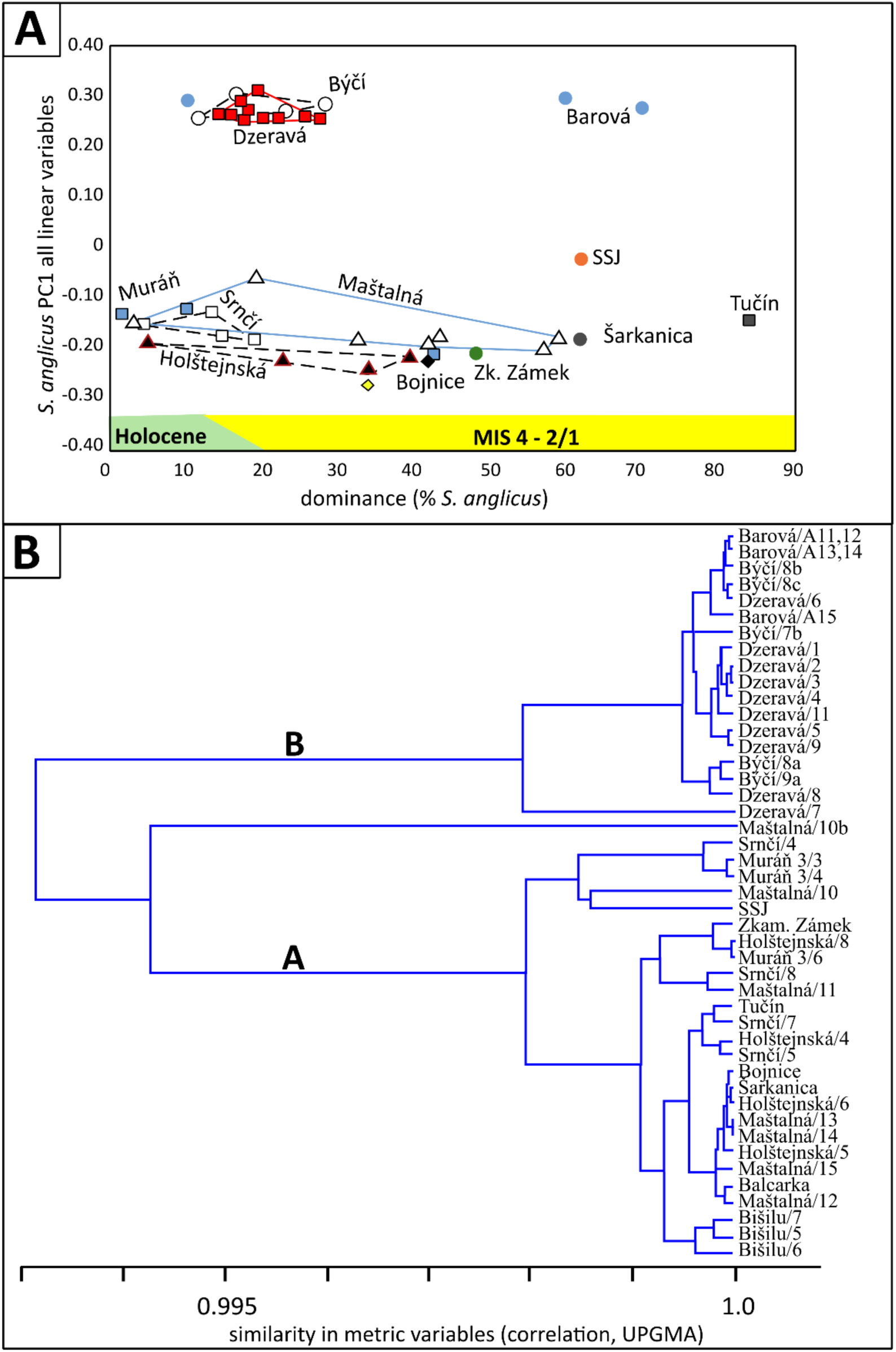
Variation in linear variables and similarities among samples. PC1 values of all linear measurements of *S. anglicus* in individual community samples plotted against its dominance (A) and UPGMA clustering of samples based on correlation in all metric variables (B). Note the high overall similarity within cluster B (Dzeravá-Býčí-Barová) and its distinct differences from other samples (cluster A).

The PCA results of a high-resolution biometric record (155 linear distances among individual landmarks) from particular sites (all layers included), demonstrated in Fig. 13, split the set of sites into two distant clusters: A- the sites of the Carpathian region and those from deeper Quaternary past; C- the sites from the Bohemian Massif (Bohemia + Moravia) with a dense cluster B of Býčí, Barová and Dzeravá situated in between them. The respective PCA results indicate that, for populations in cluster C, the essential source of variation is PC1 (roughly size characteristics); in cluster A, the role of additional variation (shape of the anteroconid complex, proportions) is distinctly more pronounced. In contrast, cluster B seems to show a balanced response to both drivers.

**Figure 13.**
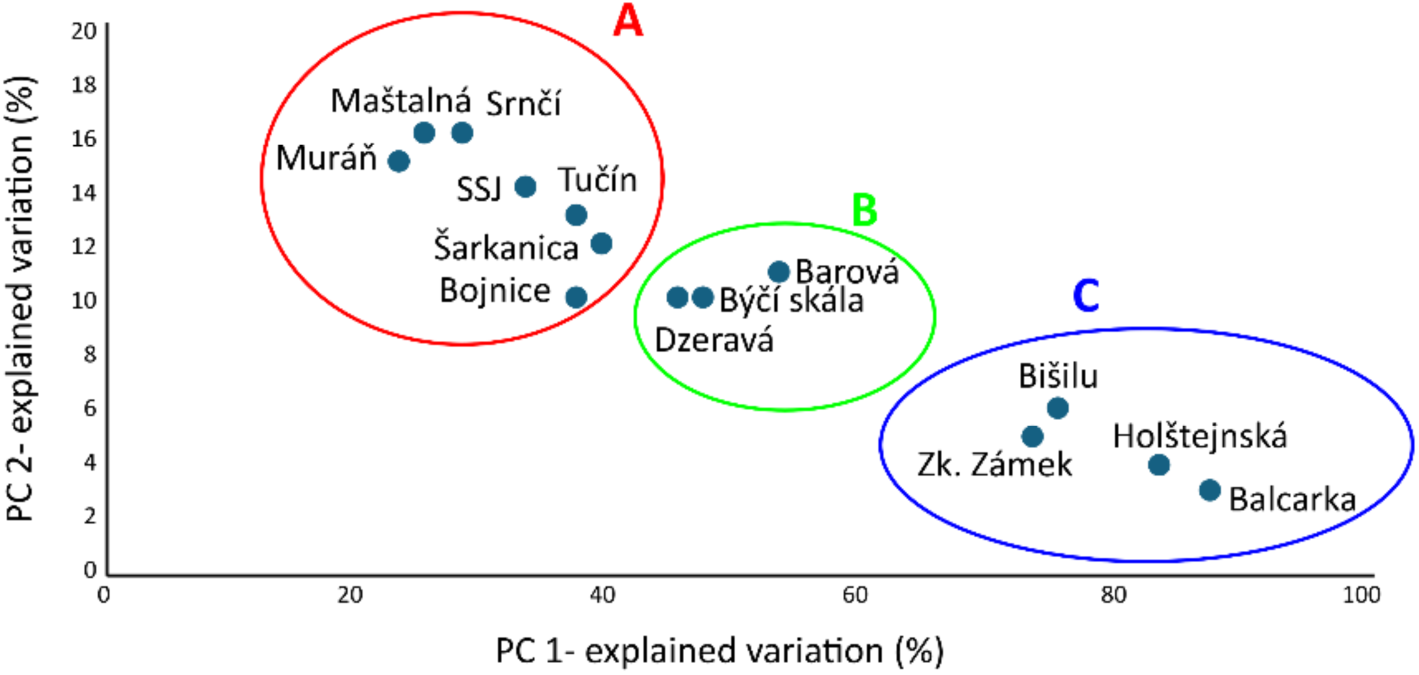
Overall phenotype similarity. Amounts of explained variation by PCA of a high-resolution set of metric variables (155 linear dimensions among individual landmarks) in total samples of individual sites. Note two distant clusters: (**A**) the sites of the Carpathian region and those from the deeper Quaternary past; (**C**) the sites from the Bohemian Massif (Bohemia + Moravia) with a dense cluster of Býčí, Barová, and Dzeravá (**B**) situated in between.

Even more distinct differences between the B cluster sites and all other sites appear in the proportion ratios of metric variables (Figs 14–15), which exhibit values larger than the overall mean of the respective ratio variables and show consistent relations with variation in metric variables (R^2^ = 0.680).

**Figure 14.**
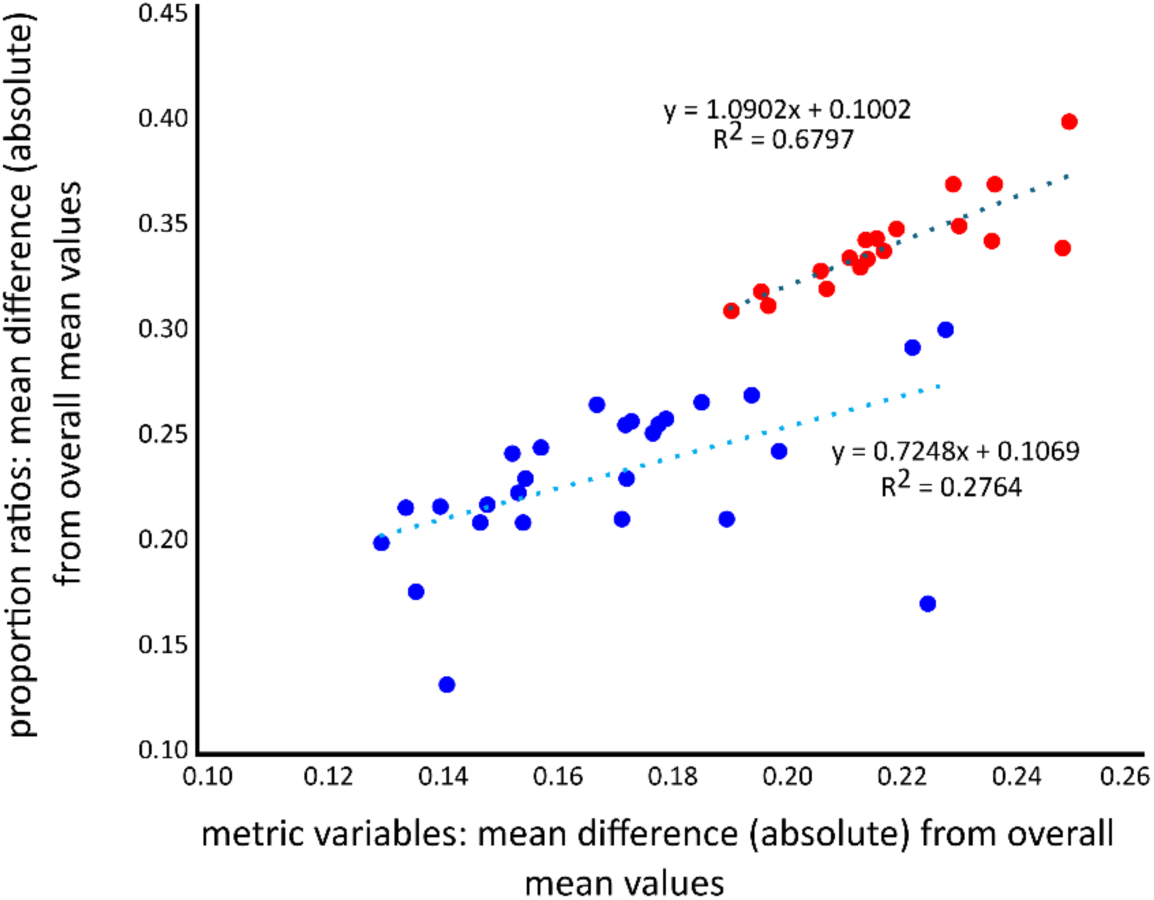
Metric variables against proportion ratios. Plot of mean differences from overall mean values in metric and proportion metric ratio variables in site cluster B (red: Dzeravá, Býčí, Barová) and all other community samples (blue).

**Figure 15.**
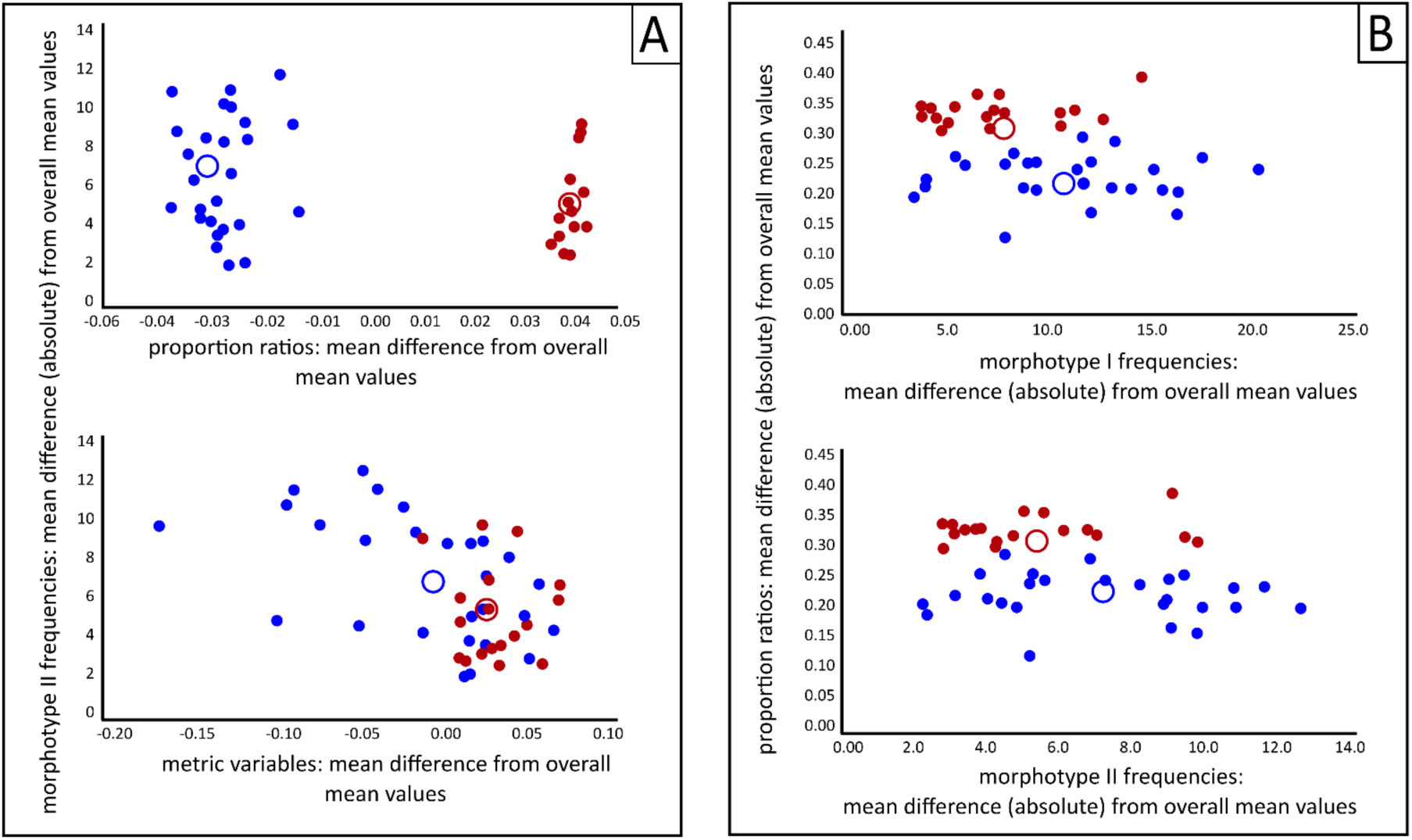
Morphotype I and II frequencies against metric variables and proportion ratios. (**A**) Plot of mean differences from overall mean values in metric and proportion metric ratio variables and morphotype II frequencies in site cluster B (red: Dzeravá, Býčí, Barová) and all the other community samples (blue). (**B**) corresponding plot of mean differences from overall mean values in morphotype I and II frequencies and proportion metric ratio variables. Centroids indicated by hollow circles.

Worth mentioning is that the results of our study concerning the phenotypic divergences among the clades from Czech Massif and Carpathians and MIS 4 sample are robustly supported also by results of aDNA analyses (Fig. 16). The early haplotypes (EF) present in the Carpathian MIS 4 sites (Bojnice, Peskö/ 12) were replaced during MIS 3 by modern haplotypes EB (in the Carpathians) and even EA distributed mainly in Bohemian Massif populations.

**Figure 16.**
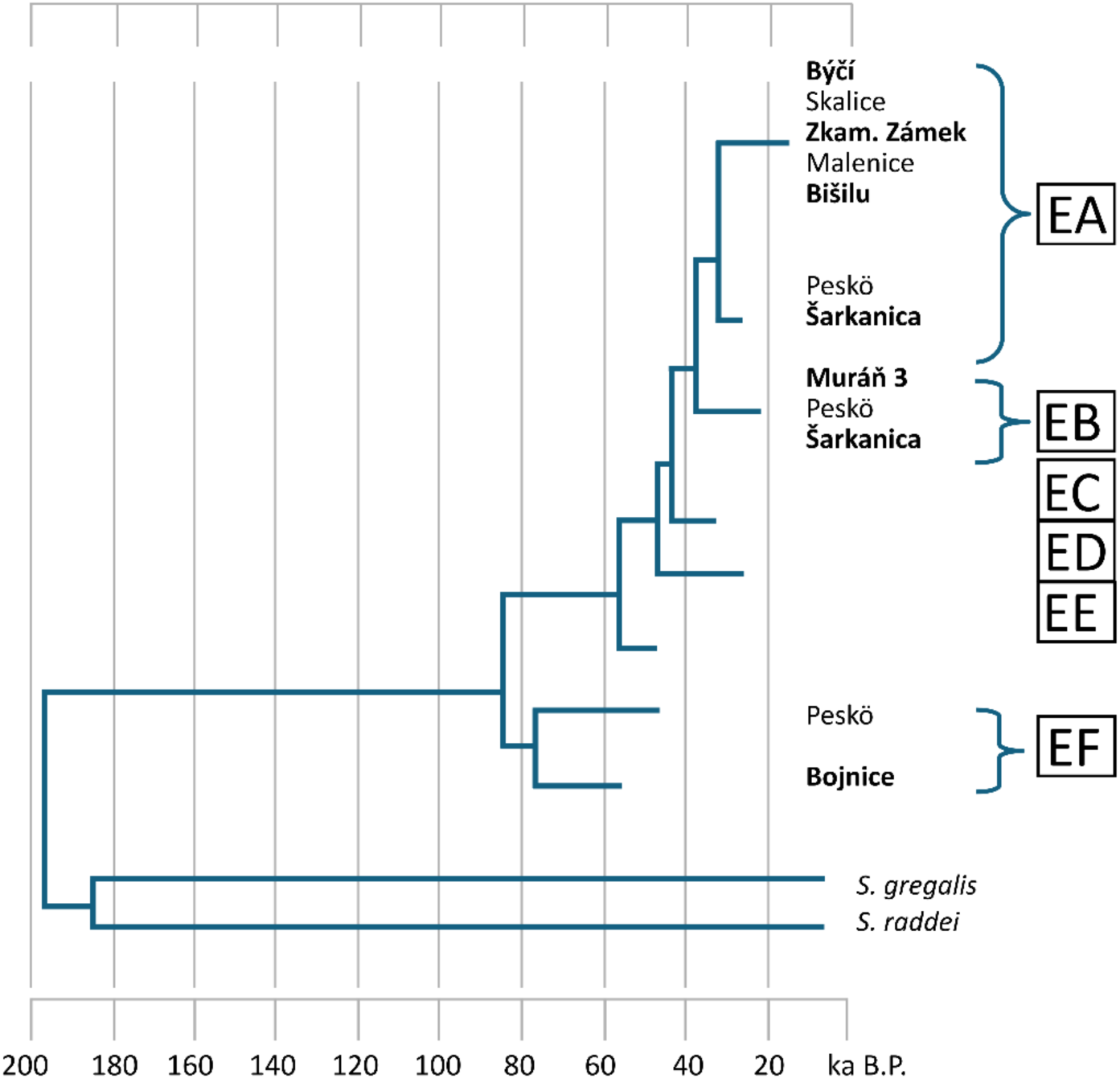
Molecular phylogeny data of the Czech and Slovak samples of *S. anglicus*. Phylogenetic relations among populations of *S. anglicus* from the Czech Republic and Slovakia based on aDNA analyses by Baca et al. (2023a), adapted and simplified. Those studied in the current project are in bold. EA–EF distinct European genotypes after Baca et al. (2023a).

## Discussion

*Stenocranius anglicus,* recently distinguished as a separate European clade from the widespread East-European and Asiatic species *S. gregalis* (Baca et al. 2019), can be similarly distinguished from the other clades of *Microtus* s.l. as its sister species (incl. *Sumeriomys, Terricola, Alexandromys,* etc. - comp.

Kryštufek and Shenbrot 2022) by asymmetric arrangement of the m1 anteroconid complex with a prominent lingual triangle (T7) combined with a rounded labial wall of it (Borodin 2009). Since the first attempts of Quaternary small mammal analyses (Nehring 1875; Woldřich 1881, 1897; Newton 1894; Hinton 1910, etc.), the species, under the name *Microtus gregalis,* was reported from a massive number of sites throughout Europe and identified as an index species of the European glacial communities (Horáček and Ložek 1988). This pattern is also robustly demonstrated by the fossil record from the Czech Republic and Slovakia (235 community samples with MNI of the species 6352), which thus provided a relevant sample for analyses of the species’ dental phenotype variation patterns (not yet reported explicitly for *S. anglicus*) and its history during the Late Pleistocene faunal rearrangements. Results of our study address, in addition to other topics, four topics that will be discussed separately below.

### (i) Phenotype variation of *S. anglicus* over time and space

In general, by its dental phenotype, the Late Pleistocene *S. anglicus* represents a distinctly homogenous unit, a single species, with normal distribution of all studied traits, notwithstanding clearly pronounced variation in both geographic and stratigraphic respects. The patterns of m1 anteroconid shape variation (T6/T7 triangles in particular) mirror findings from Siberian populations of *S. gregalis* sensu stricto (Smirnov 1990; Markova et al. 2009), suggesting Eurasian-wide trends in phenotypic rearrangements, common to both *S. anglicus* and its Asiatic sister clade.

Yet, the Late Pleistocene patterns differ distinctly from those of earlier stages. The earliest specimens in our dataset (Stránska skála cave SSJ, MIS 12 or 14) display a relatively robust, archaic morphology, resembling those of early *S. gregalis* in Kazakhstan (Dupal 1998), suggesting a widespread basal morphotype. Their anteroconid already consists of multiple elements, including distinguishable, though only weakly differentiated, salient and reentrant angles LSA5 and LRA and T6 and T7 triangles, representing an early evolutionary stage of the species, before the emergence of the more derived “arvalid” morphotype common in later populations (Dupal 1998; Serdyuk 2001). More specimens in the SSJ sample exhibit a confluence of T4 and T5 triangles resembling a state in *S. gregaloides* Hinton, 1923 (comp. Kučera et al. 2009). Similar archaic forms are known from Middle Pleistocene sites in Siberia and Kazakhstan (Smirnov 1990), indicating, once again, a broader presence of this basal form across Eurasia. At the locality Tučín (MIS 6), we observe subtle but significant advances in the development of the anteroconid, suggesting trends that terminate in the present cycle. T6 and T7 are more pronounced, and the occlusal outline becomes more complex. The Tučín (MIS 6) advancements align with broader Eurasian trends of increasing molar complexity during cooling phases (Maul and Markova 2007).

The peak of phenotypic diversity and complexity occurred during MIS 5 to MIS 3, as evidenced by sites such as Bojnice, Balcarka, and Zkamenělý Zámek. The common trend covered a teeth robustness, well-developed T6 and T7 triangles, a longer occlusal surface, and stronger asymmetry. An increase in between-site variation in that stage was supposedly driven by local environmental diversification. These features suggest adaptive rearrangements to the conditions of the highly productive mammoth steppe of MIS 3 (Zimov et al. 2012), providing a rich, highly diversified herb diet, yet combined with extreme seasonality in its availability (Pavelková Říčánková et al. 2014) and pronounced competition among herbivorous small mammals. This trend parallels findings in contemporaneous Siberian and Ukrainian assemblages (Serdyuk 2001; Markova et al. 2009; Petrova et al. 2014) as well as those in the fossil record from Ukraine (Serdyuk 2001) and Altai (Agadjanian 2009).

The disappearance of the mammoth steppe during the Last Glacial Maximum due to environmental rearrangements influenced the population density of *S. anglicus* only slightly, presumably also because of population decline or the disappearance of demanding competitors (lagurins, *Microtus arvalis, Phodopus,* etc.). Nevertheless, in *S. anglicus,* we observed a well-pronounced reduction in morphological diversity associated with a simplification of the anteroconid structure, reduced asymmetry, and less pronounced accessory elements (T6, T7, anterior cap, etc.). The frequency of morphotypes in LGM samples also differed clearly from the MIS 3 pattern. We hypothesize that the decline in intraspecific diversity may be associated with a decrease in genotypic diversity. This hypothesis is consistent with the results of comprehensive aDNA analyses by Baca et al. (2019, 2023a), which demonstrated that, from the LGM through the early Holocene, the mid-European region was colonized by only a single remaining haplotype after the disappearance of all others by the end of MIS 3.

### (ii) Phenotypic diversity was scaled by population density

Theoretical analyses (Carmona et al. 2016; Engen and Saether 2019) robustly indicate that functional diversity and trait probability densities increase significantly with species abundance. Mortelliti and Brehm (2020) demonstrated that environmental heterogeneity and population density considerably increase variation in behavioral traits and in the functional richness of adaptive responses in arvicolid rodents. Our analyses seem to support these assumptions with empirical evidence on dental phenotype. Morphotype diversity of *S. anglicus* significantly increases with population density, from 20% to 90% of species dominance. The increase is slow but nearly linear, while below a density of 20%, the phenotype diversity shows a rapid exponential decrease. It can be expected that under conditions of high population density, interdeme selection (Lewontin 1962) increases particularly strongly, eventually becoming a driving mechanism of density-related trends in *S. anglicus* phenotype variation. In any case, our results seem to support models that stress density-dependent factors as essential drivers of the increase in polymorphism (Kim 2025). In these regards, it should be remembered that diversity patterns are not essentially affected by sample size (Hernandez et al. 2006), at least in samples larger than MNI 25 (Cruz-Uribe 1988).

### (iii) Paleobiogeography, local and regional effects upon phenotypic variation

In general, our study indicates that the effects of local and regional variation outweigh those of the stratigraphic position of samples, consistent with conclusions reported for other species (Nadachowski 1982; Borodin 2006; Royer 2016; Luzi and Lopéz-Garcia 2019; Baca et al. 2020, 2024, etc.). Differences in phenotypic characteristics (particularly those revealed by geometric morphometrics) split the total sample into a group of Carpathian populations and those of the Bohemian Massif (both from Moravia and Bohemia proper). Interestingly, the populations older than Vistulian (MIS 12/14 SSJ, MIS 6 Tučín) fall despite their location in Moravia into the Carpathian cluster. This might suggest that the Carpathian populations share some of the common ancestral phenotype of the species, and supposedly, the Carpathians could be considered a source area for glacial expansions of the species during the Vistulian and former glacial stages. The Bohemian clade could then be related to a population that expanded into Central Europe during MIS 3, presumably from the north. Such an idea is also supported by a detailed record of aDNA (Baca et al. 2019 supplementary file Figure S1), where the population from southern Carpathians (Bojnice, Peskö) appears in a basal clade together with those from south Germany and France, while all the others form a separate clade associated with samples from Poland and Ukraine. Relics of the westernmost Carpathian form might then survive most of the Vistulian in isolated microrefugia. Such a possibility is indicated by records from Dzeravá skala, Býčí Skála, and Barová caves, which, by their phenotypic traits, form a compact cluster exhibiting nearly identical variation patterns and, at the same time, differ robustly from all the remaining samples under study.

The samples from diverse layers of Dzeravá skala cave (covering the period from 57.0 to 24.8 ka according to multiple cal ^14^C data - comp. Kaminská et al. 2005) warrant special attention, as they show remarkable morphological stability and consistently high morphotype diversity across all stratigraphic layers. This diversity, coupled with stable proportions of molar traits such as tooth length, width, and anterior characteristics, suggests persistent population continuity and ecological stability, as also apparent from the composition of greatly diversified communities of small ground mammals in that site (Horáček 2006).

A deep valley in the western slopes of the Carpathian Mountains, where the site is situated, provided perhaps a capacity to buffer temporal variations in climatic and environmental conditions affecting the surrounding piedmont and lowland regions, thus establishing the conditions of a long-term refugium, staying apart from extensive disturbances that restricted the narrow-headed vole to maintain an unconstrained phenotypic variation. In these respects, Dzeravá skala’s stability mirrors refugial dynamics in the Balkans (Kryštufek et al. 2020), where topographic complexity buffered climatic shifts. The distinctive phenotypic dynamics in Dzeravá skala and its identity with the characteristics of the populations in Barová and Býčí Skála (both situated in close proximity in the westernmost part of the Moravian karst, resembling the landscape pattern in Dzeravá skala) rank among the most exciting aspects of our study. This suggests the appearance of two mutually isolated (150 km) refugial populations of a single clade not directly related to neighboring populations (e.g., from the eastern part of the Moravian karst). This possibility is also suggested by data on the extinction process, which shows distinct differences between Býčí Skála cave and the sites in the eastern part of the Moravian karst (Srnčí, Holštejnská).

### (iv) Patterns of extinction dynamics of *S. anglicus*

The late Vistulian and Holocene record from the Czech Republic and Slovakia suggests a continuous presence of *S. anglicus* in the Late Vistulian and Preboreal samples across most regions. The dominance of the species, however, rapidly decreased, particularly since the beginning of the Holocene. In Boreal, the species is already missing in most regions, while locally it still survives in Slovakia and Moravia, though at relatively low abundance. Few rare records (mostly single teeth) from samples dated to middle Holocene (southern Moravia: Martinka/2,3,4, Soutěska 2/5; Moravian karst: Zazděná/3,4, Malý Lesík /4, Velká Kobylanka/4,5; Slovakia: Maštalná/3, Peskö/2,3; Central Bohemia: Capuš, Srbsko 1504/f2, Bišilu/3b, Bašta/1, Týnčany/5) indicated a possibility of local survival even to that stage, particularly in southern Moravian areas close to Vienna basin and marginal regions of the Carpathians neighboring the lowland areas of the Carpathian basin.

The late Vistulian and Holocene declines in the species were accompanied by certain rearrangements of the dental phenotype. Compared to the LGM, the phenotypic variation revealed a dramatic shift, marked by an overall reduction in morphotype diversity and a locally specific burst in the skewness and kurtosis of certain traits. The compact and robust molar shape seen during the LGM becomes less pronounced, suggesting a relaxation of the strong selective pressures that had favored this form. Anteroconid morphotypes become less structured and more variable, suggesting a breakdown of the tightly constrained adaptive morphology seen during the LGM. The rapid environmental transformation during the Late Glacial and Early Holocene, with the spread of forests and retreat of open habitats (Horáček and Ložek 1988; Németh et al. 2017), likely disrupted vole population connectivity and led to an extensive range fragmentation. Thus, the patterns of phenotype rearrangements accompanying the population decline seem to show considerable between-site differences.

At sites with stratified sequences, we observe this transition in detail. Morphological data from these localities reveal site–specific trajectories which, in general, illustrate a spectrum of evolutionary responses ranging from long–term stability to gradual simplification with terminal disappearance of the species, accompanied by well–marked between–site differences in phenotypic responses revealed, e.g., by differences in morphotype frequencies. For instance, the last populations of *S. anglicus* in Býčí Skála cave are characterized by vast predominance of the morphotype E, which invariably do not appear or is relatively rare in synchronous populations in the Eastern part of the Moravian karst (Holštejnská, Srnčí), whose phenotype is characterized by dominance of either A, F, D, or G morphotypes, similarly as in other sites (Maštalná, Muráň 3, Bišilu)- see Supplementary file S1 for details. The spikes in skewness and kurtosis of the first lower molar length, which frequently preceded extinction events in our dataset, indicate a struggle by declining populations to adapt to the rapidly changing post–glacial environment.

Besides the climatic and vegetation rearrangements, the increase in abundance of other arvicolid species, synchronous with the decline of *S. anglicus*, observed in most sedimentary sequences, is worth noting. It concerns, namely, *Microtus agrestis*, *M. oeconomus*, and particularly *M. arvalis*, which exhibits an abrupt increase in abundance since the beginning of the Holocene, potentially impacting the appearance of *S. anglicus* through a competitive–exclusion effect, at least in some patches of the environmental mosaic. The last occurrences of *S. anglicus* in Boreal horizons at sites such as Holštejnská and Býčí Skála were synchronous with a marked increase in the abundance of forms demanding bush and forest habitats (*Apodemus* spp., *Clethrionomys glareolus, Glis glis).* For the open–ground elements, expansion of these habitats restricted not only the availability of standard food resources but also dispersal possibilities- the ultimate prerequisite for survival in a variegated environmental mosaic (Peniston et al. 2023). Among rodents, it undoubtedly affected *Microtus arvalis*, *M. agrestis*, and, in particular, *M. oeconomus* and *S. anglicus*, which did not survive the unfavorable conditions. First by multiple local and regional extinctions (as in *M. oeconomus* or *M. agrestis*) or, as in *S. anglicus,* subsequently extended onto all still inhabited range fragments. In any case, similar processes resulted in a large–scale extent of the late Vistulian/Early Holocene range fragmentations and extinctions in many elements of glacial communities, including small mammals, both in Europe (Horáček and Ložek 1988; Stuart 1991; Smirnov et al. 2016; Németh et al. 2017), and abroad (Faith and Surovell 2009; Blois et al. 2010; Brace et al. 2012). This also concerns the Asian range of *S. gregalis*, even its core areas in Altai and Southern Siberia (Petrova et al. 2014, Shi et al. 2021). In general, extinction dynamics and their local variation may reflect an interplay between environmental stressors and the adaptive capacity of a particular clade. Yet, why only *S. anglicus* failed to survive the Holocene in some European refugium (as must have been the case in preceding interglacials- comp. Baca et al. 2019, 2023b) remains an open question.

## Conclusions

Patterns of m1 phenotype variation in a recently distinguished index fossil of European glacial stages, *Stenocranius anglicus*, were analyzed using several morphometric approaches on 2081 individuals from 48 community samples at 14 sites in the Czech Republic and Slovakia.

Most of the sites preserve continuous faunal sequences that document particular stages of late glacial and Holocene history. This enabled us to trace site–specific temporal trends in phenotype variation and compare the particular sites regarding these trends and between–site phenotype differences. Between–site and between–region effects were a more pronounced factor in the overall variation of the species than common temporal trends (except for a decline in variation during the LGM).

Phenotype identity of the local populations from the western part of the Moravian Karst and the Malé Karpaty Mts. (distinctly different from all other populations, including those distributed between them) demonstrated the species’ disposition toward long-term survival in mutually isolated populations. In particular, it accompanied the population decline, synchronous across Central Europe, during the early Holocene, which terminated with the extinction of the species during the Boreal or early middle Holocene. Significant between–site variation in the adaptive responses preceding extinctions (e.g., marked by increases in skewness and kurtosis of certain traits) suggests a wide range of disintegration and mutual isolation of remnant populations at that time. Expansion of woodland habitats is considered the primary driver of the decline in abundance and the eventual extinction of the species.

## Acknowledgements

The authors are obliged to all colleagues who helped with the field excavations of fossil sites and the laboratory treatment of the material. First of all, it concerns our late teachers, namely Vojen Ložek and Oldřich Fejfar, who sparked the senior author’s interest in the topics and laid the basic groundwork for the project. We thank Jano Obuch for providing us with a sample from Šarkanica cave (comp. Obuch 2021), and Stanislav Čermák, Jan Wagner, Vladimír Vohralík, Lutz Maul, Mateusz Baca, and Adam Nadachowski for valuable discussions.

The study was supported by the Specific research project MUNI/A/1576/2024 of the Faculty of Science at Masaryk University in Brno, Czech Republic (ND, MI).

## Notes

### Competing Interest Statement

The authors have declared no competing interest.

