## Supplemental File 1 for "Phenotype diversity and extinction dynamics of the European Narrow-Headed Vole, *Stenocranius anglicus* (Hinton, 1910) (Arvicolinae, Cricetidae, Rodentia), in Central Europe"

**SUPPLEMENTARY FILE S1: SITES UNDER STUDY**

**The sites and morphological variability within individual localities**  **Middle to Late Pleistocene (MIS 12- MIS 5)**

**Stránska skála – SSJ**

*Site characteristics:*
Moravian Karst, 49°11'N 16°40' E, 310 m a.s.l.

A relic of an underground cave passage opened by an artificial gallery in a slope of Stránská Skála hill in Brno, some 200 m NW from the famous sedimentary talus of Stránská skála 1 (Kučera et al. 2009). Details on microfauna collected on several occasions by Horáček et al., confirming the early Middle Pleistocene age of the deposits, were surveyed by Kučera et al. (2009). Locality is mentioned as Stránská skála Cave under number 25 in Horáček and Ložek (1988) and as “Bear Cave at Stránská skála” in Musil (1995).

*Stenocranius anglicus* patterns:

**Stránska skála SSJ** (MIS 12/14, Czech Republic) represents the oldest locality in the dataset and stands out as the most morphologically divergent population of narrow-headed voles analyzed, indicating both the greatest magnitude of shape change and significant statistical separation from other localities. Stránská skála SSJ potentially represents an ancestral stock from which later diversity evolved. Morphometrically, it is characterized by the smallest and narrowest molars in the dataset (mean width = 0.825 mm), with the second smallest molar lengths (min. length = 2.162 mm), and the most extreme AC1/L ratio (0.562) (Table 1). In terms of morphotype diversity, the Stránská skála SSJ population is consistent with an ancestral, less variable morphotype spectrum (Fig. A1). The molars were dominated by ancestral morphotypes defined by convex or plain buccal sides of the anterior cap and minimal development of the BRA4 reentrant angle. These traits align with simpler morphotypes A–C (per Nadachowski 1982), supporting the inference of a more primitive evolutionary stage. The relatively lower diversity and distinctive morphology observed here reinforce Stránská skála SSJ´s importance as a baseline for understanding morphological evolution in narrow-headed voles. Despite this distinctiveness, classification accuracy in CVA (58%) suggests that, while morphologically unique, there is partial overlap with later populations, possibly due to the retention of ancestral traits over time.

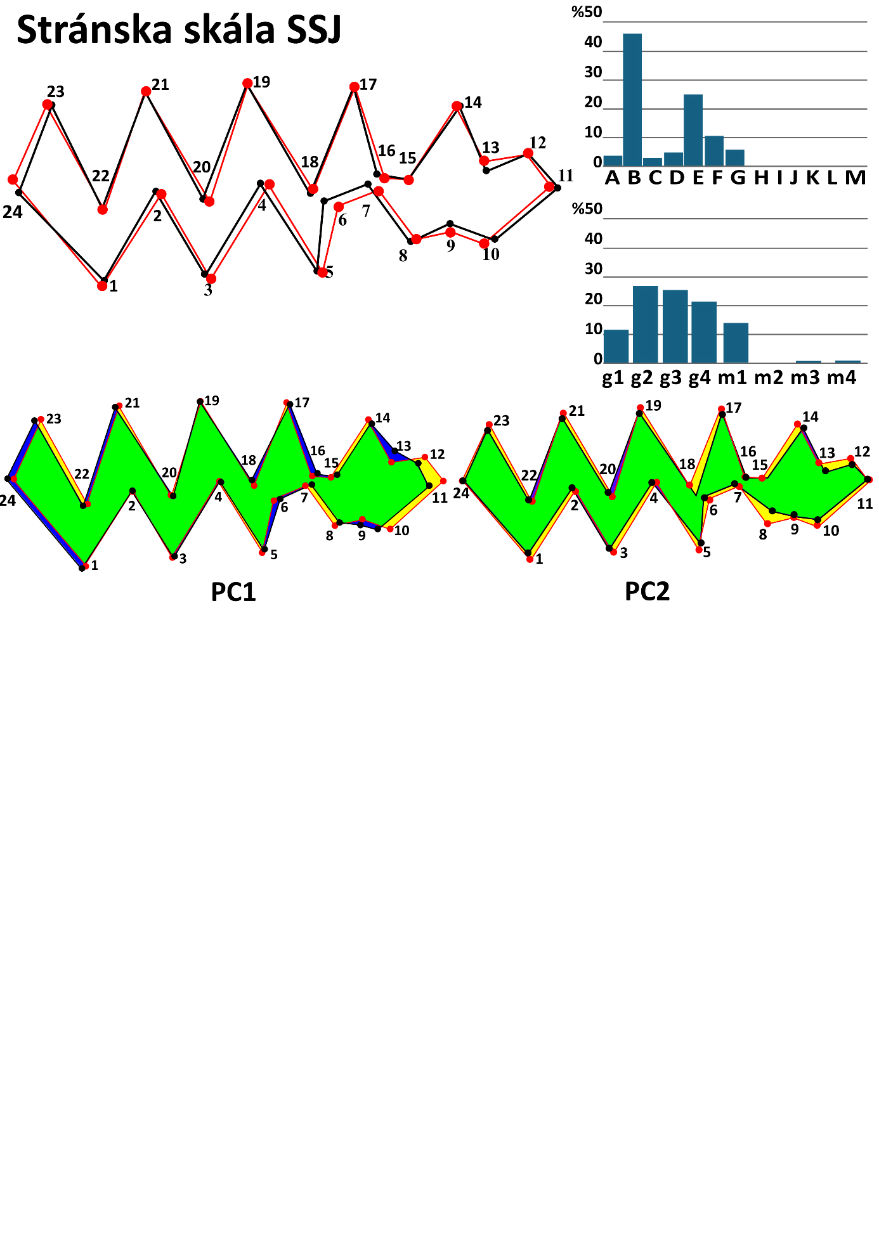

Fig. A1. Shape change and morphotype frequencies at the locality Stránska skála SSJ.
Mean m1 shape of the population of Stránská skála SSJ (red outline) against the overall mean shape (black outline); morphotype frequencies A–M per Nadachowski 1982; g1–m4 BRA4–based (after Smirnov et al. 1986; Ponomarev and Puzachenko 2017);
shape change of the narrow-headed voles' first lower molar in the first and second principal components (PC).

Table 1. Metric characteristics of the population of Stránska skála SSJ.

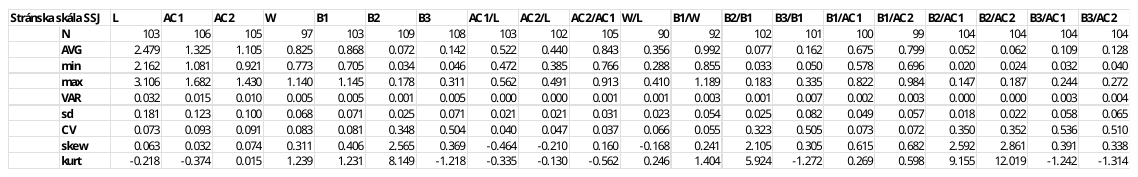

**Tučín**

*Site characteristics:*

North Moravia, 49°25'N 17°27'E; 200 m a.s.l.

The profile is approximately 200 m high. It represents a loamy filling of a cavern in Upper Pleistocene travertine, originating at a spring with CO2 effusion (Kovanda 1971). Fejfar collected a very rich microfauna at this locality in 1959-1961, which was later surveyed by Horáček and Sánchez-Marco 1984). Detailed stratigraphic research on the travertine, including the study of the mollusk fauna at the locality, is presented in Ložek and Tyráček (1958). They reported the fauna as Vistulian, yet both the extent of the travertine body and numerous faunal differences from Vistulian sites suggest an older age. Regarding the count of top stratigraphy, we tentatively place it in MIS 6.

*Stenocranius anglicus* patterns:

**Tučín** (MIS 6, Czech Republic) represents a key pre–Vistulian locality exhibiting notable morphological distinctiveness in the first lower molar (Fig. A2). Tučín’s population demonstrates substantial deviation from the overall dataset mean, suggesting marked morphological shifts likely driven by glacial environmental pressures during MIS 6. Molars from Tučín are smaller than those from Stránska skála SSJ and exhibit the lowest AC1/L ratio (0.503), indicating relatively shorter anteroconid proportions (Table 2). Notably, extreme skewness and kurtosis in B2 and B3 proportions suggest non-normal distributions, potentially reflecting localized evolutionary pressures or taphonomic biases unique to this population. Diversity levels at Tučín remained moderate to high, suggesting increased morphological complexity relative to the earlier period (Fig. A2). These values indicate a stable and possibly transitional morphological diversity during the pre–LGM cold phase, with early emergence of gregaloid-microtid morphotypes, hinting at adaptive responses to changing climatic conditions. The highest correct classification rate in CVA (82%) reinforces populations´ morphological distinctiveness and suggests limited overlap with other populations (Appendix II). Collectively, Tučín illustrates a critical evolutionary phase characterized by increasing morphological complexity and differentiation.

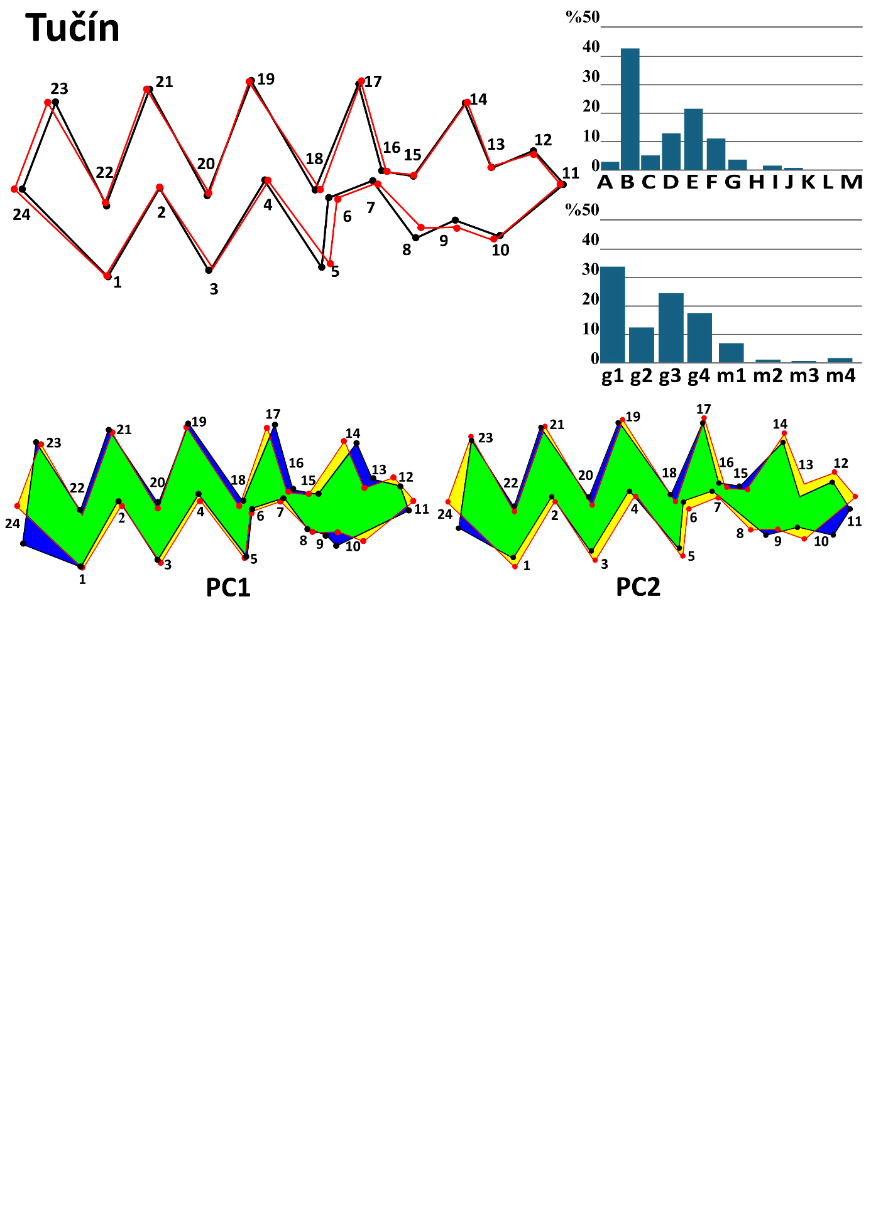

Fig. A2 Shape change and morphotype frequencies at the locality Tučín.
Mean m1 shape of the population of Tučín (red outline) against the overall mean shape (black outline); morphotype frequencies A–M per Nadachowski 1982; g1–m4 BRA4–based (after Smirnov et al. 1986; Ponomarev and Puzachenko 2017); shape change of the narrow-headed voles' first lower molar in the first and second principal components (PC).

Table 2. Metric characteristics of the population of Tučín.

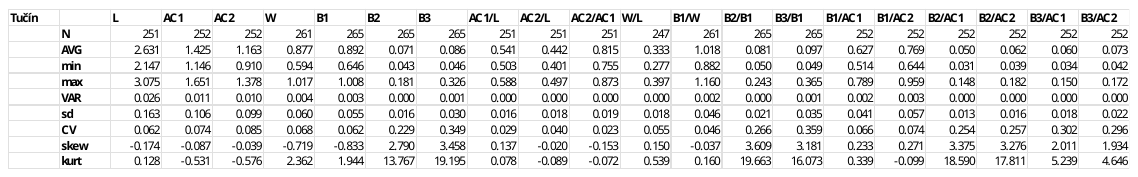

**Bojnice (Prepoštská cave)**

*Site characteristics:*
Southwestern Slovakia, 48°48'N 18°35'E; 276 m a.s.l.

Bojnice 65- Prepoštská cave (MIS 4, Slovakia) is situated close to the Bojnice Castle and represents one of two Neanderthal settlements in the vicinity of the Bojnice Castle in the Horná Nitra region. The cave represents a spacious cavity with approximately 8 m deep, 11 m wide, and 4–8 m high internal volume eroded in a body of Eemian travertine cascade deposited by a local thermal spring (Prošek 1952). A rich mammalian microfauna obtained from revision sections by Prošek and Ložek (1950-1964) was analyzed by Horáček and Sánchez-Marco (1984). In total, it includes remains of at least 694 individuals (MNI) of 16 spp. with predominating *Microtus arvalis* (357) and *Stenocranius anglicus* (250), the other glacial elements are quite rare. In agreement with lithological, archeological, and malacological inferences concerning the site, they proposed a biostratigraphic dating to the early Vistulian (supposedly MIS 4). Radiocarbon dating (three records spanning from cal.^14^C 55224 to 65206) supports that interpretation.

*Stenocranius anglicus* patterns:

**Bojnice 65** represents a pivotal transitional stage in the evolution of the narrow-headed vole, characterized by high morphological diversity and increasingly robust tooth morphology. It exhibited the second-largest average molar length in the dataset and the widest measured molar length, confirming the presence of larger, more robust molars at this locality. Among proportional characteristics, the Bojnice population also displayed high relative values for the anteroconid complex (AC1 and AC2), comparable to suggesting prominent anterior tooth structures (Table 3). Diversity analyses further highlight Bojnice’s distinctiveness, indicating an increase in both morphological complexity and variability (Fig. A3). These values mark the Bojnice population as a peak in diversity during the Middle to Late Pleistocene, potentially linked to an environmental dynamic that promoted morphotype diversification, especially gregaloid-microtid and early microtid forms (Fig. A3). In shape analysis, moderate morphological divergence from the dataset mean likely reflects a transitional morphology. However, the lowest classification accuracy at CVA (24.1%) (Appendix II), with specimens frequently misclassified as Šarkanica, suggests substantial morphological overlap or convergence with populations of the LGM. Overall, Bojnice marks a key evolutionary inflection point, with larger, more complex molars emerging alongside elevated diversity.

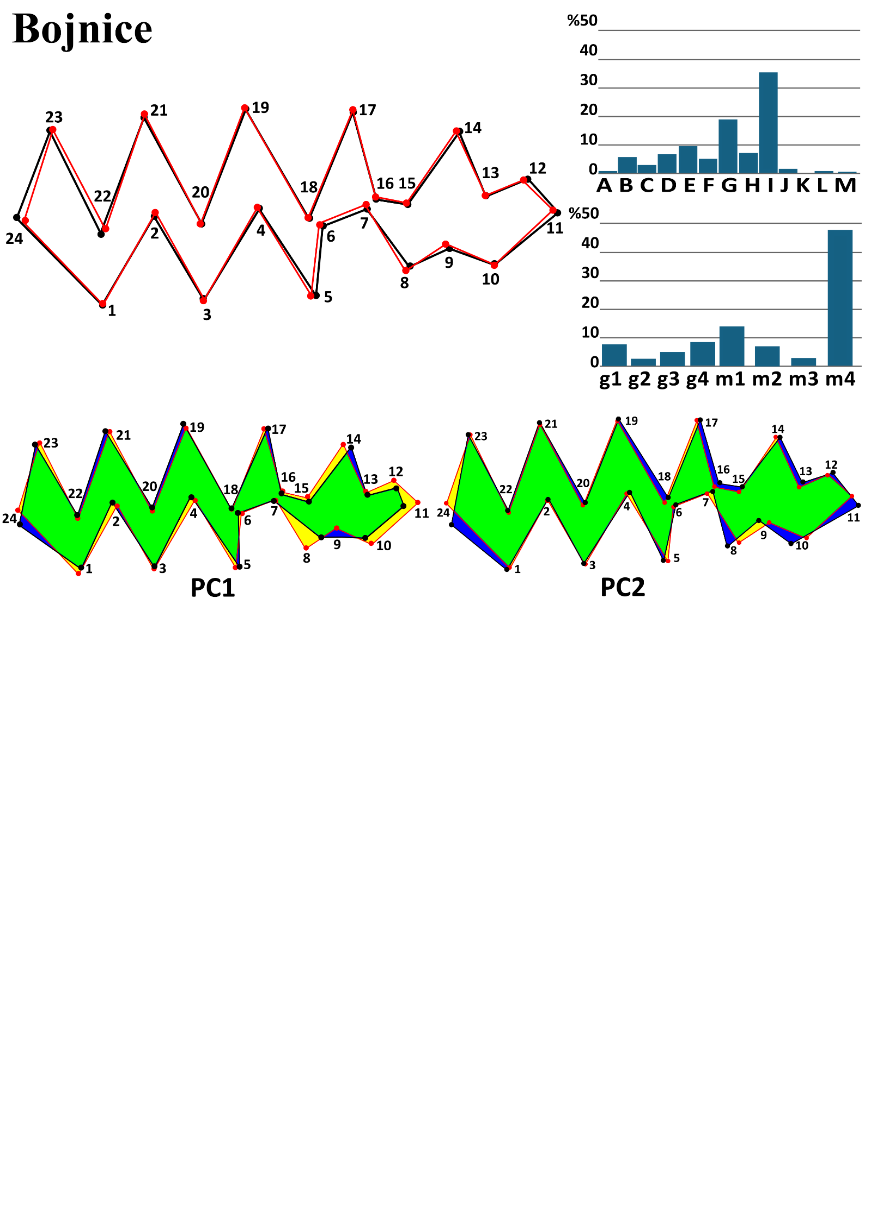

Fig. A3 Shape change and morphotype frequencies at the locality Bojnice.
Mean m1 shape of the population of Bojnice (red outline) against the overall mean shape (black outline); morphotype frequencies A–M per Nadachowski 1982; g1–m4 BRA4–based (after Smirnov et al. 1986; Ponomarev and Puzachenko 2017); shape change of the narrow-headed voles' first lower molar in the first and second principal components (PC).

Table 3. Metric characteristics of the population of Bojnice.

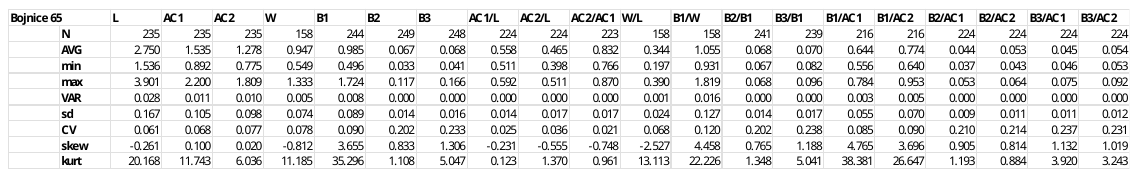

**Late Pleistocene to Last Glacial Maximum (LGM)**

**Balcarka**

*Site characteristics:*
Moravian karst, 49°22'N 16°45'E; 450 m a.s.l.

Balcarka cave is situated in a karst valley near Ostrov u Macochy. The underground labyrinth of passages, fissures, and caves has two floors. Balcarka was formed in tectonically broken, soluble limestones by the gradual erosion and corrosion caused by rainwater. It lies close to the border of limestone and non-karst rocks and functioned as a sinkhole. Interdisciplinary research on the Balcarka cave, its origin, and development can be found in Nývltová Fišáková et al. 2011. The studied sample was excavated by I. Horáček from a sedimentary talus below a chimney near the entrance corridor.

*Stenocranius anglicus* patterns:

**Balcarka** (MIS 3, Czech Republic) represents a morphologically diverse and robust population of narrow-headed voles during the pre–LGM glacial period. The molar dimensions at Balcarka rank among the largest and most robust in the dataset, mirroring patterns seen in other MIS 3 localities (Table 4). Balcarka exhibited the highest morphological diversity recorded pre–LGM (Fig. A4). This extensive morphotype variation is likely associated with advanced microtid evolution and may point to increased environmental variability or regional microhabitat specialization. Shape analysis indicated moderate shape divergence from the dataset mean, but notable statistical separation from other localities. Despite the modest Procrustes divergence, the high Mahalanobis distance suggests strong internal morphological differentiation. High classification accuracy (63.2%) in the CVA (Appendix II) reinforces the distinctiveness and cohesion of its morphometric profile.

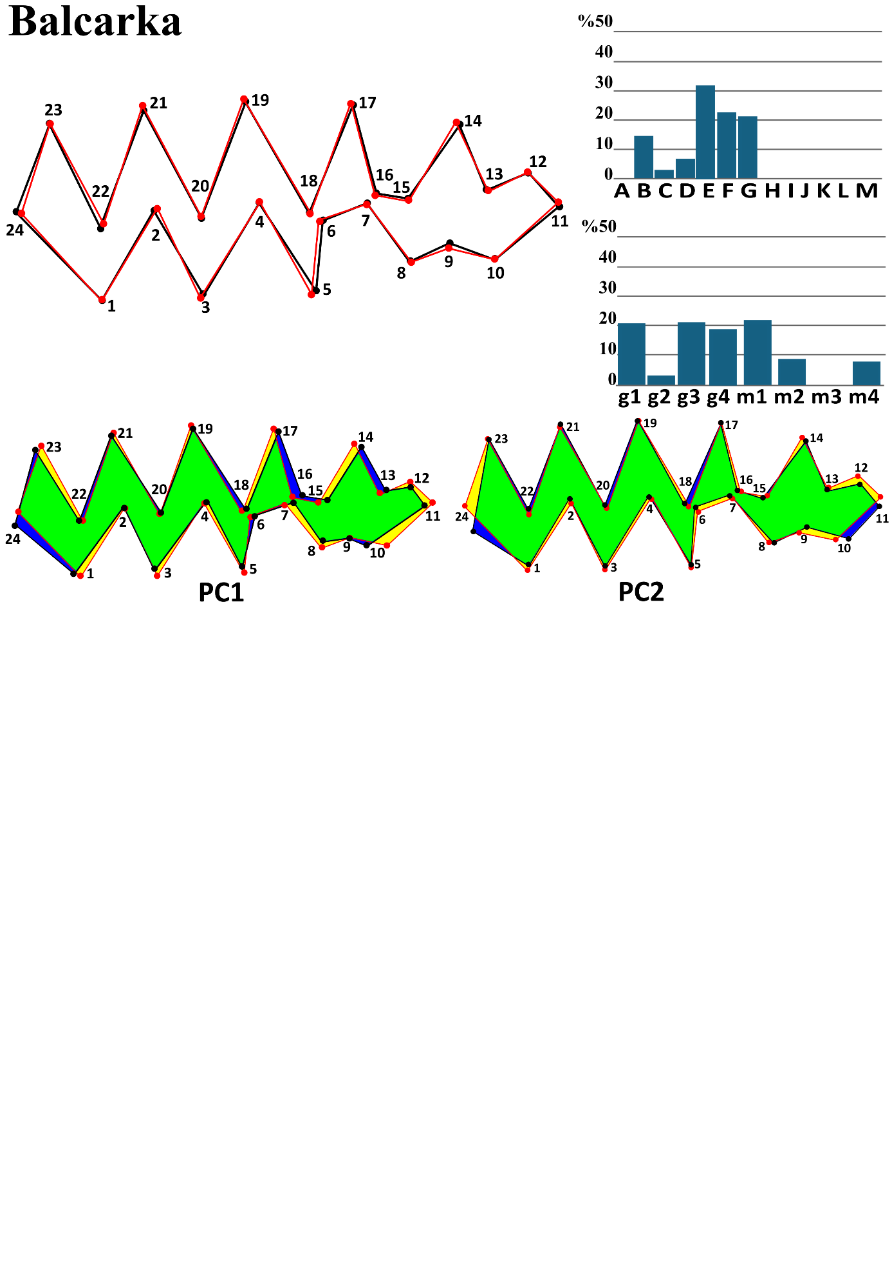

Fig. A4 Shape change and morphotype frequencies at the locality Balcarka.
Mean m1 shape of the population of Balcarka (red outline) against the overall mean shape (black outline); morphotype frequencies A–M per Nadachowski 1982; g1–m4 BRA4–based (after Smirnov et al. 1986; Ponomarev and Puzachenko 2017); shape change of the narrow-headed voles' first lower molar in the first and second principal components (PC).

Table 4. Metric characteristics of the population of Balcarka

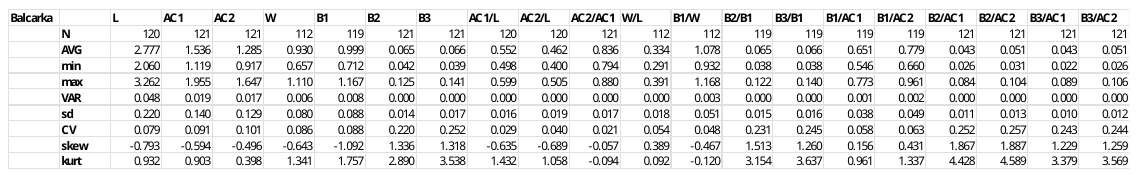

**Zkamenělý Zámek**

*Site characteristics:*
N Moravia, 49°40'N 16°56'E; 768 m a.s.l.

Zkamenělý Zámek is located between Vojtěchov and the Javoříčko caves at the southwestern slope of the Borovský forest in the mountain range Žďárske Vrchy. Material excavated during several years from the bottom, darker sandy loam sediments with approximately 15 cm thick bone layer at a deep part of the cave (the first samples surveyed by Horáček and Sánchez-Marco 1984).

*Stenocranius anglicus* patterns:

**Zkamenělý Zámek** (MIS 3, Czech Republic) represents the most morphologically diverse population of narrow-headed voles in the entire dataset, indicating a peak of diversification during the late Pleistocene. This locality showed the highest morphotype diversity among all sampled localities, surpassing even most Holocene values. Such rich, even morphological variation implies an ecologically complex or mixed environment, possibly with stable, favorable local conditions that support a wide range of morphotypes. Morphometrically, Zkamenělý Zámek's molars were larger and more robust, with notably high values in the anteroconid complex (AC1 and AC2). It had the lowest AC2/AC1 ratio (0.775), indicating a relatively short AC2, and the smallest relative basin width (B1/W = 0.944), which contributed to its unique shape profile (Table 5). Despite this distinct internal variation, on average, the mean shape was similar to the overall dataset mean (Fig. A5). Lower Mahalanobis distances and the CVA classification accuracy (40.9%) (Appendix II) also support the idea that, while morphological diversity within the locality was high, the average shape aligned closely with general patterns across localities.

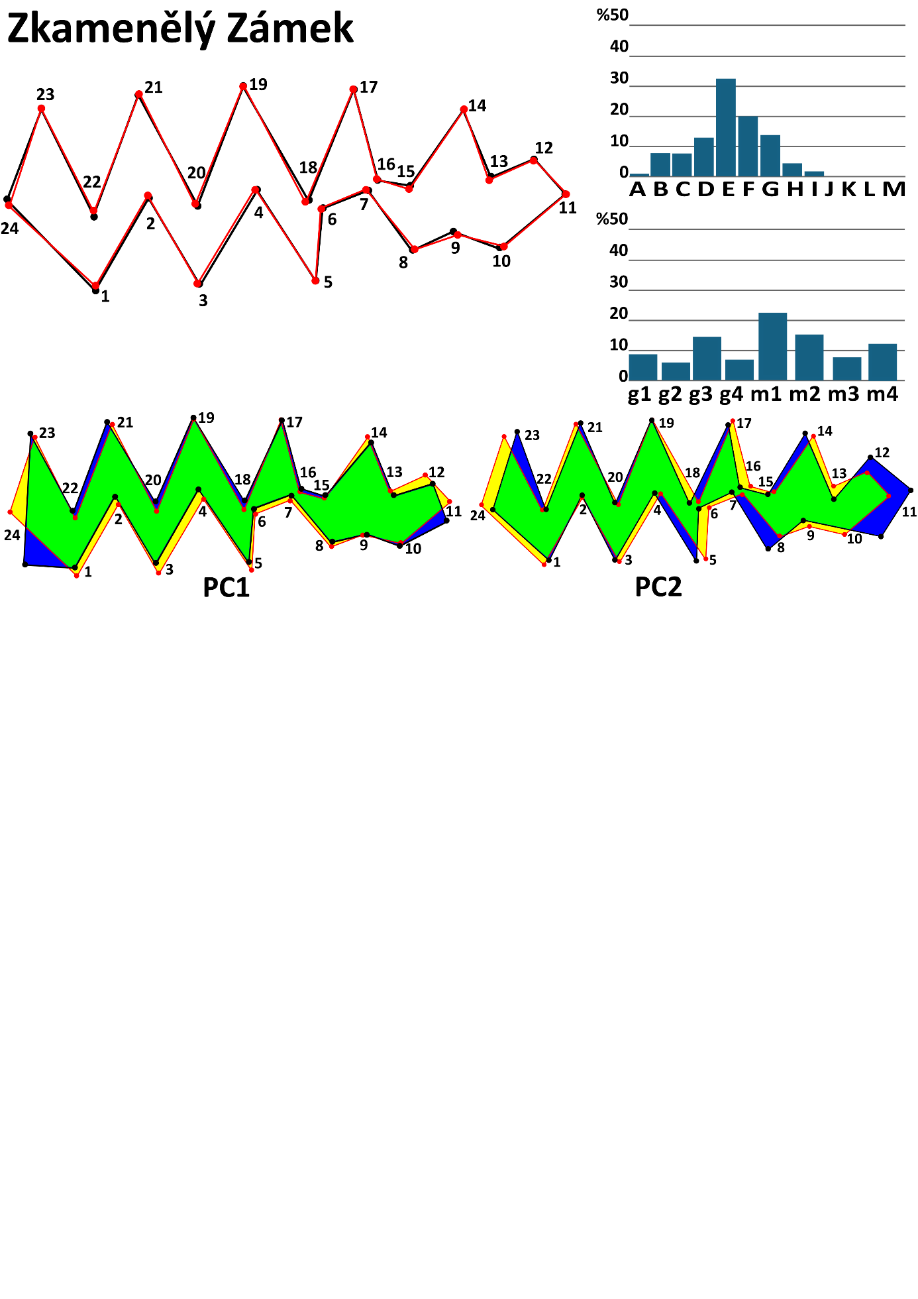

Fig. A5 Shape change and morphotype frequencies at the locality Zkamenělý Zámek.
Mean m1 shape of the population of Zkamenělý Zámek (red outline) against the overall mean shape (black outline); morphotype frequencies A–M per Nadachowski 1982; g1–m4 BRA4–based (after Smirnov et al. 1986; Ponomarev and Puzachenko 2017); shape change of the narrow-headed voles' first lower molar in the first and second principal components (PC).

Table 5. Metric characteristics of the population of Zkamenělý Zámek

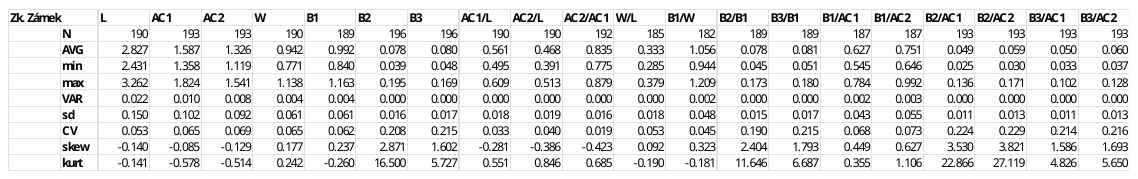

**Šarkanica**

*Site characteristics:*
Slovakia, 48°42'N 19°58'E; 925 m a.s.l.

Cave Šarkanica is in eastern Slovakia, near Tisovec, on the Muráň Plateau. The fossil fauna found at Šarkanica was probably protected during the coldest periods of the Late Pleistocene by warmer, sheltered places in the Muránka River Valley. Material representing former snowy owl nests contained fossil material in three layers, radiocarbon-dated to 17 970–21 000 years BP (Obuch 2021). Yet, three other datings (spanning from cal.^14^C 20 919 to 21 080) suggest an older age around the beginning of the LGM. The material reported in this study was provided by Jan Obuch, who collected the samples and reported the fauna (Obuch 2021).

*Stenocranius anglicus* patterns:

**Šarkanica** (MIS 2, Slovakia) offers a compelling example of morphological stabilization under glacial–steppe conditions, which align with the ecological optimum of the narrow-headed vole. The average molar length at Šarkanica (L = 2.698 mm) remained comparable to that of pre–LGM localities, suggesting that significant size reductions likely occurred post–LGM during the Holocene. Molar width was similarly stable (W = 0.933 mm), but certain proportional traits exhibited notable variability. Specifically, high coefficients of variation for B2/B1 and B3/B1 ratios, along with extreme skewness and kurtosis in B2, suggest asymmetrical trait distributions, potentially due to adaptive flexibility or population–level heterogeneity (Table 6). These long–tailed distributions point to a core of stable morphologies alongside occasional extreme forms, likely reflecting localized ecological pressures or subpopulation differences. Morphologically, the population displayed moderate to high diversity consistent with population stability and ecological specialization. From a shape analysis perspective, the population of Šarkanica showed significant divergence from the dataset mean (Fig. A6). The classification accuracy in CVA (64.2%) highlights a strong group distinction, confirming Šarkanica’s morphological uniqueness while maintaining some overlap with other populations. In summary, the population exemplifies morphological stability and moderate diversity during the LGM, reflecting a specialized, well–adapted vole population thriving in harsh glacial–steppe ecosystems.

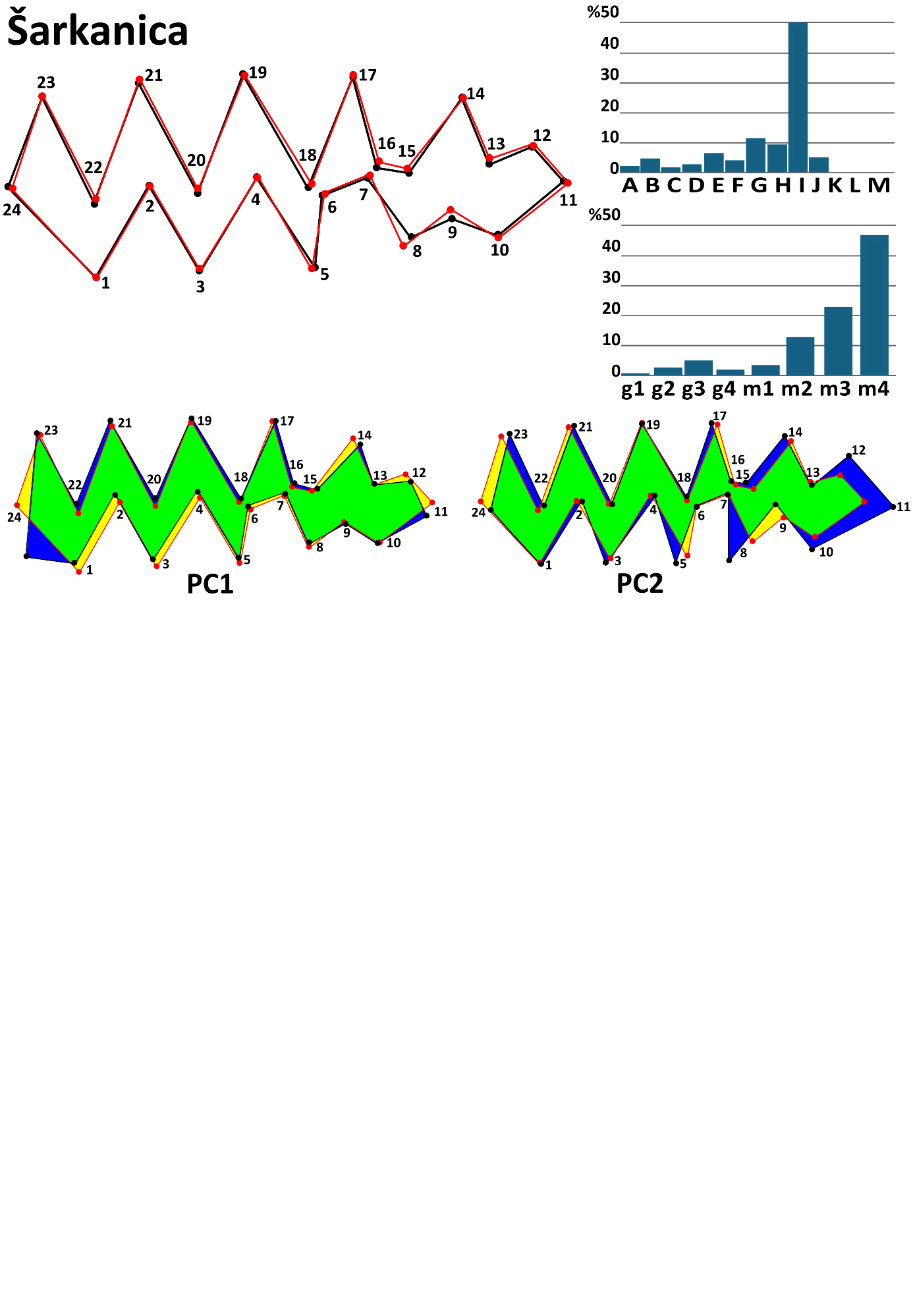

Fig. A6 Shape change and morphotype frequencies at the locality Šarkanica.
Mean m1 shape of the population of Šarkanica (red outline) against the overall mean shape (black outline); morphotype frequencies A–M per Nadachowski 1982; g1–m4 BRA4–based (after Smirnov et al. 1986; Ponomarev and Puzachenko 2017); shape change of the narrow-headed voles' first lower molar in the first and second principal components (PC).

Table 6. Metric characteristics of the population of Šarkanica

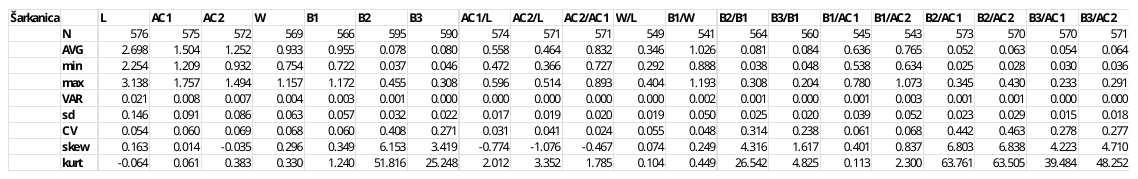

**Continuous Stratigraphical Sequences**

**Muráň 3**

*Site characteristics:*
Slovakia, 48°45'N 20°3'E; 1559 m a.s.l.

The third log of Muráň cave on the southern wall of the mountain Muráň (1827 m a.s.l.), excavated in 1994 by Horáček et al., represents a unique mountainous small mammal assemblage. The stratigraphic sequence spans the period from the LGM to the Middle Holocene. It contains five clearly distinct stratigraphic layers of boulders filled with frost debris and silty matrix. The ages of the base layers were determined by radiocarbon dating to 24 368 ±335 (layer 6) and 16 314 ±427 (layer 4) years BP (Horáček 2015). The third layer represents the late glacial period, and the top layer belongs to the early Holocene. The further cal. 14C data show the following mean ages (layer 6: 24346, 24254; layer 4: 14298; layer 3: 12308, 12395; layer 1: 11953, 8711). Glacial species dominate the small mammal assemblage of the site, however, with the appearance of insectivores, species typical of the mosaic environment of humid open landscapes and wetlands, and even typical forest species (Horáček 2015; Horáček et al. 2015).

*Stenocranius anglicus* patterns:

The **Muráň** locality provides a detailed stratigraphic record of morphological variability in the first lower molars of narrow-headed voles across the Late Glacial Maximum (LGM) and into the post–LGM transition, preceding the species’ local extinction. Three distinct layers highlight key trends: Layer 6 (24.4ky BP) exhibited the highest morphotype diversity (H′ = 1.923) that declined in Layer 4 (16.3–14.2ky BP; H′ = 1.230) and remained low in Layer 3 (12.3–13.4ky BP), suggesting population contractions toward the end of the LGM. Metric characteristics indicate a reduction in molar length from 2.790 mm (Layer 6) to 2.486 mm (Layer 4), followed by a minor recovery (2.527 mm) in the post-LGM phase (Layer 3). Despite these shifts in size, molar proportions remained relatively stable, with W/L and AC1/L ratios showing only minimal variation, suggesting conserved tooth shape across fluctuating environments (Table 7). However, variability measures reveal subtle but meaningful changes: standard deviation remained stable (~0.15–0.18), but skewness became progressively less negative, indicating a shift from left-skewed to more symmetric distributions as extinction approached. Layer 6 displayed extreme skewness and kurtosis for B1, reflecting asymmetric distributions and outliers, which later stabilized, potentially signaling population homogenization. Additionally, the coefficient of variation (CV) for proportional traits peaked in Layer 4, highlighting increased relative variability at the LGM peak. Morphometric analyses suggest that the population was morphologically distinct, reinforcing the idea of localized adaptation and differentiation. The population of Muráň retained relatively high diversity and morphological stability, suggesting its potential as a glacial refugium (Fig. A7). However, the eventual decline in diversity and morphological plasticity signals the final phase of population contraction, culminating in extinction. These trends collectively point to a gradual morphological stabilization rather than abrupt disruption, suggesting that narrow-headed voles in Muráň exhibited resilience under LGM conditions but ultimately succumbed to post–glacial environmental shifts.

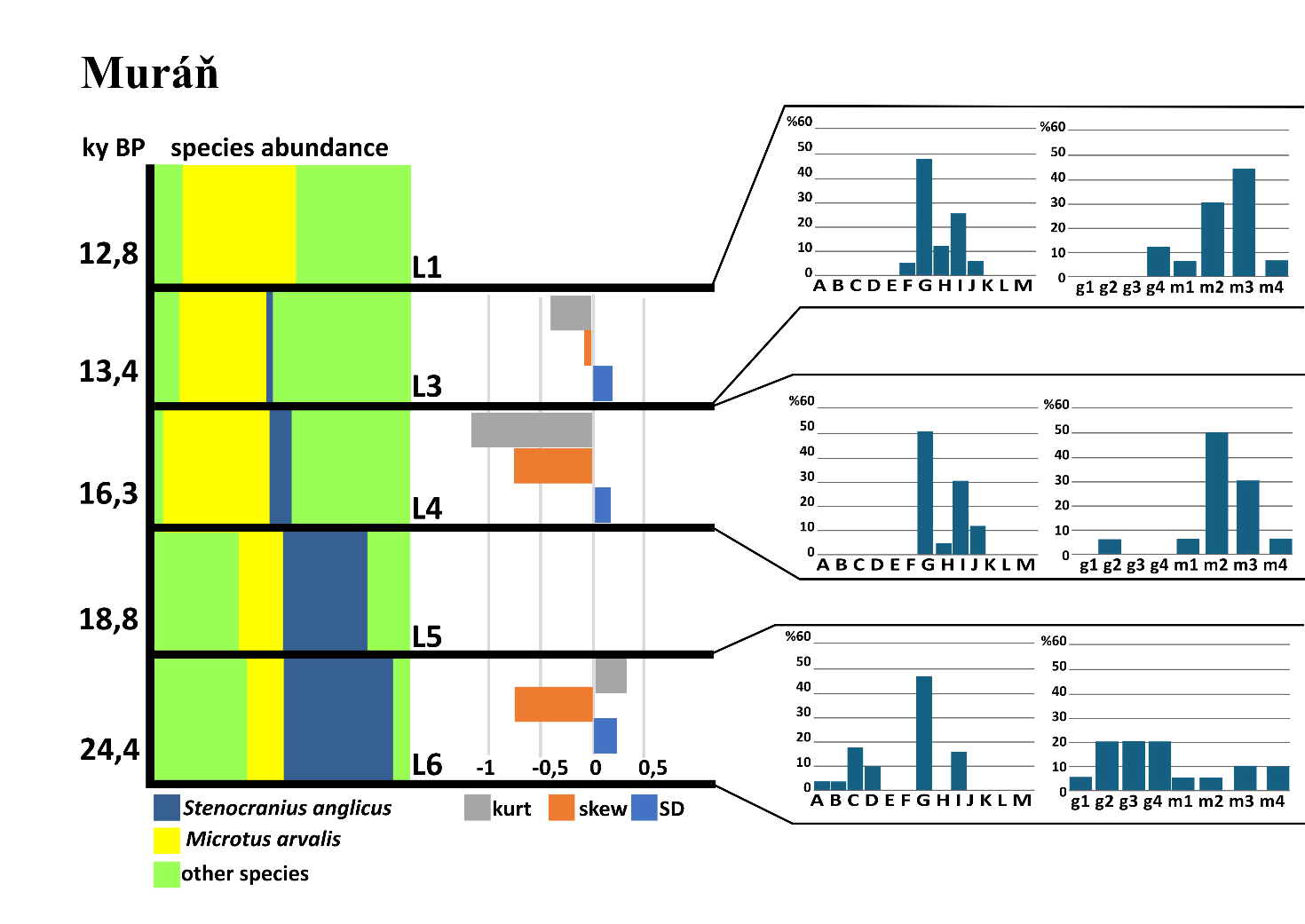

Figure A7. Characteristics of the narrow-headed voles at the locality Muráň.
Species distribution, skewness, and kurtosis of m1 length and morphotype distribution (per Nadachowski 1982, BRA4 angle–based) of individual layers (L) at particular sedimentary sequences of locality Muráň.

Table 7. Metric characteristics of individual layers in the population of Muráň

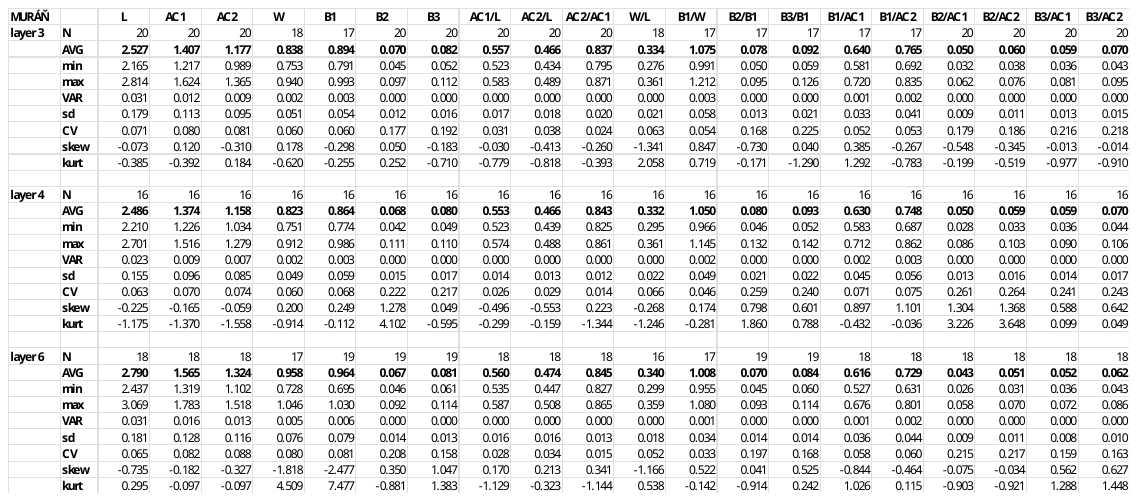

**Bišilu**

*Site characteristics:*
Czechia, 49°56'N 14°7'E; 290 m a.s.l.

Bišilu is a continuous stratigraphic section at the buried entrance of a cave in the central part of the Bohemian karst, near the village of Tetín. The section contains a greatly diversified glacial fauna in base layers 5–7 and a faunal sequence to the middle Holocene in upper layers. The radiocarbon data shown mean cal.14C ages from 25511 to 35535 in layer 6 and 12000 in layer 4. The site was excavated by Horáček et al. in 2014–2015.

*Stenocranius anglicus* patterns:

Within **Bišilu**, a gradual decrease in morphotype diversity is observed from Layer 7 to Layer 5, as indicated by Shannon indices. This trend suggests temporal variation in morphotypes, possibly driven by environmental or demographic changes. The progressive diversification within this locality might reflect microevolutionary differentiation or population structuring in response to shifting climatic conditions. Despite this decrease, the diversity indices at Bišilu indicate a moderate level of morphological variability (Fig. A8). Morphometric analyses reveal that width and length remain relatively stable, showing only slight decreases throughout the sedimentary layers (Table 8). These findings suggest a continuous and gradual morphological transformation in the narrow-headed vole population at Bišilu from the Late Pleistocene into the early Holocene. Geometric morphometric analyses further support this pattern. Bišilu exhibits moderate Procrustes and Mahalanobis distances, consistent with Balcarka, indicating substantial morphological deviation from the dataset mean. These distances, along with the temporal changes in diversity and size, reinforce the notion that Bišilu experienced dynamic evolutionary processes, potentially in response to environmental pressures leading up to the species' extinction at the locality.

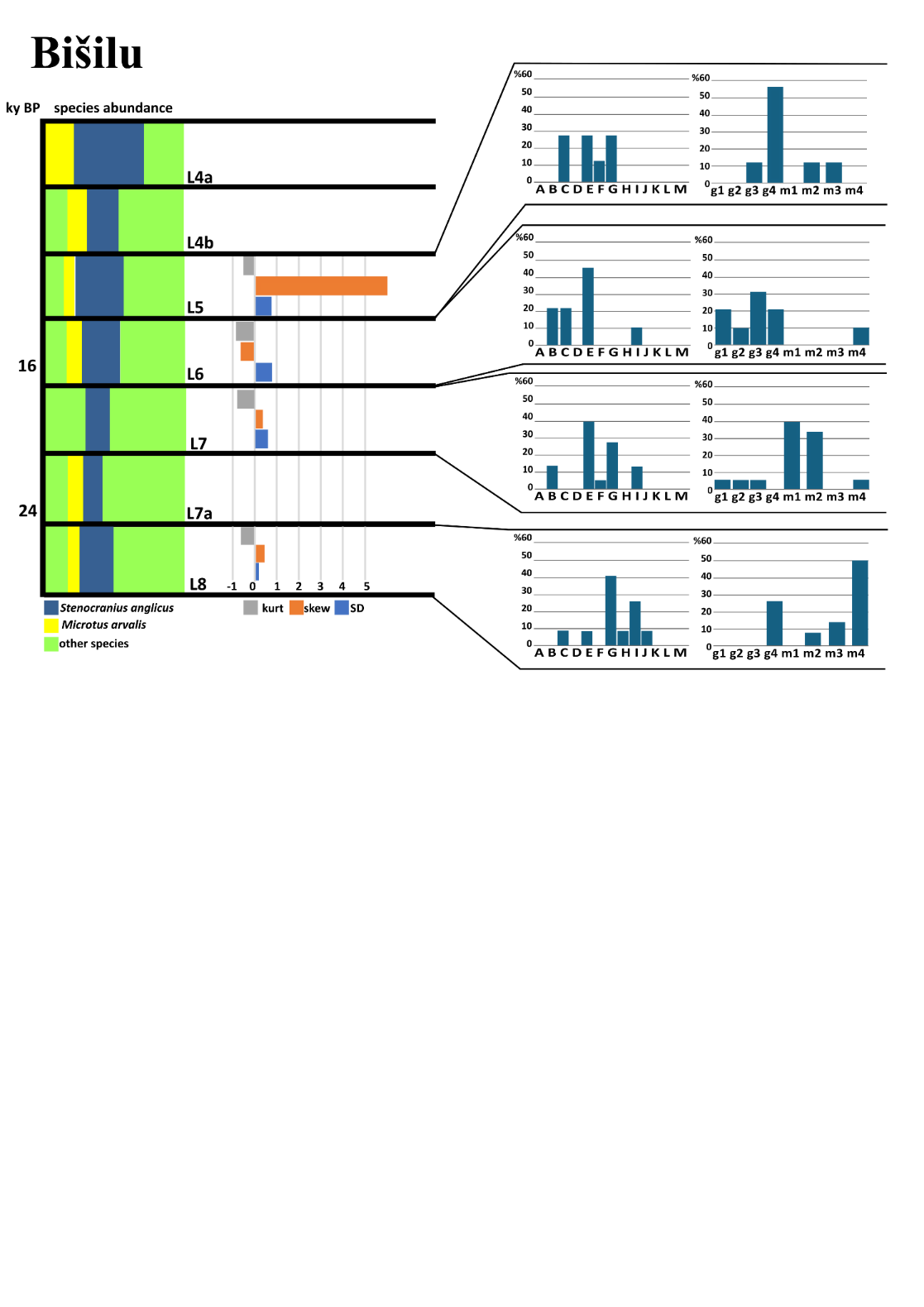

Figure A8. Characteristics of the narrow-headed voles at the locality Bišilu.
Species distribution, skewness, and kurtosis of m1 length and morphotype distribution (per Nadachowski 1982, BRA4 angle–based) of individual layers (L) at particular sedimentary sequences of locality Bišilu.

Table 8. Metric characteristics of individual layers in the population of Bišilu.
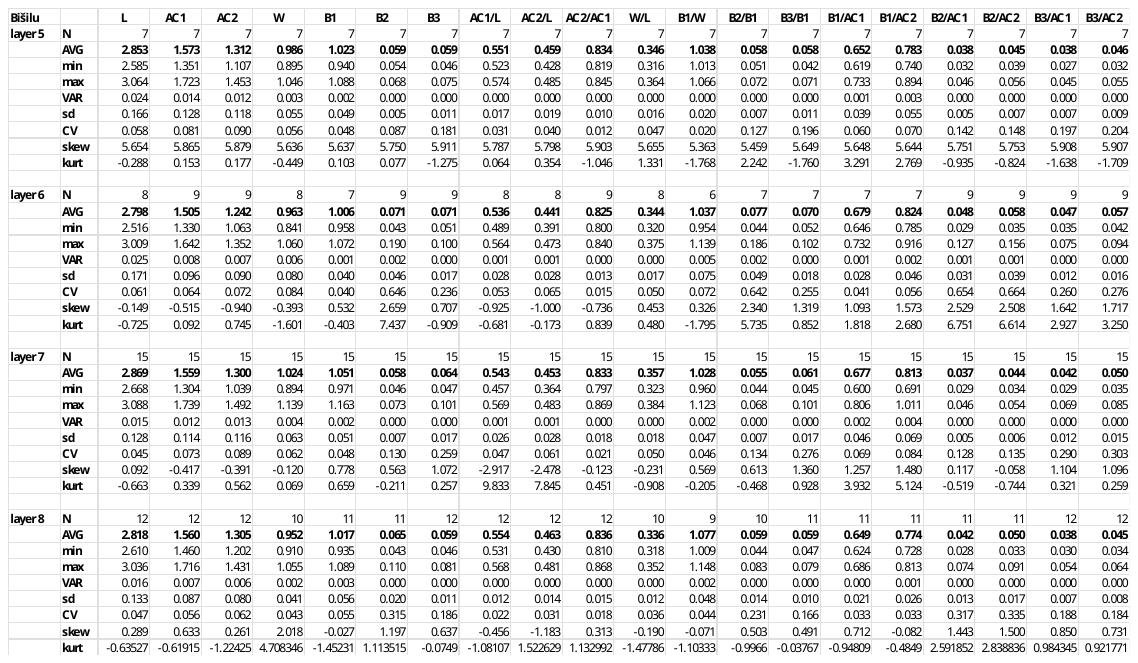

**Býčí Skála**

*Site characteristics:*
Moravian karst, 49°18'N 16°41'E; 316 m a.s.l.

Býčí Skála Cave, located in the western part of the Moravian Karst (Křtinský valley) near the town of Adamov, is famous for exceptional archaeological records of the Hallstatt period. Yet, the site reported here is situated in a side chamber near the cave entrance, infilled with a 10m thick talus deposit alternating 20 scree and soil colluvial layers. During the excavations by Horáček et al., which took place between 2007 and 2013, over 3 tons of fine-grained sediment were washed, yielding approximately 100,000 small mammal remains representing 4 525 MNI of 52 mammalian species, including 20 bat species. The studied sequence spans the period 12.5–8.4 ka cal BP (based on multiple dating of all layers). The ~3,500-year glacial–Holocene faunal transition is preserved here at particularly high resolution (ca. 100–200 years per layer). It provides a top record of vertebrate community turnover, unparalleled in European paleozoology (Horáček et al. 2014, Horáček 2015, Horáček and Sázelová 2017).

*Stenocranius anglicus* patterns:

**Býčí Skála** exhibited high morphological diversity, particularly around ~11.7–12.1ky BP, coinciding with early Holocene warming and potential niche expansion. This period likely represents one of the final phases of morphological complexity in the region, with gregaloid-microtid morphs still present. Despite moderate classification success (66.5%) in CVA, Procrustes and Mahalanobis distances indicate significant morphological divergence, possibly linked to adaptations before extinction (Appendix II). Morphotype diversity declined in younger layers, suggesting a transition toward species disappearance. Changes in metric characteristics across stratigraphic layers reveal a trend of increasing variation toward the end of the LGM, followed by a stabilization or slight reduction in variability in the early Holocene. Standard deviation trends suggest greater morphological plasticity during the late glacial, which decreased post–12ky BP. Skewness values fluctuate, with certain layers indicating shifts in trait distributions, potentially reflecting selective pressures or environmental filtering. Specimens of Layer 9a (~12.1ky BP) exhibit very high diversity (H′ = 1.845), possibly representing one of the final cold-adapted populations before Holocene warming, and reflect a mix of ancestral morphologies persisting alongside early Holocene adaptations. At Layer 8c (~12ky BP), the diversity remains high but slightly lower than in Layer 9a. Metric characteristics show a stabilization trend, with some reduction in variability compared to older layers. Skewness suggests potential shifts in trait distributions, possibly due to selective pressures during the transition. At Layer 8b (~11.9ky BP), morphological diversity remains notable, but a gradual decline begins. The decrease in standard deviation suggests reduced plasticity, potentially due to populations adapting to new environmental conditions or facing ecological pressures. Layer 8a (~11.7ky BP, early Holocene warming) shows moderate to high diversity, aligning with a phase of niche expansion. Morphological variation stabilizes, possibly reflecting the last remnants of complex morphotypes before a decline in variability (Fig. A9). The transition suggests species may have experienced temporary stability before final disappearance. Specimens from layer 7b (~11.1ky BP) exhibit the lowest diversity among the layers analyzed, indicating the species' decline. Morphological data suggest a reduction in variation, likely corresponding with environmental changes and the loss of cold-adapted traits. The increasing morphological homogeneity hints at a population in the final extinction phase. These findings support the hypothesis that the Býčí skála population retained substantial morphological complexity during the late glacial. Still, as the Holocene progressed, diversity and plasticity declined, leading to eventual extinction.

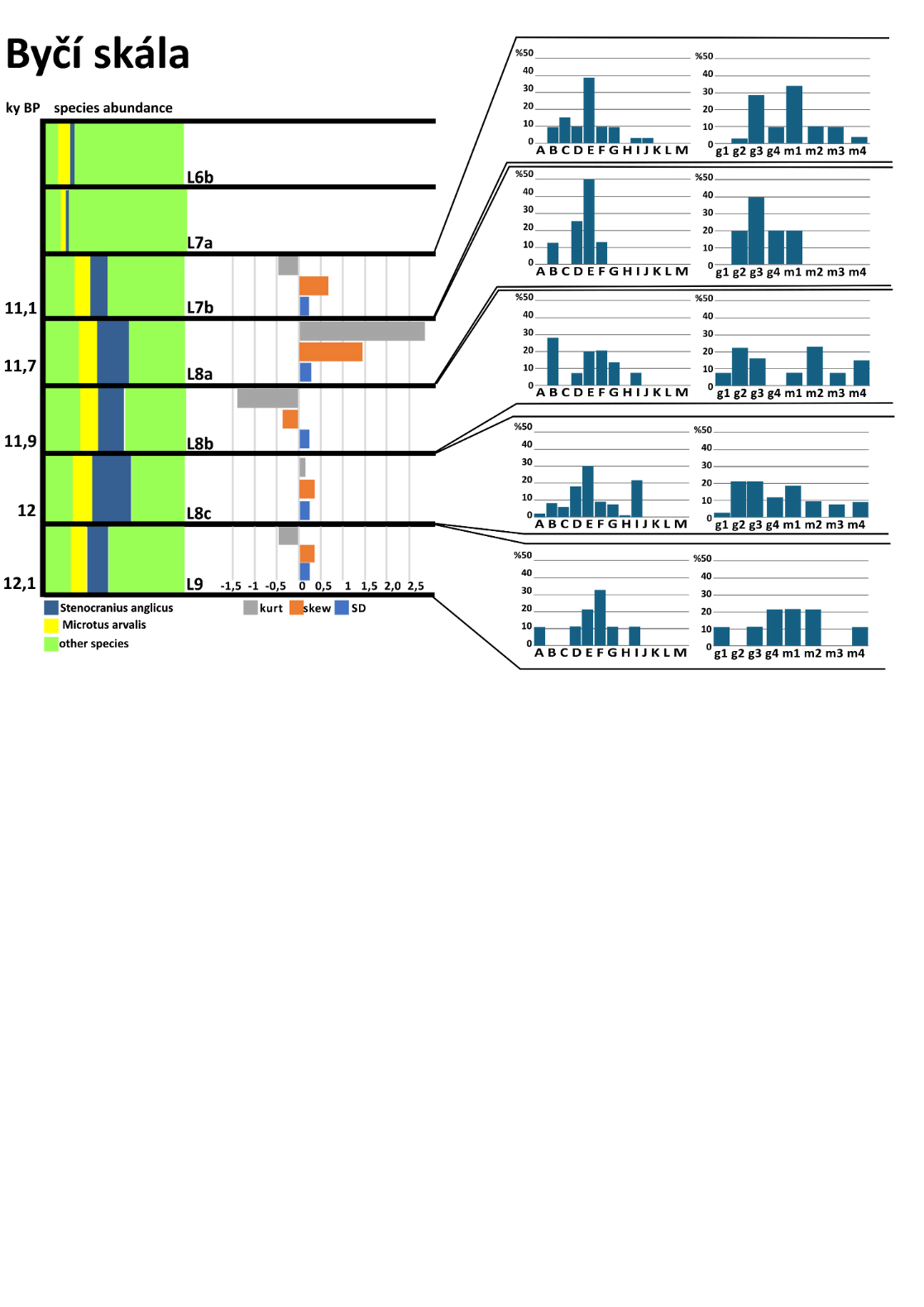

Figure A9. Characteristics of the narrow-headed voles at the locality Byčí skála.
Species distribution, skewness, and kurtosis of m1 length and morphotype distribution (per Nadachowski 1982, BRA4 angle-based) of individual layers (L) at particular sedimentary sequences of locality Býčí Skála.

Table 9. Metric characteristics of individual layers in the population of Byčí skála.

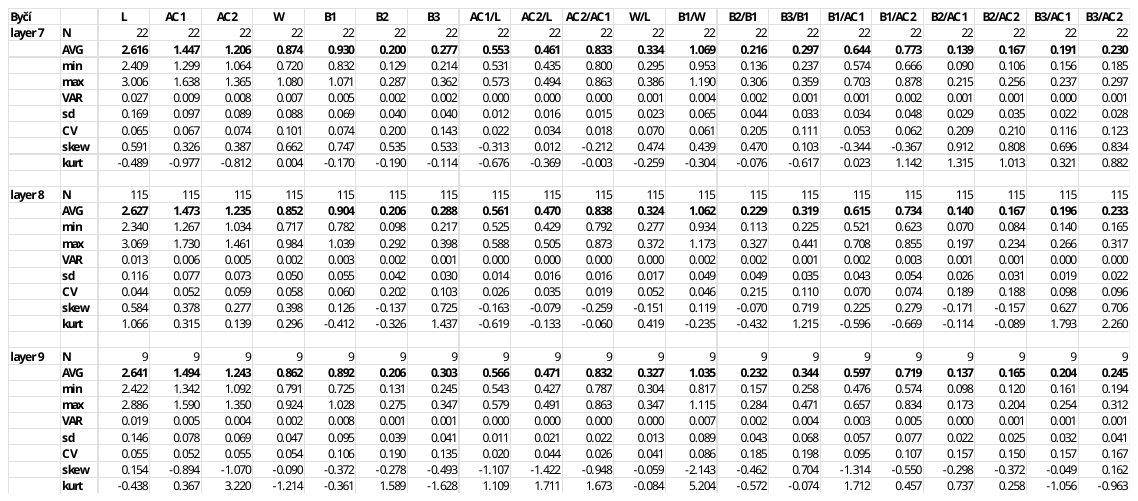

**Barová**

*Site characteristics:*
Moravian karst, 49°18'N 16°41'E; 345 m a.s.l.

Barová Cave is located adjacent to Býčí Skála in the central Moravian Karst. The main fossil-bearing layers (A15–A11) date to the Last Glacial. The cave was first made accessible in 1947, with early excavations carried out by Sobol (1948, 1949). Rich fossil material was recovered from three trenches (A, B, C), situated at the cave entrance, which covers a timespan from the Last Glacial through the Holocene to the Recent (Seitl et al. 1986; Horáček et al. 2002). The present analysis focuses on *Microtus* remains from layers A11–A15 and B1–B2, corresponding to the Last Glacial period (~40,000–10,300 BP) (Horáček et al. 2002).
 *Stenocranius anglicus* patterns:

Locality **Barová** exhibits high morphological variability, suggesting significant morphotype heterogeneity. Notably, morphotype diversity was highest in the older layers but declined in a more ambiguous layer, suggesting a transition from complex morphotype dominance before the Holocene to a simplification of forms in later periods. The site shares a substantial morphological overlap with Dzeravá, as evidenced by frequent misclassification in CVA. The stratigraphic sequence at the Barová section spans from the oldest layer A15 (~50ky BP) through 50-14ky BP (A14, 13) to the youngest A12, 11 (14-10ky BP), revealing a subtle but consistent pattern of morphological change over time. Molar length shows slight fluctuations, increasing from 2.574 mm around 50ky BP to 2.661 mm during 50-14ky BP, then decreasing to 2.627 mm in the youngest layers, indicating minor size variability without a clear directional trend. Anteroconid dimensions (AC1, AC2) remain stable, suggesting no significant shifts in anterior proportions. However, the posterior and buccal structures exhibit a subtle expansion trend. Ratios such as B1/AC1 increase from 0.614 (~50ky BP) to 0.634 (50-14ky BP) before slightly decreasing to 0.622 (14-10ky BP), while B3/AC1 rises from 0.205 in the oldest layer to 0.200 in the youngest. Additionally, the B2/B1 and B3/B1 ratios show slight increases over time, indicating a proportional enlargement of the distal molar regions. Despite these modifications, width (W) remains stable at ~0.84 mm, maintaining a W/L ratio of approximately 0.32 across all layers (Table 10). Overall, while absolute size variability at Barová is minimal, the observed posterior expansion and increasing complexity of buccal structures suggest gradual morphological rearrangements.

Table 10. Metric characteristics of individual layers in population of Barová.

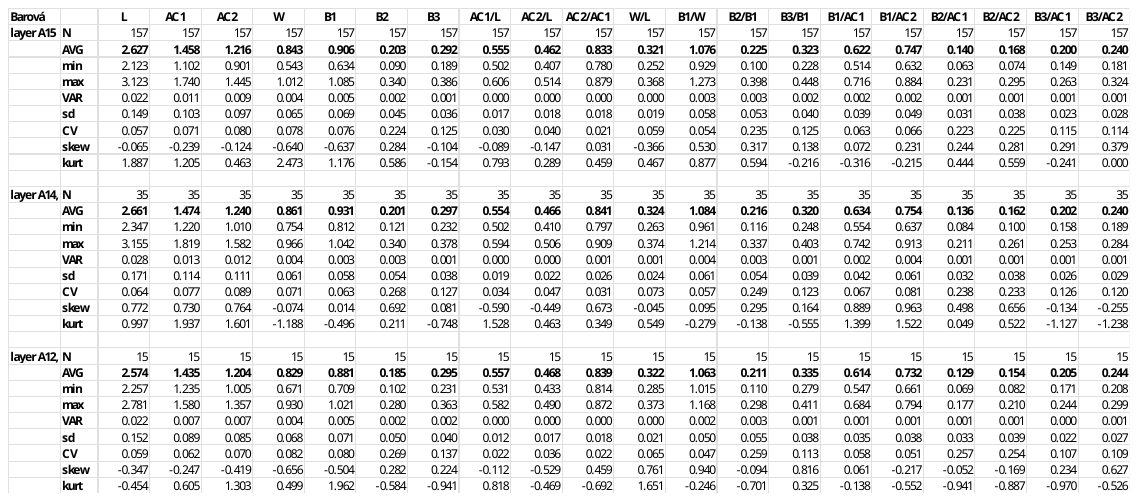

**Dzeravá skala**

*Site characteristics:*
Slovakia, 48°29'N 17°19'E; 450 m a.s.l.

Dzeravá skala cave, situated in the Mokrá dolina Valley in the Malé Karpaty Mountains (Plavecký Mikuláš), is a well-known archaeological, anthropological, and paleontological site in Western Slovakia. The site is located in a short but deep-cut karstic valley on the western slopes of these mountains, facing the Morava River plain. The cave entrance is 18 m wide, 22 m long, and 10 m high. Excavations in 2002–2003, which yielded the samples examined in this paper, were led by Kaminská, Kozlowski, and Svoboda. It unveiled complex stratigraphy combining in situ sediments, loess, clays, paleosols, and roof-collapse clasts. The sedimentary series excavated in Dzerava skala cave (Male Karpaty Mts., SW Slovakia) provided a sequence of rich assemblages of small vertebrates, particularly mammals (MNI = 2013, 52 spp.), which exhibited a repeated fluctuation in percentage of core glacial elements and species diversity corresponding to the climatic development from early middle Weichselian to the beginning of the Holocene (MIS 4 to MIS 1). Together with dominating elements of glacial fauna (*S.anglicus* in particular), two species demanding woodland habitat, *Clethrionomys glareolus* and *Sorex araneus*, appeared continuously throughout the whole section, which demonstrated their refugial survival in the W Carpathians throughout the most severe part of the Weichselian glacial period (Horáček 2006, 2015).

*Stenocranius anglicus* patterns:

**The Dzeravá skala** sequence in Slovakia demonstrates remarkable morphological stability over more than 40 000 years, with consistently high morphotype diversity across most stratigraphic layers. This sustained diversity suggests long-term population persistence, preserving multiple morphotypes through environmental fluctuations until their final disappearance in the Holocene, spanning 53 350 cal. BP (Layer 11) to 12ky cal. BP (Layer 1), molar proportions remain highly conserved, with tooth length (L) fluctuating narrowly between ~2.6–2.7 mm. Anteroconid ratios and width proportions remain stable, indicating no significant anterior elongation or reduction. However, a gradual increase in B2 and B3 from older to younger layers suggests a subtle posterior expansion over time. Layer-by-layer analysis reveals a consistent molar morphology, with only slight variations in size and proportions (Table 11). Morphometric analyses reinforce this stability, indicating localized adaptation or terminal differentiation. The combination of high diversity, morphological conservation, and subtle posterior expansion suggests a long-time continuity of a local population in Dzeravá skala, maintaining vole populations through changing climatic conditions well into the late Pleistocene.

Table 11. Metric characteristics of individual layers in the population of Dzeravá skala
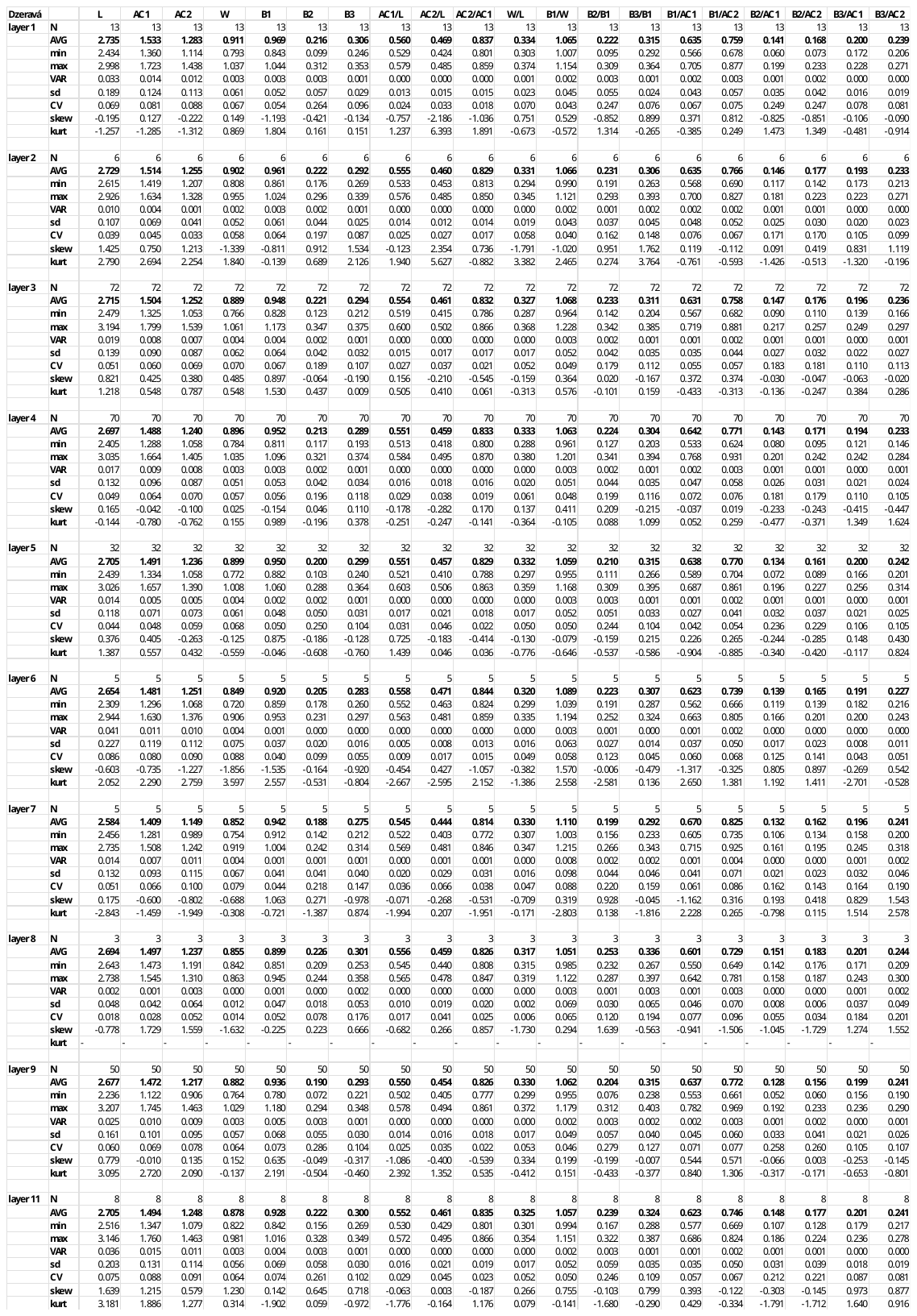

**Holštejnská Cave**

*Site characteristics:*
Moravian karst, 48°29'N 17°19'E; 470 m a.s.l.

Holštejnská Cave, located near the village of Holštejn in the Eastern margin of the Moravian Karst, is a karst cavity nearly entirely filled with late Pleistocene sediments. Paleozoological analyses from the entrance deposits were reported by Horáček and Ložek (1988), who emphasized the stratigraphic importance of Holštejnská for reconstructing faunal changes along Vistulian/ Holocene transition. Apart from the stratigraphic section at the entrance of the cave, further material was obtained from intracave loess sediments, supposedly of MIS 3 age (labelled here as Holštejnská intracave).

*Stenocranius anglicus* patterns:

**Holštejnská Cave** sequence in the Czech Republic exhibits intermediate levels of size and shape variation, with a gradual trend of morphological simplification and decreasing diversity leading up to local extinction. Moderate divergence is consistent with other late–surviving populations. Morphometric data from individual layers reveal a clear pattern of size reduction and morphological homogenization over time (Fig. A10). The oldest layer (8) contained the largest and most robust molars, with an average length of 2.839 mm and a width of 0.958 mm. Proportional traits indicate broad, well–developed molars typical of a stable, diverse population. Variation was moderate, and trait distributions were balanced, suggesting a well–established population structure. By layer 6, molars exhibited a slight decrease in size (L = 2.748 mm, W = 0.936 mm), while proportional traits remained stable. However, variation in buccal traits increased, with skewness and kurtosis values indicating growing asymmetry and peakedness, possibly reflecting emerging population stress or environmental fluctuations during the Late Glacial (Table 12). In layer 5 (~13.4ky cal BP), the trend of size reduction continued (L = 2.743 mm, W = 0.957 mm), though it varied slightly. Morphological traits stabilized (AC1/L = 0.560, W/L = 0.347), but asymmetry in buccal proportions increased, suggesting heightened trait concentration and reduced variability. By the youngest layer (4) (~5.6ky cal BP), molars were notably small and uniform (L = 2.673 mm, W = 0.901 mm, AC1/L = 0.561). Variability was low and trait distributions were tightly constrained, with moderate skewness. Towards the extinction of the narrow-headed vole at the locality Holštejnská, the length of the tooth (L) as well as the length of the anteroconid (AC1) and anteroconid complex (AC2) decreases with time (R^2^ = 0,9107; R^2^ = 0,8598; R^2^ = 0,8337, respectively). These patterns strongly indicate a final phase of morphological simplification and declining diversity, likely associated with population stress and eventual local extinction under Holocene ecological pressures.

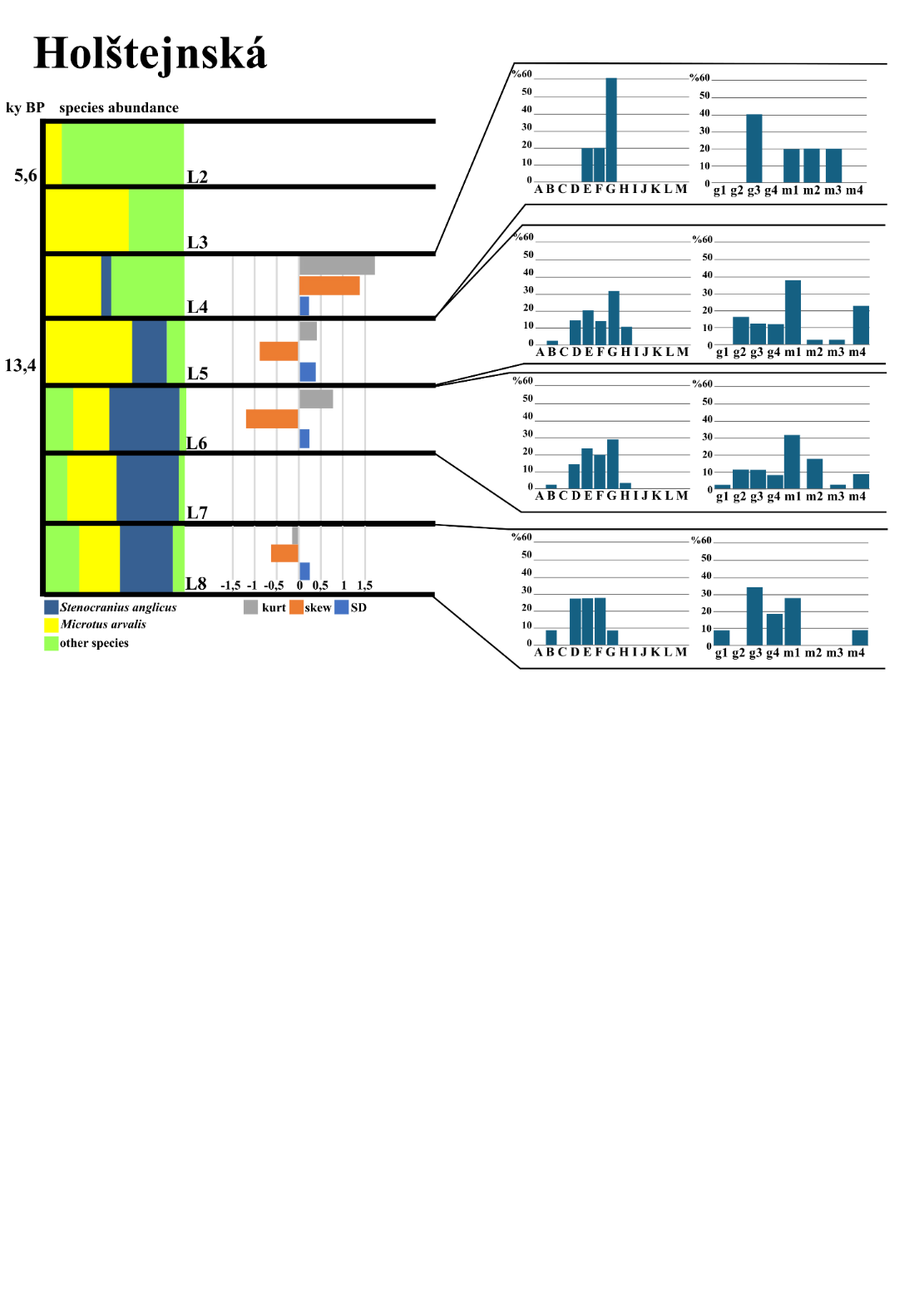

Figure A10. Characteristics of the narrow-headed voles at the locality Holštejnská.
Species distribution, skewness, and kurtosis of m1 length and morphotype distribution (per Nadachowski 1982, BRA4 angle-based) of individual layers (L) at particular sedimentary sequences of locality Holštejnská.

Table 12. Metric characteristics of individual layers in the population of Holštejnská.

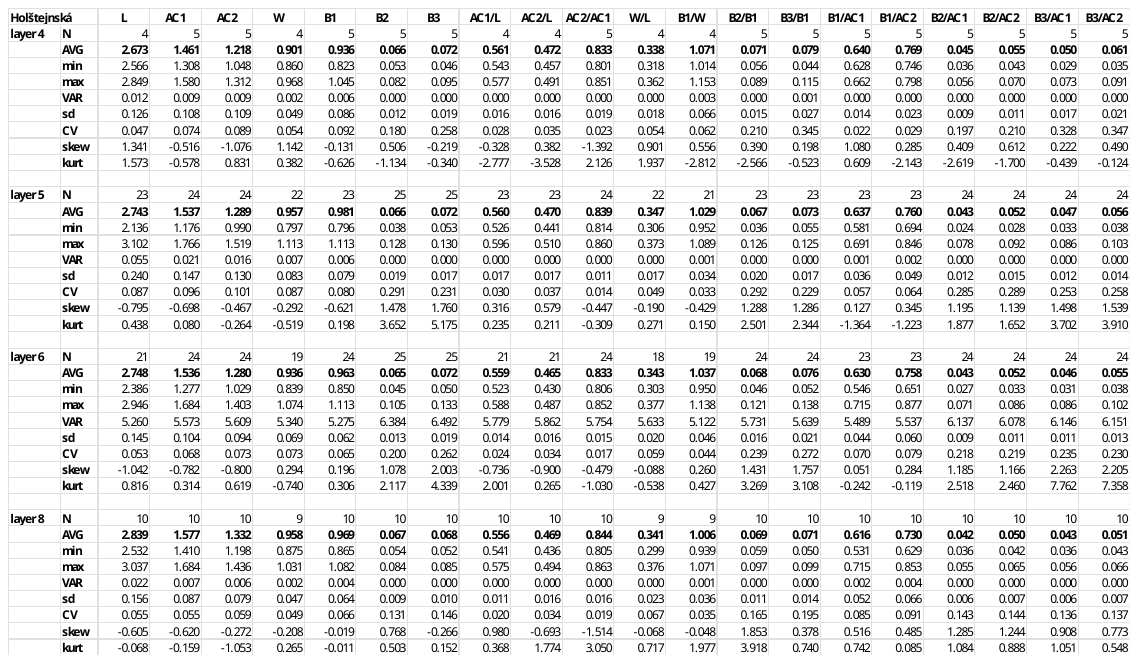

**Maštalná**

*Site characteristics:*
Slovakia, 48°35'N 20°27'E; 600 m a.s.l.

Maštalná is located in southeastern Slovakia, in the Slovakian karst near Slavec, at the uppermost wall of the high slopes of Plešivecká plateau. A 4.5 m deep stratigraphical section in a ca 15 m thick debris talus at the cave entrance, excavated in 1982–1984 by V. Ložek and I. Horáček, revealed a continuous sedimentary sequence of 17 layers covering a time span from the Late Vistualian to the Recent. It provided a quite rich faunal record, both of molluscs and small vertebrates, the latter being represented by MNI 1851 of 55 spp. For more details, see Horáček and Ložek (1988), Horáček (2015), or Horáček and Sázelová (2017).

*Stenocranius anglicus* patterns:

The population of **Maštalná** has the smallest average molars among all studied localities. Procrustes and Mahalanobis distance suggest minimal morphological divergence and shape convergence toward the dataset mean, potentially reflecting stabilizing selection before extinction. Morphotype diversity trends in Maštalná reveal a sharp decline over time. While layers 13–15 (~15ky cal BP) maintained relatively high diversity, younger layers (10b, 10) exhibited a drastic reduction in diversity. The extreme diversity loss in layer 10 suggests a small, isolated population with restricted morphotype spectra at the edge of extinction (Fig. A11). Morphological data from Maštalná indicate a pattern of progressive size reduction and trait simplification over time. Molars from layer 15 (16ky cal BP) were slightly smaller than in layers 13 and 14 (L = 2.735 mm, W = 0.946) but exhibited high variation in basin traits. Skewness and kurtosis were extreme, indicating early population expansion with diverse morphotypes. Molars from layer 14 were slightly larger (L = 2.752 mm, W = 0.927 mm) and exhibited morphological consistency. Variability was lower than in layer 13, and skewness and kurtosis remained moderate, suggesting population stabilization. At layer 13, a modest increase in molar size was observed (L = 2.745 mm, W = 0.929 mm), with robust traits and high variation in basal features. Extreme skewness and very high kurtosis in basin traits suggest strong asymmetry and trait peaking, likely reflecting environmental instability or morphotype divergence. Molars of layer 12 showed a slight size reduction (L = 2.671 mm, W = 0.906 mm), but proportions remained stable. Variability remained moderate. A notable increase in molar size occurred in layer 11 (L = 2.771 mm, W = 0.857 mm), possibly reflecting a brief population resurgence. Proportional traits suggested elongated molars with less breadth, while variation remained moderate. Trait distributions were skewed and peaked, suggesting selective pressures. Molars of layer 10 (11.4ky cal BP) were smaller (L = 2.601 mm, W = 0.919 mm), with proportions remaining consistent. Trait variation was moderate, and skewness suggested mild asymmetry in basal measurements. This layer likely represents a declining but still viable population. At the youngest layer, 10b (10.4ky cal BP in 10), molars reached their smallest and most gracile form (L = 2.308 mm, W = 0.775 mm) (Table 13). High variability in several traits and skewness/kurtosis values indicate asymmetric distributions with extreme outliers. This suggests a terminal morphological phase before extinction.

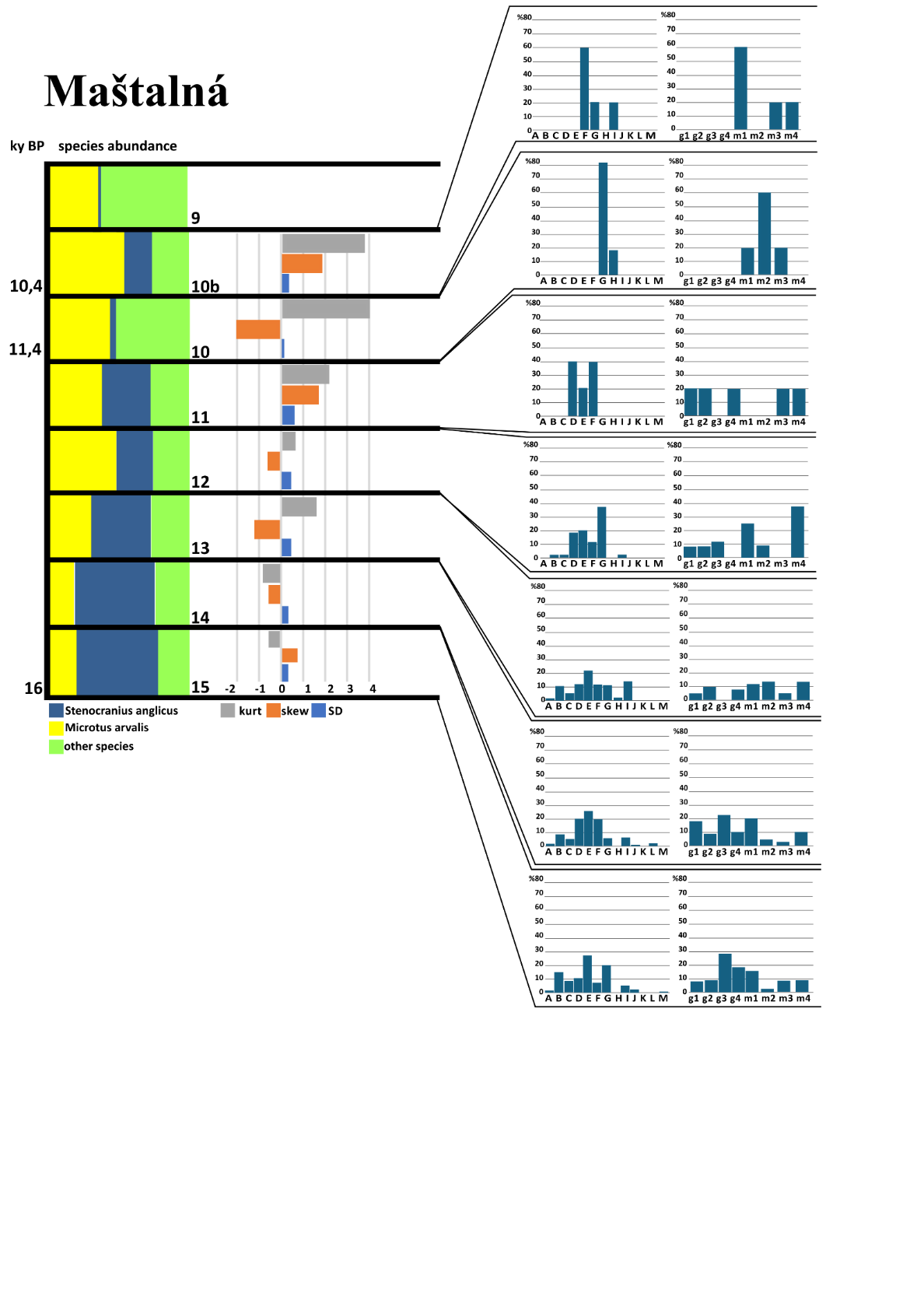

Figure A11. Characteristics of the narrow-headed voles at the locality Maštalná.
Species distribution, skewness*,* and kurtosis of m1 length and morphotype distribution (per Nadachowski 1982, BRA4 angle-based) of individual layers (L) at particular sedimentary sequences of locality Maštalná.

Table 13. Metric characteristics of individual layers in the population of Maštalná.

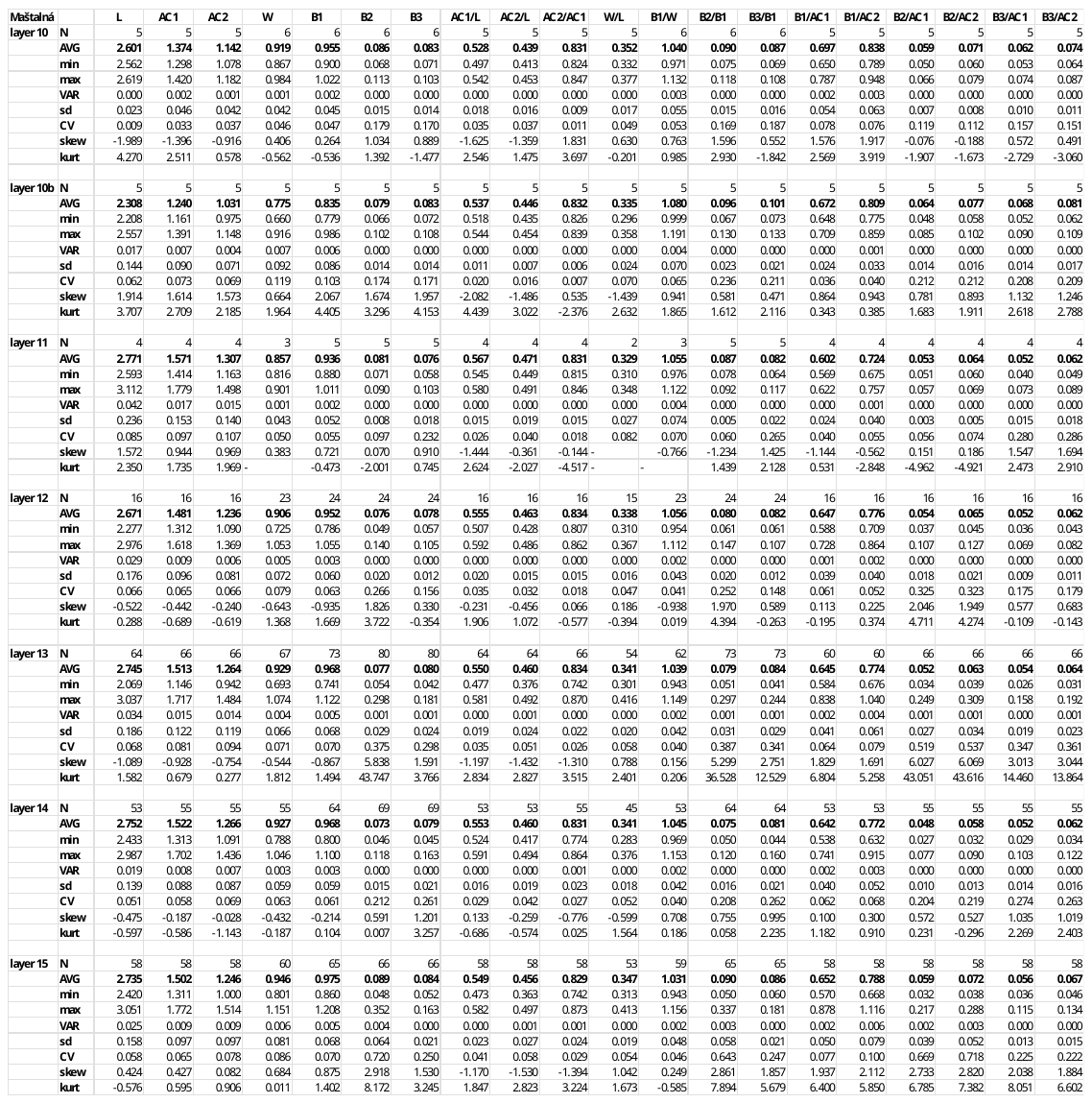

**Srnčí**

*Site characteristics:*
Moravian karst, 49°22'N 16°44'E; 470 m a.s.l.

Srnčí is a small tunnel cave located in the upper part of the Macocha slope, east of Macocha Abyss, in the Moravian Karst. Historical excavations brought numerous records of large mammals such as *Ursus spelaeus*, *Panthera spelaea*, and *Coelodonta antiquitatis.* The material surveyed in this study was excavated from the intact cave entrance deposits by V. Ložek, J. Vašátko, and I. Horáček in 1978. Small vertebrates (MNI 7122, 44 spp.) were presented in all 13 layers, with *S. anglicus* in 10 lower layers of the section.

Stenocranius anglicus patterns:

The population of Srnčí Cave represents one of the last occurrences of the species in the area. The site is characterized by low and fluctuating diversity, with some of the lowest post–LGM diversity values. These values suggest localized population bottlenecks or environmental pressures that may have contributed to the species’ decline. Progressive morphotype homogenization as the population approached extinction is apparent across all analyzed layers. Morphometric analysis further supports the idea of a declining population, exhibiting minimal morphological divergence from the dataset mean. Canonical Variate Analysis (CVA) also revealed substantial misclassification (40.9%), reinforcing the idea that the Srnčí population closely resembled the average morphological form across all sites (Appendix II). Measurements from layers 8, 7, 5, and 4 spanning the Late Glacial to early Holocene indicate a relatively stable molar size. The average molar length and width decreased from layer 8 to layer 4 (L = 2.673 mm, W = 0.861 mm; L = 2.501 mm, W = 0.845 mm, respectively). However, width remained remarkably consistent across layers (~0.845–0.864 mm), and proportional traits such as AC1/L and AC2/L exhibited stability, suggesting limited morphological change over time. Despite overall stability, layer 5 showed greater variation in traits such as B2 and B3, as well as in their proportional relationships (e.g., B2/B1, B3/B1), suggesting episodes of greater individual variability. Additionally, skewness and kurtosis values for B2 and B3 in layer 5 indicate asymmetry and the presence of outliers, potentially reflecting periodic environmental shifts or small-scale population fluctuations (Table 14). However, these fluctuations did not lead to major morphological shifts, and overall, the population retained a consistent molar morphology across all analyzed layers. The morphological stasis characterized by reduced morphotype variation observed at Srnčí persisted prior to its final decline, until its last appearance in layer 4, which exhibited a pattern not recorded at other sites. The combination of low diversity, shape convergence, and lack of significant differentiation from the dataset mean implies that this population was not undergoing substantial evolutionary change but instead maintained a relatively stable form until extinction. This may indicate stabilizing selection, in which reduced genetic and morphological variation limited adaptive potential, ultimately contributing to the species’ disappearance from the region.

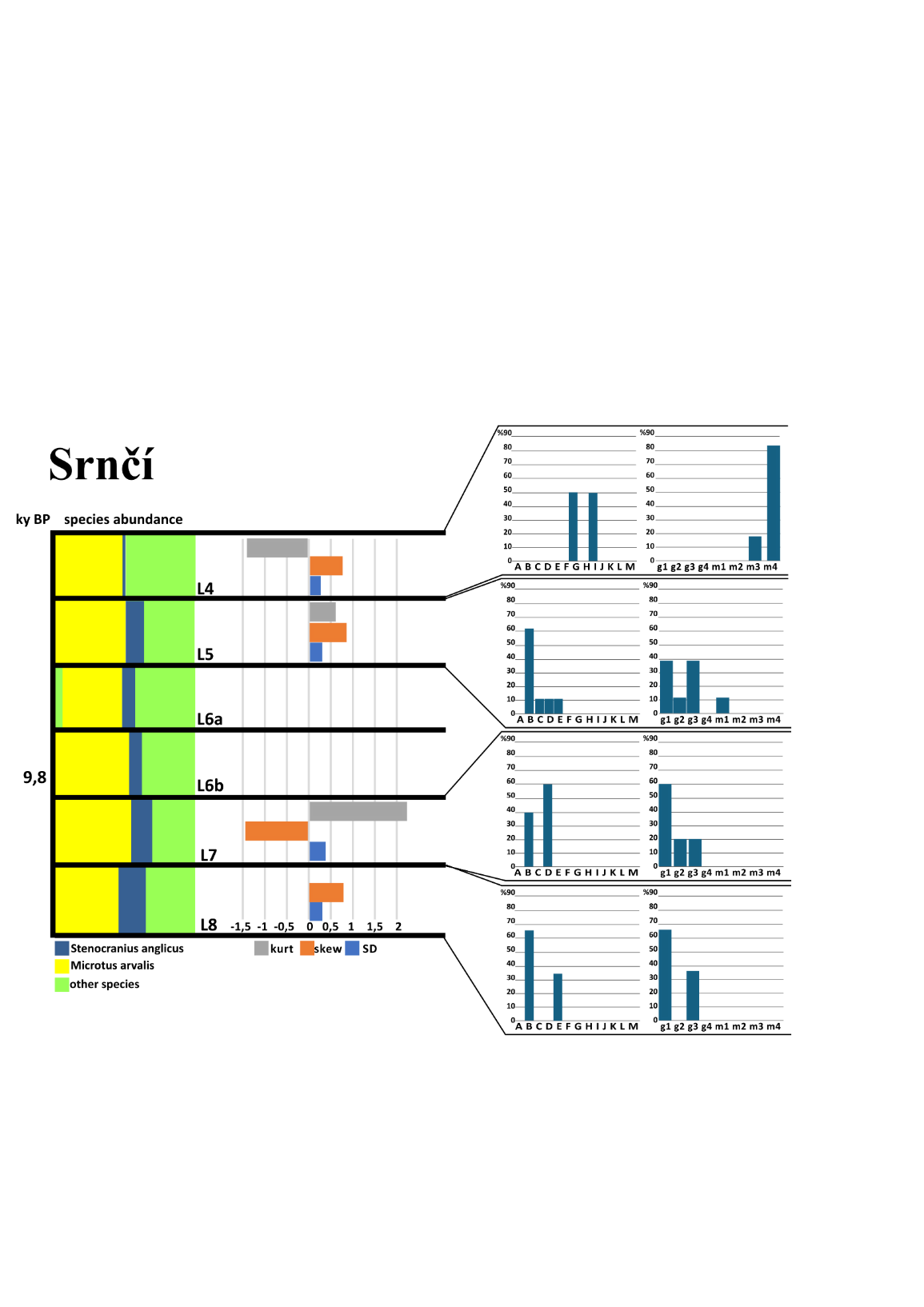

Figure A12. Characteristics of the narrow-headed voles at the locality Srnčí.
Species distribution, skewness, and kurtosis of m1 length and morphotype distribution (per Nadachowski 1982, BRA4 angle-based) of individual layers (L) at particular sedimentary sequences of locality Srnčí.

Table 14. Metric characteristics of individual layers in the population of Srnčí.

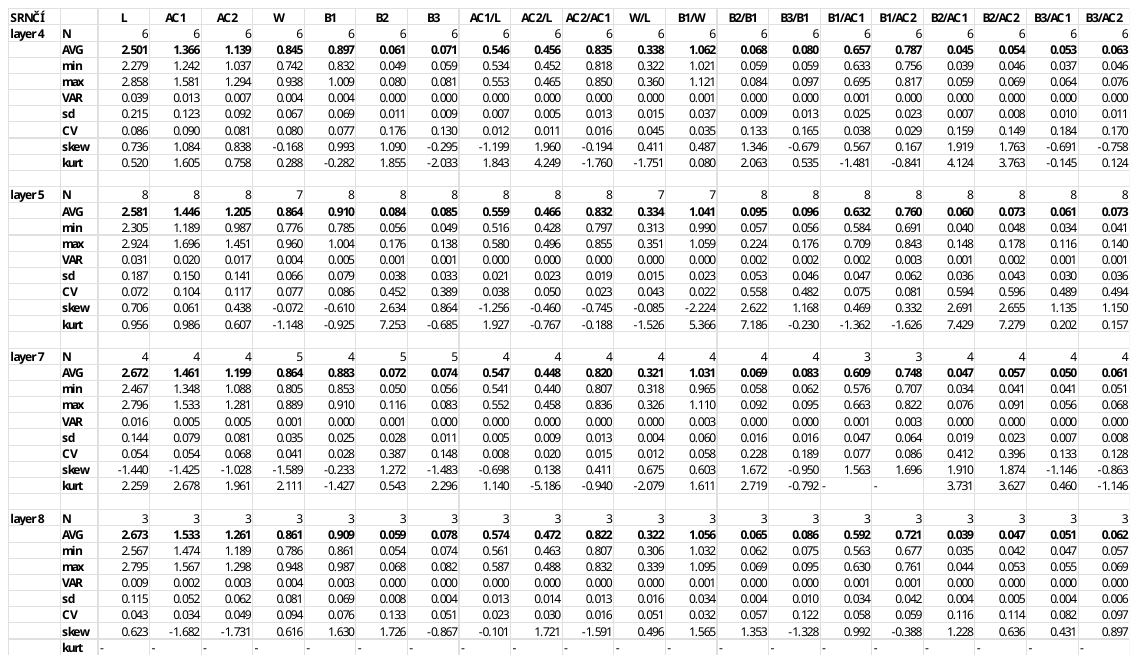
